# O-GlcNAcylation-Ubiquitin Crosstalk of METTL1 Drives m7G Epitranscriptomic Collapse and Lipid Metabolic Reprogramming in Diabetic Cardiomyopathy

**DOI:** 10.1101/2025.07.07.663452

**Authors:** Xin Gu, Haoyu Meng, Jiabao Liu, Yan Liang, Baihong Li, Fen Wang, Qian Liu, Zhixuan Zhang, Jialong Liang, Xiaodong Zhang, Jialong Sun, Jianwei Li, Fenqi Liu, Wang Xiao, Guangyi Huang, Tianya Gu, Su Peng, Xin Huang, Ruijuan Zhuang, Jun Zhang, Yafei Li, Jiabao Ye, Lijie Lu, Xiaoyan Wang, Fenglai Yuan, Junbo Ge, Yingqiang Du

## Abstract

**Background:** Metabolic remodelling and memory in cardiomyocytes is a pivotal mechanism underlying cardiomyopathy pathogenesis. Clinical observations demonstrate persistent progression of hyperglycaemia-induced multiorgan damage following blood glucose stabilization, which is predominantly mediated through epigenetic regulation. While prior studies have identified epigenetic contributions to hyperglycaemic myocardial injury, the involvement of RNA methylation-particularly N7-methylguanosine (m7G) modification-in this regulatory network remains undefined.

**Methods:** Clinical specimens were collected from diabetic patients, and cardiomyocyte-specific METTL1 and ob/ob knockout murine models were established in parallel. Multiomic profiling (proteomics, glycoproteomics, ubiquitinomics, m7G-MeRIP sequencing, and metabolomics) was systematically conducted. The molecular mechanisms governing METTL1 regulation via O-GlcNAcylation and ubiquitination were elucidated through integrated *in vitro* and *in vivo* assays. A DUB siRNA library and computational strategies combining molecular docking with molecular dynamics simulations were employed for screening drugs targeting METTL1 O-GlcNAcylation, followed by *in vivo* therapeutic validation.

**Results:** Comparative analysis of diabetic murine and human samples revealed strong METTL1 downregulation in cardiomyopathy contexts. Tamoxifen-inducible METTL1 knockout mice presented exacerbated diabetic cardiomyopathy phenotypes, confirming its cardioprotective function. Multiomic integration demonstrated that METTL1-mediated m7G modification critically regulates cardiomyocyte fatty acid metabolism. Mechanistically, hyperglycaemia was found to induce O-GlcNAcylation at the METTL1-T268 residue, suppressing m7G methyltransferase activity by 38% (p < 0.01). Subsequent investigations revealed that USP5 deubiquitinase activity is impaired under hyperglycaemic conditions, leading to accelerated METTL1 degradation. Notably, administration of the first-in-class small drug HIT106265621 significantly attenuated cardiomyopathy-associated pathological alterations in ob/ob mice *in vivo*.

**Conclusion:** Hyperglycaemia promotes METTL1 O-GlcNAcylation, which impedes USP5-mediated deubiquitination, consequently reducing cardiomyocyte METTL1 protein levels and m7G modification. METTL1 deficiency drives diabetic cardiomyopathy progression through fatty acid metabolic dysregulation, inflammatory activation, and myocardial hypertrophy. Pharmacological inhibition of the OGT-METTL1 interaction using HIT106265621 has therapeutic potential for metabolic cardiomyopathy intervention.

## Introduction

The global prevalence of diabetes has demonstrated a consistent upwards trajectory in recent years, resulting in substantial mortality attributable to chronic complications associated with prolonged disease duration^1,2^. The Diabetes Atlas (2021) reported that approximately 10.5% of the adult population worldwide is affected by diabetes mellitus, with epidemiological projections indicating a 46% increase in the number of affected individuals by 2045^3,4^. Conventional therapeutic approaches for metabolic disorders are strongly limited by the metabolic memory phenomenon. Even with successful glycemic regulation via lifestyle modifications and insulin therapy, many patients exhibit delayed-onset complications, indicating that transient hyperglycemia induces persistent cellular reprogramming through metabolic memory mechanisms, thereby sustaining tissue damage post-treatment^5,6^. The effects of diabetic metabolic memory are more profound than we thought. It uses epigenetic changes to create lasting harm in the heart^7,8^, kidneys^9^, retinopathy^10,11^, and other systems. Although the scientific community has operationalized the term “metabolic memory” to describe this clinical paradox, the molecular mechanisms underlying this process remain multifaceted and incompletely elucidated.

The mechanistic basis of metabolic memory is considerably complex, with epigenetic modifications being widely acknowledged as the predominant regulatory pathway driving this phenomenon in diabetic patients or other nutrients^12–15^. RNA methylation is a critical epigenetic regulatory mechanism that is hypothesized to critically affect metabolic memory programming^16–18^. The progressive evolution of genomic technologies has facilitated the identification and characterization of numerous RNA modifications, with over 150 distinct chemical alterations in RNA molecules being systematically catalogued and investigated by 2017. Contemporary research efforts have focused predominantly on m6A and m5C modifications^17^; however, the recent emergence of N7-methylguanosine (m7G) as a novel RNA modification subtype has attracted increasing scientific attention^19,20^. Notably, m6A modification has been implicated as a pivotal regulatory mechanism in the establishment of metabolic memory during diabetic nephropathy progression^21^. Prior investigations have further established associations between m6A RNA modification and multiple pathogenic factors contributing to diabetic cardiomyopathy (DCM) development, including but not limited to inflammatory cascades, fibrotic processes, and oxidative stress pathways^22–24^.

m7G, a prevalent post-transcriptional RNA modification, plays essential regulatory roles in diverse RNA processing events, metabolic pathways, and functional mechanisms^25,26^. The biochemical regulation of m7G modification is principally mediated through the coordinated interaction between methyltransferase-like 1 (METTL1) and WD repeat domain-containing protein 4 (WDR4)^19,27^. Substantial experimental evidence from multiple studies has demonstrated that the METTL1/WDR4 complex performs context-dependent oncogenic or tumour-suppressive functions across various malignancies-including head and neck carcinomas^28^, pulmonary neoplasms^29^, hepatic tumours^30^ and esophageal squamous cell carcinomas^31^-through the modulation of tRNA or miRNA m7G methylation status. Aberrant m7G was reported to alter B-cell responses in systemic autoimmunity^32^. Emerging research has further elucidated the involvement of METTL1-mediated m7G modification of mRNAs in diverse pathological conditions, including angiogenic regulation^33^, and neurological system disorders^34,35^. Previous investigations have substantiated the critical participation of METTL1 in the modulation of myocardial fibroblast activity and cardiac hypertrophic responses^36,37^.Additionally, METTL1 was reported to regulated ketogenesis through translational regulation and drives metabolic reprogramming in postnatal heart development^38,39^. Nevertheless, the functional contribution of m7G modification to metabolic memory dynamics and its pathophysiological role in DCM pathogenesis remain predominantly uncharacterized.

Post-translational protein modifications are essential regulatory mechanisms governing cellular protein functionality. Among these modifications, glycosylation processes have been identified as major contributors to hyperglycaemia-induced metabolic memory phenomena. Glycosylation can be systematically categorized into N-linked GlcNAcylation and O-linked GlcNAcylation subtypes^40,41^. The hexosamine biosynthetic pathway (HBP) generates uridine diphosphate N-acetylglucosamine (UDP-GlcNAc), which is enzymatically conjugated to serine or threonine residues via O-GlcNAc transferase (OGT) catalytic activity. UDP-GlcNAc additionally participates in asparagine residue modification through alternative HBP-derived intermediates, which exhibit biological activities distinct from those of canonical O-GlcNAcylation processes^42^. Dysregulated GlcNAcylation mechanisms have been aetiologically linked to numerous severe human pathologies, including cardiac failure^43,44^, pancreatic tumor^45^, and Alzheimer’s disease (AD)^46^. Specifically, increased levels of protein O-GlcNAcylation are found in both rodent and human hypertrophic hearts^42^. AMPK activation might improve cardiac hypertrophy predominantly by inhibiting O-GlcNAcylation via controlling GFAT phosphorylation^47^. Furthermore, aberrant O-GlcNAcylation patterns have been implicated in the pathogenesis of DCM^44^. While the regulatory importance of O-GlcNAcylation in various cardiac pathologies has been well documented, its potential involvement in cardiomyocyte metabolic memory formation and RNA modification processes during DCM development remains unclear.

In the present investigation, we identified significant downregulation of METTL1 protein expression in DCM through comprehensive analysis of specimens obtained from multiple diabetic murine models and clinical patients with confirmed DCM. Furthermore, we experimentally validated the essential cardioprotective role of METTL1 in DCM pathophysiology using tamoxifen-inducible cardiomyocyte-specific METTL1 knockout murine models (METTL1^CKO^ (METTL1^fl/fl^Myh6^CreERT^)). Our integrative multiomic sequencing analysis-encompassing proteomic profiling, metabolomic characterization, and m7G sequencing (m7G-seq)-revealed that METTL1-mediated m7G modification is a critical regulatory mechanism governing key enzymatic components involved in fatty acid metabolic remodelling within cardiomyocytes. This epigenetic modification mechanism was further demonstrated to maintain fatty acid metabolic homeostasis in cardiomyocytes under chronic hyperglycaemic conditions. Through synergistic integration of glycoproteomic and ubiquitinomic datasets, we observed that sustained hyperglycaemia induces O-GlcNAcylation at the threonine 268 (T268) residue of the METTL1 protein in cardiomyocytes, resulting in functional inhibition of its methyltransferase activity. Systematic screening employing a deubiquitinating enzyme (DUB) small interference library, complemented by mass spectrometric data analysis and *in vitro* functional validation, revealed that hyperglycaemic conditions critically regulate O-GlcNAcylation of ubiquitin-specific peptidase 5 (USP5), thereby inhibiting METTL1 deubiquitination processes and promoting its proteasomal degradation in DCM. Our experimental findings establish a novel association between RNA m7G modification and fatty acid metabolic remodelling in diabetic cardiomyocytes while elucidating the hyperglycaemia-induced dysfunction of m7G modification mechanisms through coordinated O-GlcNAcylation and ubiquitination regulatory pathways.

The present study aims to provide novel mechanistic insights into regulatory targets underlying metabolic memory phenomena in DCM while identifying potential therapeutic targets for the clinical management of this pathological condition.

## Materials and methods

All necessary Materials and Methods are described in the Supplementary Materials.

## Results

### METTL1 protein but not mRNA levels are decreased in diabetic cardiomyocytes

To elucidate the role of the METTL1 protein in cardiomyocytes, we performed an analysis using the BioGPS database, GTEx dataset, and Expression Atlas database (Fig. 1A and S1B and C). The analysis revealed significantly high expression of METTL1 in cardiac tissue, indicating a pivotal role of the METTL1 protein in heart disease. Additionally, Gene Ontology (GO) analysis of mice with DCM highlighted the involvement of RNA methylation and tRNA methylation in this process (Fig. 1B). Further investigation into the expression pattern of the METTL1 protein in DCM involved a comparison of GO datasets (GSE211106, GSE197999, GSE144796, GSE26887, and GSE215979) in mouse models (Fig. 1C). Transcriptomic sequencing analysis revealed no significant difference in METTL1 mRNA expression levels between the tissue of normal mice and the myocardial tissue of diabetic mice (Fig. S1A). Subsequent studies on PBMCs from DCM patients and healthy donors and heart tissues from a mouse model of spontaneous diabetes (db/db) or a type 2 diabetic mouse model induced by streptozotocin (STZ) combined with high-fat diet also revealed no significant difference in METTL1 mRNA expression (Fig. 1D-F). Enzyme-linked immunosorbent assays (ELISAs) were then conducted to determine the METTL1 protein level in the peripheral blood mononuclear cells (PBMCs) of DCM patients, which revealed decreased expression compared with that in healthy individuals (Fig. 1I). Furthermore, a negative correlation was observed between the expression levels of the METTL1 protein and NT-proBNP and sST2 in the peripheral blood of DCM patients, suggesting a potential association with heart failure and myocardial fibrosis in these patients (Fig.1G and H). One year later, follow-up studies confirmed the decreased METTL1 protein expression in DCM patients (Fig. 1L), with a continued negative correlation with NT-proBNP and sST2 expression (Fig. 1J and K). Immunoblotting analysis of METTL1 expression in the heart tissues of different diabetic mice revealed decreased expression in the myocardial tissues of diabetic mice (Fig. 1M-O) and in the PBMCs of patients with type 1 and type 2 diabetes (Fig. 1P).

**Figure 1.**
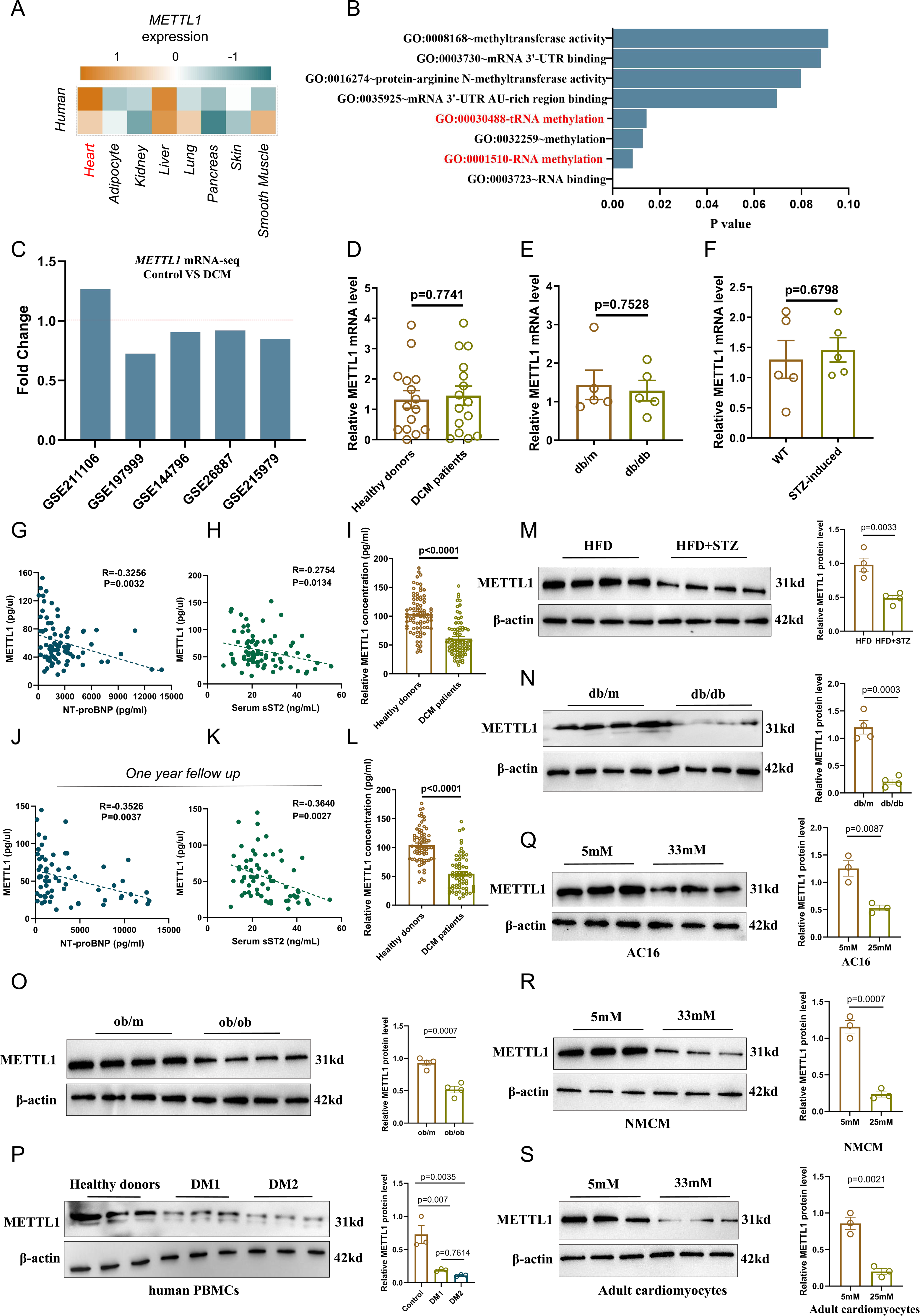
Diabetic cardiomyocytes exhibited a significant reduction in METTL1 protein expression levels *in vivo* and *in vitro*. (A) The main tissue-specific METTL1 expression profile was shown from BioGPS database. (B) Changes in RNA methylation and tRNA methylation involved in diabetic cardiomyopathy mice analysed by Gene Ontology (GO) using GTEx Dataset data (GSE173384). (C) METTL1 expression in diabetic cardiomyopathy mice was showed from Go Dataset data (GSE211106, GSE197999, GSE144796, GSE26887, GSE215979). (D) mRNA expression level of METTL1 in PBMCs (Peripheral Blood Mononuclear Cells) derived from DCM patients (n=15) and healthy donors (n=15). Data are expressed as mean ± SEM. (E-F) Detection of METTL1 mRNA level in heart tissue from spontaneous diabetes (db/db) mice (n = 5) and STZ combined with high-fat diet induced diabetic mice (n=5) using qPCR assay. Data are expressed as mean ± SEM. (G) The spearman correlation analysis for expression of METTL1 concentration detected via ELISA assay in PBMCs and serum NT-proBNP levels from DCM patients (n = 80) (R = -0.3256. p = 0.0032). (H) The spearman correlation analysis for expression of METTL1 concentration detected via ELISA assay in PBMCs and serum sST2 levels from DCM patients (n = 80) (R = -0.2754. p = 0.0134). (I) Expression of METTL1 concentration detected via ELISA assay in PBMCs from DCM patients and healthy donors. (J-K) After one-year follow up, the expression levels of METTL1 in PBMCs, serum NT-proBNP and sST2 were detected using ELISA assay and showed by spearman correlation analysis. (L) Expression of METTL1 concentration detected via ELISA assay in PBMCs from DCM patients and health donors after one-year follow up. (M) The expression protein level of METTL1 from heart tissues from Control and diabetic mice via immunoblotting analysis. Data are expressed as mean ± SEM. (N) The expression protein level of METTL1 in heart tissue from db/m or db/db diabetic mice model via immunoblotting analysis. Data are expressed as mean ± SEM. (O) The expression protein level of METTL1 in heart tissue from ob/m or ob/ob diabetic mice model via immunoblotting analysis. Data are expressed as mean ± SEM. (P) The expression protein level of METTL1 from PBMCs from healthy donors and patients with type 1 and type 2 diabetes via immunoblotting analysis. Data are expressed as mean ± SEM. (R-S) The expression protein level of METTL1 was detected in human cardiomyocyte-like AC16 cells or NMCM (neonatal mouse cardiac-myocyte) cells or 8-week-old adult mouse cardiomyocytes induced by low (5mM) or high glucose (33mM) for 24 hours (24h). Data are expressed as mean ± SEM.

To elucidate the expression pattern of the METTL1 protein in diabetic heart tissue, we performed an analysis of METTL1 expression levels using single-cell datasets from the HPA database (Fig. S1D). The analysis revealed high expression of METTL1 in cardiomyocytes and myocardial fibroblasts. Subsequent investigations utilizing the Tabula Muris database and Single Cell Portal database also confirmed the high expression of METTL1 in cardiomyocytes and myocardial fibroblasts, with relatively low expression in endothelial cells and lymphocytes (Fig. S2A-C). Quantitative PCR was employed to assess METTL1 expression levels in various components of ob/ob mouse heart tissue following separation and extraction of mouse immune cells, including T-cell sorting. The findings indicated that the mRNA expression levels of METTL1 in the cardiac tissue (CMs), cardiac fibroblasts (CFs), endothelial cells (ECs), bone marrow-derived macrophages, and spleen T lymphocytes of ob/ob mice did not significantly differ from those of ob/m mice (Fig. S1E). Immunoblotting assays revealed no significant changes in METTL1 protein levels in CFs, CMs, or bone marrow-derived macrophages (Fig. S2E-G). Furthermore, investigations into the impact of high glucose on METTL1 expression levels in cardiomyocytes demonstrated that high glucose treatment inhibited METTL1 protein expression, a finding that was consistent with results from cultured mouse primary cardiomyocytes (Fig. 1Q-S). Concurrent assessment of WDR4, an essential interacting protein of METTL1, revealed no significant differences in WDR4 protein or mRNA levels between high-sugar-induced AC16 cardiomyocytes and primary cardiomyocytes (Fig. S2H-L). These results suggest that high glucose content primarily suppresses METTL1 protein expression in cardiomyocytes but has no significant effect on WDR4.

### Myocardial-specific knockout of METTL1 exacerbates the inflammatory response and myocardial hypertrophy in DCM

To further investigate the pivotal role of the METTL1 protein in DCM, we generated myocardial-specific knockout mice with tamoxifen-induced METTL1 deficiency (Fig. 2A and Fig. S3A). Immunoblotting assays revealed a significant decrease in METTL1 protein expression in the heart following tamoxifen treatment, whereas no notable changes were observed in the lung, liver or kidney (Fig. S3B). This outcome indicates the successful generation of mice with cardiomyocyte-specific METTL1 knockout. We found no significant alterations in blood glucose levels or body weight due to METTL1 myocardial knockout (Fig. S3C and D). Additionally, no obvious differences were observed in METTL1^CKO^ and METTL1^Cre^ after treating with tamoxifen for 12 weeks in heart changes (Fig. S3E) and glucose tolerance (Fig. S3F). Using this model, we subsequently conducted a comprehensive study involving high-fat diet administration and STZ treatment. Three months after high-fat diet feeding and STZ intervention, we found no significant alterations in blood glucose levels or body weight due to METTL1 myocardial knockout (Fig. S4A and B). Glucose tolerance remained unchanged in the mice three months after modelling, as shown by the intraperitoneal glucose tolerance test (IPGTT) (Fig. S4C). Interestingly, the heart tissue of the METTL1^CKO^ mice was notably larger than that of the METTL1^Cre^ mice (Fig. 2B and C). Elevated BNP levels were found in the peripheral blood serum of the METTL1^CKO^ mice (Fig. 2D) but not in non-diabetic mice (Fig. S3G-I). After METTL1 knockout, a notable reduction in the left ventricular ejection fraction and left ventricular fractional shortening rate was observed, indicating a decline in heart function in the METTL1-deficient mice (Fig. 2E and F). Nevertheless, no significant changes in heart rate or blood pressure were noted in non-diabetic mice (Fig. S3K-M) and diabetic mice (Fig. S4D and E). Histological analyses using HE and Masson staining revealed substantial myocardial hypertrophy and fibrosis in the METTL1^CKO^ mice compared with those in the METTL1^Cre^ mice (Fig. 2J and K). WGA staining further demonstrated that knockout of the METTL1 protein in the myocardium increased myocardial hypertrophy in diabetic mice (Fig. 2L). Additionally, qPCR analysis revealed a significant increase in the expression levels of heart hypertrophy-related genes (Nppa, Myh7, Nppb, and TTNT2) following METTL1 knockout (Fig. 2G). Moreover, the expression of proinflammatory genes (IL-1β, IL-6, TNF-α, and Cxcl2) (Fig. 2H) and fibrosis-related genes (Col1α1, ACTA2, TGF-β1, and fibronectin) (Fig. 2I) was significantly upregulated. However, we found no significant changes in non-diabetic mice (Fig. S3J). The expression of cardiac hypertrophy-related proteins (CAMK2D, AGT, ADK, PARP1, GSN, PTK2, MTPN, CAV3, TTN, TCAP, and CSRP3) also increased after METTL1 knockout (Fig. 2M). Western blot analysis revealed significant increases in the expression levels of hypertrophic proteins (ANP and BNP), collagen I (αSMA), and inflammatory factors (IL-1β, IL-6, and TNF-α) in the heart tissue of the METTL1^CKO^ mice compared with those in the heart tissue of the METTL1^Cre^ mice (Fig. 2N-O). ELISA results for IL-1β, IL-6, TNF-α, TGF-β1 and IL-1α (Fig. 2P) in heart tissue further confirmed that knocking out cardiomyocyte METTL1 could increase the expression of hypertrophy-related proteins, profibrotic proteins, and proinflammatory factors in the heart.

**Figure 2.**
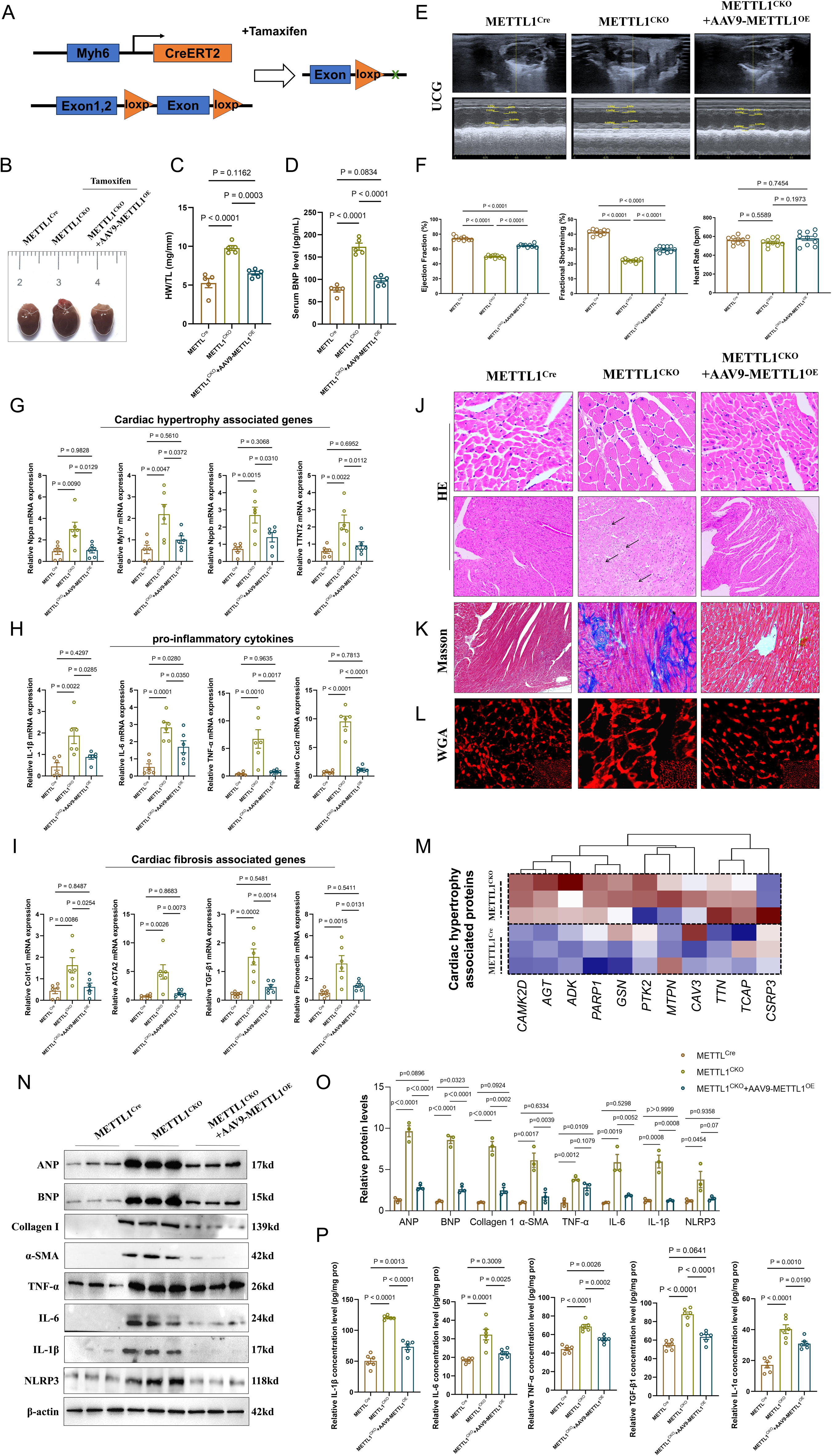
Cardiomyocyte-specific knockout of METTL1 exacerbated myocardial inflammation and fibrosis in diabetic mouse cardiomyocytes. (A) Schematic diagram of tamoxifen induced myocardial METTL1 knock-out mice model. Mice were dived into three groups: METTL1^Cre^ (METTL1^fl/fl^Myh6^Cre^ without tamoxifen induced), METTL1^CKO^ (METTL1^fl/fl^Myh6^Cre^ with tamoxifen induced) and METTL1^CKO^ + AAV9-METTL1^OE^ mice. (B) Representative general photograph of heart tissues, statistical analysis of ratio of heart weight to tibia length (HW/TL) and serum BNP levels compared among METTL1^Cre^, METTL1^CKO^ and METTL1^CKO^ + AAV9-METTL1^OE^ mice. Data are expressed as mean ± SEM. (E) Representative UCG images of METTL1^Cre^, METTL1^CKO^ and METTL1^CKO^ + AAV9-METTL1^OE^ mice. (F) Statistical analysis of ejection fraction, fractional shortening and heart rate among METTL1^Cre^, METTL1^CKO^ and METTL1^CKO^ + AAV9-METTL1^OE^ mice. (G) Comparison of relative mRNA expression levels of genes associated with cardiac hypertrophy (Nppa, Myh7, Nppb and TINT2) among METTL1^Cre^, METTL1^CKO^, and METTL1^CKO^ + AAV9-METTL1^OE^ mice. (H) Comparison of relative mRNA expression levels of genes associated with pro-inflammatory cytokines (IL-1β, IL-6, TNF-α and Cxcl2) among METTL1^Cre^, METTL1^CKO^, and METTL1^CKO^ + AAV9-METTL1^OE^ mice. (I) Comparison of relative mRNA expression levels of genes associated with cardiac fibrosis (Col1α1, Acta2, TGF-β1 and Fibronectin) among METTL1^Cre^, METTL1^CKO^, and METTL1^CKO^ + AAV9-METTL1^OE^ mice. (J-L) HE, Masson and WGA staining of myocardial tissues among METTL1^Cre^, METTL1^CKO^, and METTL1^CKO^ + AAV9-METTL1^OE^ mice. (M) Expression of cardiac hypertrophy associated proteins via proteomics analysis in METTL1^Cre^ and METTL1^CKO^ mice. (N-O) Representative images and statistical analysis of ANP, BNP, Collagen I, α-SMA, TNF-α, IL-6, IL-1β and NLRP3 were detected by immunoblotting analysis in heart tissues from METTL1^Cre^, METTL1^CKO^, and METTL1^CKO^ + AAV9-METTL1^OE^ mice. (P) Relative concentration levels of pro-inflammation factors including IL-1β, IL-6, TNF-α, TGF-β1 and IL-1α in heart tissues among METTL1^Cre^, METTL1^CKO^, and METTL1^CKO^ + AAV9-METTL1^OE^ mice via ELISA assay. Data are expressed as mean ± SEM.

### AAV9-mediated re-expression of METTL1 rescues inflammation and cardiac remodeling in METTL1-deficient mice

To determine whether restoration of METTL1 expression could reverse the pathological phenotypes caused by METTL1 deficiency, we delivered AAV9 vectors expressing METTL1 (AAV9-METTL1^OE^) into METTL1^CKO^ mice. AAV9-METTL1^OE^ injection effectively restored cardiac METTL1 expression and gross morphology showed reduced cardiac enlargement compared to untreated METTL1^CKO^ mice (Fig. 2B). Serum BNP levels were significantly elevated in METTL1^CKO^ mice, and were markedly reduced upon METTL1 rescue (Fig. 2C and D).

Echocardiography revealed that METTL1^CKO^ mice exhibited decreased ejection fraction and fractional shortening, consistent with impaired systolic function, whereas AAV9-METTL1^OE^ partially restored cardiac performance (Fig.2E and F). RT-qPCR analyses showed that the upregulation of cardiac hypertrophy-related genes (e.g. Nppa, Nppb, Myh7), proinflammatory cytokines (e.g. IL-1β, IL-6, TNF-α), and fibrosis-associated markers (e.g., Col1α1, ACTA2) in METTL1^CKO^ hearts was significantly attenuated upon METTL1 re-expression (Fig. 2G-I).

Histological analyses further confirmed that AAV9-mediated METTL1 rescue alleviated structural disarray (HE), collagen deposition (Masson), and cardiomyocyte hypertrophy (WGA staining) seen in METTL1^CKO^ mice (Fig. 2J-L). Western blotting and ELISA also confirmed that METTL1 re-expression normalized the expression of pathological markers including ANP, BNP, collagen I, α-SMA, and inflammatory proteins such as TNF-α, IL-1β, and NLRP3 (Fig. 2N-P).

Collectively, these results demonstrate that AAV9-mediated METTL1 rescue effectively reverses cardiac remodeling, fibrosis, and inflammation in METTL1-deficient mice, establishing METTL1 loss as a causative driver of diabetic cardiomyopathy.

### Myocardial-specific knockout of METTL1 exacerbates the inflammatory response and myocardial hypertrophy in spontaneous ob/ob diabetic mice

To elucidate the role of METTL1 in DCM, we generated METTL1^CKO^ ob/ob mice by crossing METTL1^CKO^ mice with ob/m mice, which exhibit spontaneous diabetes (Fig. S5A). Subsequent investigations revealed that deletion of METTL1 led to a significant decrease in the left ventricular ejection fraction and left ventricular fractional shortening in ob/ob mice with diabetes (Fig. S5B, E and F). Interestingly, the heart tissue of the METTL1^CKO^ mice was notably larger than that of the METTL1^Cre^ mice (Fig. S5C). Histological analyses using HE and Masson staining demonstrated notable hypertrophy of myocardial cells and severe fibrotic changes in the hearts of ob/ob mice following METTL1 knockout (Fig. S5D). Additionally, WGA staining revealed a notable increase in cardiomyocyte hypertrophy in ob/ob mice upon METTL1 deletion (Fig. S5D). BNP level was highly elevated in METTL1 knockout mice (Fig. S5G). Western blot analysis revealed significant increases in the expression levels of hypertrophy-related proteins (ANP and BNP), fibrosis-related proteins (collagen I, α-SMA), and inflammatory-related factors (NLRP3, ASC, IL-1β, IL-6, and TNF-α) (Fig. S5H-I) in the heart tissue of the METTL1^CKO^ mice compared with those of the METTL1^Cre^ mice with an ob/ob background.

Furthermore, qPCR analysis revealed a significant increase in the expression levels of heart hypertrophy-related genes (Nppa, Myh7, Nppb, and TTNT2) (Fig. S5J) following METTL1 knockout. The expression of proinflammatory genes (IL-1β, IL-6, TNF-α, and Cxcl2) (Fig. S5K) and fibrosis-related genes (Col1α1, ACTA2, TGF-β1, and fibronectin) (Fig. S5L) was significantly upregulated.

### METTL1 knockout reprograms fatty acid metabolism in the cardiomyocytes of diabetic mice

Metabolic remodelling has been identified as a critical mechanism contributing to target organ damage in individuals with diabetes mellitus. This study aimed to investigate the association between m7G modification and cardiomyocyte metabolism in diabetic mice (Fig. 3A). Quantitative proteomic and metabolomic analyses using TMT labelling were conducted on METTL1^CKO^ and control METTL1^Cre^ mice. The findings revealed that the differentially expressed proteins in the METTL1^CKO^ mice and control mice were involved primarily in biological processes such as fatty acid β-oxidation and fatty acid synthesis (Fig. 3B, S6 C and D). Additionally, gene set enrichment analysis (GSEA) indicated that compared with that of the METTL1^Cre^ mice, the fatty acid decomposition capacity of the METTL1^CKO^ mice was lower, whereas cholesterol metabolism was significantly greater (Fig. 3C). Further elucidation of the role of these proteins in the fatty acid β-oxidation pathway was achieved through a model diagram, which demonstrated that key proteins associated with fatty acid β-oxidation (ACADM, ACADL, ACAA, HADHA, etc.) in the cardiac tissue of the METTL1^CKO^ mice were notably decreased compared with those in the METTL1^Cre^ mice (Fig. 3D and E). Moreover, the fatty acid mitochondrial transporter CPT2 was significantly reduced (Fig. 3D). Concurrently, metabolic enzymes involved in acetyl-CoA metabolism and the subsequent TCA cycle were significantly reduced (Fig. S6E-G). These results suggest that the oxidative process of fatty acid β-oxidation in the myocardial tissue of diabetic mice is compromised following METTL1 deletion, leading to a substantial decrease in the oxidative phosphorylation capacity. Principal component analysis (PCA) revealed significant differences in cardiometabolic profiles (Fig. 3F) between the METTL1^CKO^ mice and METTL1^Cre^ mice. Furthermore, metabolites related to the TCA cycle and fatty acid metabolism were notably diminished in the METTL1^CKO^ mice compared with the control mice, whereas metabolites in the cholesterol anabolic pathway were significantly increased (Fig. 3G and S7A). Kyoto Encyclopedia of Genes and Genomes (KEGG) analysis of selected differentially abundant metabolites revealed that, relative to those in the control mice, the differentially abundant metabolites in the heart tissues of the METTL1^CKO^ mice were predominantly associated with biological processes such as the TCA cycle, fatty acid synthesis and degradation, and oxidative phosphorylation (Fig. 3H). BODIPY probe labelling of primary mouse cardiomyocytes revealed that lipid droplet synthesis was induced by high glucose. Compared with those of Cre mice, the primary cardiomyocytes of CKO mice presented increased lipid droplet synthesis, potentially related to impaired fatty acid metabolism in cardiomyocytes following METTL1 regulation (Fig. 3I). The fatty acid decomposition level in cardiomyocytes was assessed through FAO metabolism. The results indicated that reducing METTL1 expression in AC16 cells attenuated the fatty acid decomposition triggered by stock, whereas METTL1 overexpression amplified the fatty acid decomposition in AC16 cells induced by high glucose (Fig. 3J) and in NMCM cells (Fig. S7D). Additionally, oxygen consumption rate (OCR) analysis revealed that METTL1 overexpression increased oxidative phosphorylation in cardiomyocytes stimulated with high glucose, whereas oxidative phosphorylation levels in cardiomyocytes were notably reduced following METTL1 knockdown (Fig. 3K and Fig. S7E). ELISA measurements of metabolites associated with fatty acid decomposition demonstrated that, compared with Cre mice, CKO mice presented significantly elevated levels of free fatty acids and palmitic acid in cardiomyocytes, along with decreased levels of NADPH/NADP+, acetyl-CoA, and ATP (Fig. 3M). Furthermore, in heart tissue from ob/ob background mice, compared with Cre mice, CKO mice presented significantly increased levels of free fatty acids in cardiomyocytes, along with decreased ACS synthetase activity and β-hydroxybutyric acid, ethyl acetate, acetyl-CoA, and ATP levels (Fig.3L). However, METTL1 over-expression in AC16 could rescue this metabolic process (Fig. S7F). Quantitative PCR analysis revealed that key genes involved in cholesterol synthesis were significantly upregulated in CKO mice compared with Cre mice, whereas the expression of fatty acid β-oxidation-related genes was notably downregulated (Fig. 3N). However, we found no significant differences in the key genes related to fatty acid synthesis (Fig. 3N). Additionally, the expression levels of key genes involved in oxidative phosphorylation metabolism were significantly lower in CKO mice than in Cre mice (Fig. 3O). We confirmed this conclusion in METTL1^CKO^ with ob/ob background mice (Fig. S7B). Also, we analysis the results of RNA-seq from ob/m and ob/ob mice (GSE173384) and found no differences of genes associated with Fatty acid metabolism and glycolysis pathway (Fig. S7G). These results indicated that changes of genes profile in METTL1^CKO^ ob/ob mice was mainly attribute to the effect of METTL1 but not ob/ob background Overall, these findings suggest that METTL1 deficiency induced by high glucose levels may diminish fatty acid decomposition and oxidative phosphorylation in cardiomyocytes, leading to the accumulation of fatty acids and excessive lipid droplet production. Excessive fatty acid oxidation (FAO) and metabolic disturbances after METTL1 deletion could result in direct toxic effects of fatty acids on cardiomyocytes, provoke proinflammatory responses, and drive the pathological progression of DCM.

**Figure 3.**
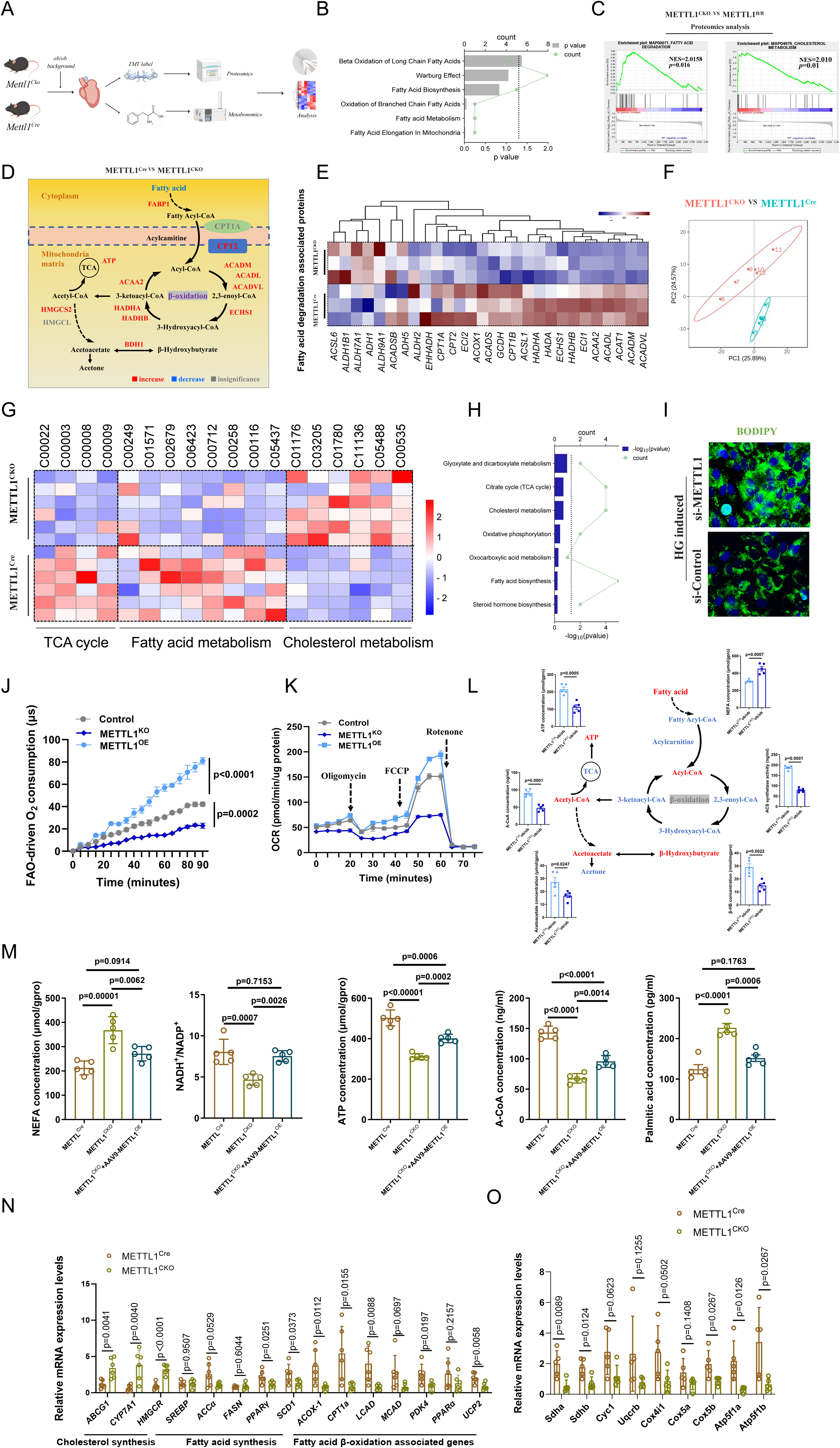
METTL1 knockout reprogrammed fatty acid metabolism in the cardiomyocytes of diabetic mice. METTL1^Cre^ and METTL1^CKO^ mice were treated with HFD+STZ to constructed the diabetic mice model and cardiac tissues were collected for TMT labeled proteomics assay and metabonomics analysis. (A) Graphic abstract of the whole experimental process. (B) Differential proteins between METTL1^Cre^ and METTL1^CKO^ mice was analysed by via KEGG pathway analysis online (DAVID website). (C) GSEA analysis for differential proteins in proteomics analysis between METTL1^Cre^ and METTL1^CKO^ mice. (D) Metabolic pathway map shown the differential proteins involved in fatty acid metabolism in METTL1^Cre^ and METTL1^CKO^ mice. Red indicted up-regulation proteins, blue indicted down-regulated proteins (METTL1^Cre^ compared with METTL1^CKO^ mice). (E) Heatmap expression levels of fatty acid degradation associated proteins via proteomics analysis in heart tissues from METTL1^Cre^ and METTL1^CKO^ mice. (F) PCA analysis for the metabonomics analysis of heart tissues from METTL1^Cre^ and METTL1^CKO^ mice (n = 6) (Red indicated METTL1^CKO^ group and blue indicated the METTL1^Cre^ group). (G) Heatmap expression levels of metabolites associated TCA cycle, fatty acid metabolism and cholesterol metabolism in heart tissues from METTL1^Cre^ and METTL1^CKO^ mice. (H) KEGG analysis for the differential metabolites in heart tissues from METTL1^Cre^ and METTL1^CKO^ mice in MetaboAnalyst 5.0 website. (I) Representative image of BODIPY staining for the AC16 cell line transfected with METTL1 small interference RNA (si-METTL1) and NC group (si-NC) after high glucose induced for 24h. (J) Fatty Acid Oxidation (FAO) colorimetric assay for METTL1 knockdown (METTL1^KO^) and METTL1 overexpression (METTL1^OE^) AC16 cells *in vitro* induced via high glucose for 24 h. (K) Seahorse analysis for the O_2_ consumption rate (OCR) for METTL1 knockdown (METTL1^KO^) and METTL1 overexpression (METTL1^OE^) AC16 cells *in vitro* induced with high glucose for 24 h. (L) Statistical analysis of metabolites of fatty β-oxidation pathway in heart tissues from METTL1^Cre^ob/ob and METTL1^CKO^ob/ob mice at age 16 weeks after induced via tamoxifen for 8 weeks. (M) Statistical analysis of metabolites of fatty β-oxidation pathway in heart tissues from METTL1^Cre^ and METTL1^CKO^ diabetic mice. (N) Relative expression levels of genes associated with cholesterol metabolism, fatty acid synthesis and fatty acid β-oxidation pathways in heart tissues from METTL1^Cre^ and METTL1^CKO^ diabetic mice. (O) Relative expression levels of genes associated with TCA cycle pathway in heart tissues from METTL1^Cre^ and METTL1^CKO^ diabetic mice.

### METTL1 modulates the m7G modification and mRNA stability of crucial enzymes involved in fatty acid metabolism in cardiomyocytes

Since METTL1 plays a crucial role as an enzyme in m7G modification, we hypothesized that the regulation of fatty acid metabolism in cardiomyocytes by METTL1 might be dependent on the m7G modification of downstream target genes by METTL1. To investigate this hypothesis, we performed m7G-seq analysis on the cardiac tissue of the aforementioned mice (Fig. 4A). m7G-seq analysis revealed that m7G modification predominantly occurred in the 5’ untranslated region (UTR) of mRNA (28.8% in Cre; 30.92% in CKO) and the CDS region, with a smaller distribution in the 3’ UTR (Fig. 4B and S8A-8B). Additionally, 27,254 peaks were identified in the diabetic heart tissues (Fig. 4C and D and Fig. S8C). Subsequently, KEGG analysis was performed on 335 differentially expressed genes related to metabolism to further elucidate the differential peaks and primary functions of the genes following METTL1 knockout (Fig. 4E). The results revealed significant enrichment in pathways such as fatty acid biosynthesis, fatty acid metabolism, glycolysis/gluconeogenesis, and the PPAR signalling pathway (Fig. 4E). Furthermore, the KEGG analysis demonstrated that the genes whose expression decreased in CKO mice were notably enriched in the apelin signalling pathway (Fig. 4F and Fig. S8F). These findings suggest that the m7G modification level of crucial genes involved in the activation of the apelin signalling pathway in cardiomyocytes was significantly reduced after METTL1 knockout. Apelin is a pivotal signalling molecule that regulates glucose and lipid metabolism in cardiomyocytes, and the suppression of the apelin signalling pathway may be pivotal in the impairment of fatty acid metabolism in cardiomyocytes after METTL1 knockout. Moreover, through the intersection enrichment analysis of the quantitative proteomic outcomes of M7G-modified differentially expressed genes and TMT markers, proteins associated with fatty acid metabolism (Scp2, Ehhadh, Scd2, Acsl5, Elovl6, Acsl4, Scd1, Hacd3, Acsf3, Mcat, Acc, Fads1), glycolysis and TCA cycling proteins (G6pc3, Adpgk, Aldh3b1, Pgm2, Pck1, Aldoa, Eno2, Pck2), and the apelin signalling pathway (Plcb1 Plcb2, Hdac4, Pik3r6, Hras, Adcy3, Sphk2, Mylk, Itpr2, Map2k2, Mapk1, Klf2, Slc8a1, Gna13) exhibited notable enrichment (Fig.4G). The model diagram revealed an increase in the m7G level of key proteins involved in lipid droplet synthesis following METTL1 knockout, whereas the m7G modification level of key proteins involved in fatty acid decomposition notably decreased (Fig. 4H). Visual analysis using Integrative Genomics Viewer (IGV) revealed a significant reduction in the m7G levels of key enzymes involved in glycolipid metabolism, such as Scd1, Aldoa, Sdha, and Ascl5, after METTL1 knockout (Fig. 4I). m7G modification of these key proteins may lead to increased lipid droplet synthesis, reduced fatty acid decomposition, and dysfunction in oxidative phosphorylation in cardiomyocytes induced by hyperglycaemia. The restructuring of fatty acid metabolism in cardiomyocytes results in lipid droplet peroxidation, decreased ATP synthesis, inflammation, energy deficiency, and ultimately pathological remodelling of cardiomyocytes.

**Figure 4.**
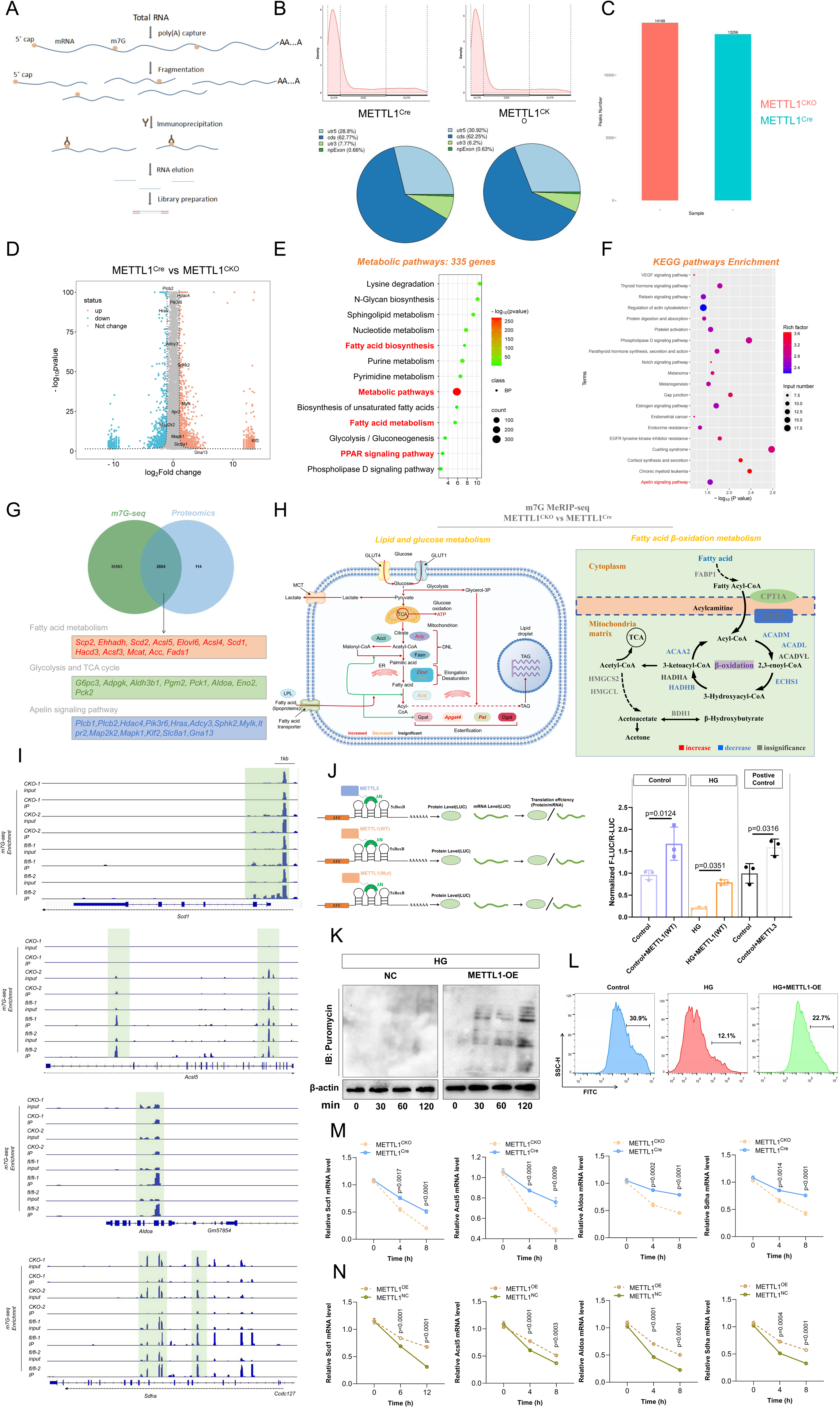
METTL1 modulated the m7G modification and mRNA stability of crucial enzymes involved in fatty acid metabolism in cardiomyocytes. Total RNA was isolated from myocardial tissue of METTL1^Cre^ and METTL1^CKO^ diabetic mice for m7G-MeRIP-seq. (A) Schematic diagram of m7G-MeRIP-seq experiment process. (B) Distribution of m7G peaks across mRNA segments. The pie chart showed the percentage of m7G site in each segment. (C) m7G methylated peaks number of mRNA in heart tissues after m7G-MeRIP-seq analysis. (D) Differential genes compared between METTL1^CKO^ and METTL1^Cre^ mice after m7G-MeRIP-seq analysis were shown by Volcano plots. (E) GO analysis for the differential genes involved in metabolic pathways after m7G-MeRIP-seq analysis. (F) Differential genes were analysed by KEGG pathways. (G) Venn diagram showed the same difference genes analysis of m7G-MeRIP-seq and Proteomics involving in fatty acid metabolism, glycolysis and TCA cycle and apelin signaling pathway. (H) Lipid synthesis pathway map shown the differential genes of m7G-MeRIP-seq involved in fatty acid metabolism in METTL1^Cre^ and METTL1^CKO^ mice (left). Fatty acid β-oxidation metabolism pathway map shown the differential genes of m7G-MeRIP-seq in METTL1^Cre^ and METTL1^CKO^ mice (right) (right means the up-regulation levels of genes, blue means the down-regulation levels of genes, METTL1^CKO^ compared with METTL1^Cre^ mice). (I) Integrative Genomics Viewer (IGV) shown the m7G peak of differential genes associated fatty acid metabolism pathway. (J) Statistical analysis for the luciferase assay of METTL1 activity in regulating the translation efficiency in AC16 cells. (K) Immunoblotting analysis for the METTL1 over-expression stable AC16 cell lines treating with Puromycin for different time points. (L) Flow cytometric analysis for the detection level of the protein synthesis. (M) NMCM cells from METTL1^Cre^ and METTL1^CKO^ mice were cultured and treated with CHX for different durations and cells were collected for further qPCR assays to determine the half-life of multiple genes associated fatty acid metabolism. Data are expressed as the mean ± SEM. (N) Stable METTL1-overexpression (METTL1^OE^) AC16 cells were cultured and treated with CHX for different durations and cells were collected for further qPCR assays to determine the half-life of multiple genes associated fatty acid metabolism. Data are expressed as the mean ± SEM.

Research has indicated that RNA m7G modification can affect intracellular translation processes. In this study, luciferase reporter experiments were conducted on cardiomyocytes following METTL1 knockout to assess the translation efficiency of nascent peptides (Fig. 4J). These findings revealed a decrease in the expression of firefly luciferase in cardiomyocytes after METTL1 knockout, suggesting a reduction in protein synthesis in these cells. Additionally, analysis of nascent peptide levels following puromycin stimulation and nascent peptides label assay revealed that METTL1 overexpression augmented neonatal peptide synthesis in cardiomyocytes (Fig. 4K and L). High glucose levels hindered peptide production in cardiomyocytes; however, the overexpression of METTL1 mitigated this effect. As the previous research indicated the m7G modification of mRNA also affected its degradation efficiency in cells. We then treated the cells with cycloheximide (CHX) and detected the half-life time of mRNA associated with fatty acid metabolism pathway. We found that METTL1 over-expression could delay the degradation of mRNA, while METTL1 knockout contribute to the degradation (Fig.4M and N). All these results indicated that METTL1 could regulate the m7G modification and mRNA stability of crucial enzymes involved in fatty acid metabolism in cardiomyocytes.

### High glucose induces O-GlcNAcylation and proteasome-mediated degradation of ubiquitinated METTL1 but not lysosomal degradation

This study revealed that high glucose levels did not significantly affect the transcriptional level of the METTL1 gene; however, they did affect METTL1 protein expression (shown in Fig.1). These findings suggest that the regulation of METTL1 by high glucose may be associated with post-translational modification of the protein. Research has demonstrated that glucose can generate UDP-GlcNAc molecules through the activity of various crucial metabolic enzymes. Subsequently, UDP-GlcNAc molecules can facilitate the O-GlcNAcylation of target proteins under the influence of the OGT protein (Fig. 5A). This investigation aimed to determine whether high glucose content induced O-GlcNAcylation of METTL1. To validate this hypothesis, we conducted glycoproteomic analysis on ob/ob mice and ob/m mice (Fig. 5B). The findings revealed 362 O-GlcNAc sites and a total of 179 proteins (Fig. S9B and C). Additionally, biological process (BP) analysis highlighted significant enrichment of metabolic pathways among the different proteins (Fig. S9D). GO analysis of these diverse proteins revealed that O-GlcNAcylated proteins are involved in glucose metabolism, glycolysis, the PPAR signalling pathway, RNA methylation regulation, and myocardial cytoskeletal function in cardiomyocytes (Fig. 5C) and KEGG shown that enrichment of O-gylan or N-gylan biosynthesis in ob/ob mice (Fig. 5D). To support the outcomes of glycoproteomics, we confirmed the substantial colocalization between OGT and METTL1 through fluorescence colocalization experiments (Fig. 5E). Furthermore, coimmunoprecipitation experiments on HL-1 mouse cardiomyocytes, AC16 human cardiomyocytes, and primary lactating mouse cardiomyocytes following high-glucose stimulation demonstrated that high glucose increased the level of O-GlcNAcylation of METTL1 (Fig. 5F). To further investigate the specific sites of O-GlcNAcylation of the METTL1 protein induced by high glucose, we predicted potential O-GlcNAcylation sites of METTL1 using the YinOYang 1.2website. The analysis suggested that T58, T268, and T269 could be crucial sites for O-GlcNAcylation modification of the METTL1 protein (Fig. 5G). For identification of the specific site of O-GlcNAcylation modification of METTL1, the three aforementioned amino acid sites were subsequently individually mutated to alanine (T to A). Coimmunoprecipitation assays were then conducted to assess the levels of O-GlcNAcylation. The results indicated that O-GlcNAcylation of METTL1 occurred predominantly at tyrosine 286 (T268) (Fig. 5H).

**Figure 5.**
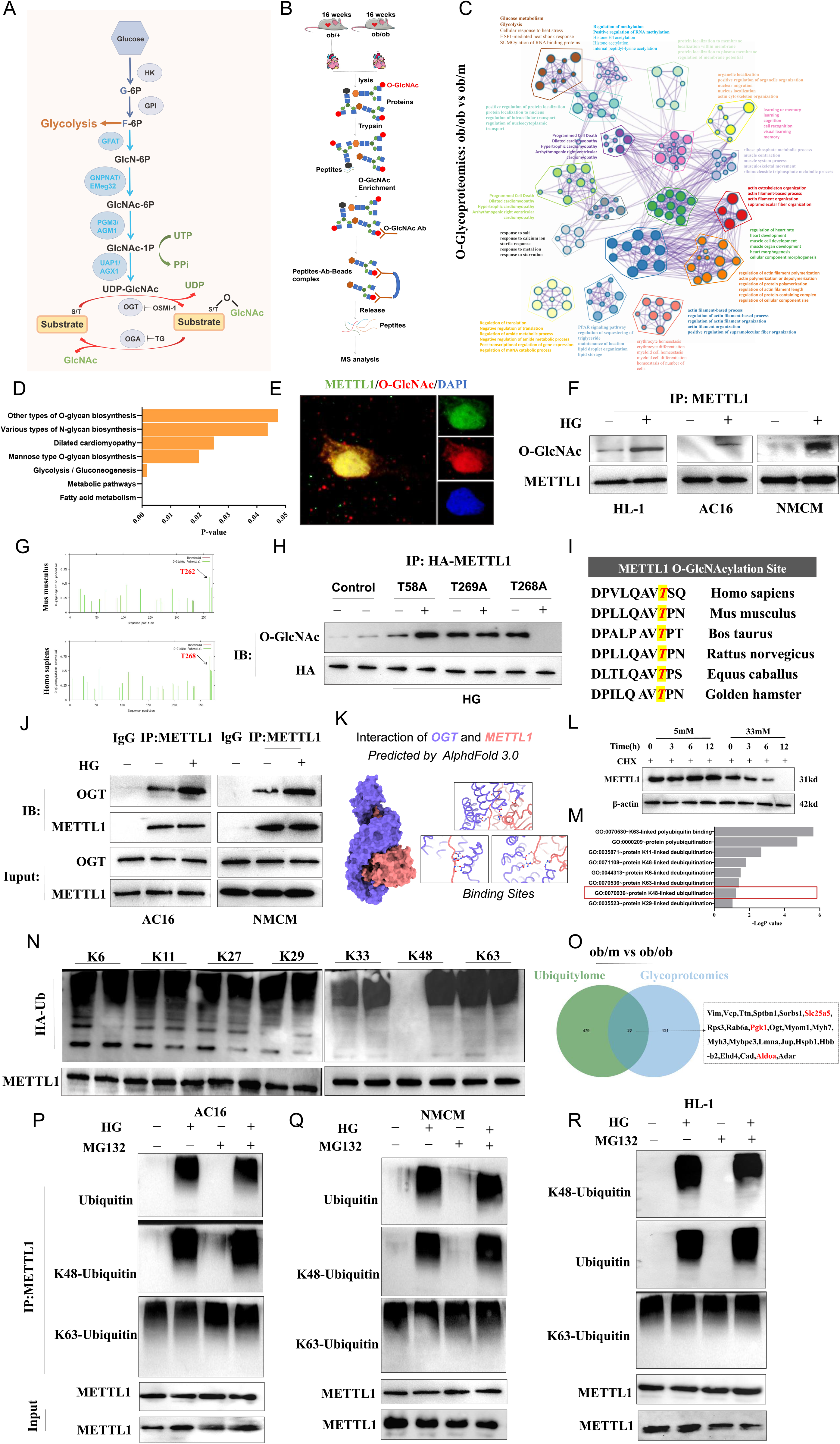
High glucose induced O-GlcNAcylation and K48-linked polyubiquitination-mediated degradation of METTL1. (A) Schematic diagram of metabolic process from glucose to UDP-GlcNAc on substrate protein modification. (B) Schematic diagram of Glycoproteomics analysis detected in myocardial tissue taken from 16-week-old ob/m and ob/ob mice. (C)The main cellular function enrichment analysed by enrichment pathway after Glycoproteomics detection. (D) KEGG enrichment pathway in ob/m and ob/ob mice. (E) Co-location assay shown the intracellular interaction between METTL1 and O-GlcNAc in AC16 cell. DAPI shown the nucleus, green shown the METTL1. red shown the O-GlcNAc. (F) AC16, NMCM and HL-1 cells were treated with 33mM high glucose for 24h and cells were collected for further assay. Co-immunoprecipitation was used to detect the O-GlcNAcylation modification level of METTL1 in AC16, NMCM and HL-1 cells. (G) Potential O-GlcNAcylation modification sites of METTL1 protein in *Homo sapiens* and *Mus musculus* were predicted using YinOYang 1.2 website (https://services.healthtech.dtu.dk/services/YinOYang-1.2/). (H) HEK29T cells were transfected with different potential O-GlcNAcylation modification HA-tag mutation sites METTL1 (threonine (T) to alanine (A) mutation). METTL1 (T58A, mutation site of threonine (T) to alanine (A) at 58 of METTL1 protein), METTL1 (T268A, mutation site of threonine (T) to alanine (A) at 268 of METTL1 protein) and METTL1 (T269A, mutation site of threonine (T) to alanine (A) at 269 of METTL1 protein) plasmids for 24h and then treated cells with high glucose for 24h for further immunoprecipitation assay to detected the O-GlcNAcylation modification level of METTL1. (I) Comparative analysis of METTL1 protein sequence conservation across species. (J) AC16 cells and NMCM cells were cultured and treated with high glucose for 24h and collected for the Co-immunoprecipitation assay to detected the interaction level of OGT and METTL1. (K) The structure of METTL1 and OGT were predicted via AlphaFold 3.0 and for further Molecular docking and molecular dynamics simulations analysis. (L) AC16 cells were treated with high glucose for 24h and then stimulated with Cycloheximide for different time points. Then collected the cells for further immunoblotting analysis at different time points. (M) GO analysis shown that the differential genes in proteomic analysis in heart tissues from METTL1^Cre^ and METTL1^CKO^ mice. (N) HEK293T cells was transfected with different plasmids of poly-Ubiquitination site (K6, K11, K27, K29, K33, K48 and K63) of ubiquitin and Flag-METTL1 plasmid then treated cells with high glucose for 24h, then collected the cells for further immunoprecipitation. (O) Venn diagram showed the same difference genes analysis of Glycoproteomics and Ubiquitylome study involving in metabolism pathway. (P) AC16 cells were treated with high glucose and MG132 for 12h and then collected the cells for *in vivo* ubiquitination of METTL1. (Q) NMCM cells were treated with high glucose and MG132 for 12h and then collected the cells for *in vivo* ubiquitination of METTL1. (R) Mouse HL-1 cells were treated with high glucose and MG132 for 12h and then collected the cells for *in vivo* ubiquitination of METTL1. Data are expressed as mean ± SEM.

Sequence conservation analysis across multiple species revealed that tyrosine at position 268 is conserved in various species, including *Homo sapiens*, *Mus musculus*, *Bos taurus*, *Rattus norvegicus*, *Equus caballus*, and the *golden hamster* (Fig. 5I). The interaction between OGT and METTL1 was identified through quantitative immunoprecipitation experiments (Fig. 5J). These findings indicated that elevated glucose levels could promote the interaction between OGT and METTL1.

To further confirm the interaction between OGT and METTL1, we then carried out structural modelling and molecular dynamics (MD) analysis of the OGT-METTL1 complex. The amino acid sequences of OGT and METTL1 were retrieved from the UniProt database. We first predicted the OGT-METTL1 complex structure using AlphaFold3. Subsequent structural refinement was performed using Gromacs 2024.3. Stable MD trajectories were extracted for energy landscape and binding mode analyses (Fig. 5K). To investigate OGT-METTL1 interactions at the molecular level, we performed denovo modelling of their binding interface using AlphaFold3. Given the presence of extensive intrinsically disordered loop regions in the OGT and METTL1 structures that complicate structural analysis, we retained their core regions (residues 33-332 for OGT; 19-246 for METTL1) for AlphaFold3 server-based complex prediction (Fig. 5K). As shown in Fig. S10A and B, the predicted complex demonstrates excellent shape complementarity between OGT and METTL1. The predicted structure was subjected to 100-ns MD simulations using Gromacs 2024.3 to optimize structural stability. Root Mean Square Deviation (RMSD) analysis (Fig. S10C) revealed that METTL1 maintained structural stability throughout the simulation, whereas OGT exhibited greater conformational fluctuations before reaching equilibrium at 20 ns. The composite RMSD of the OGT-METTL1 complex stabilized after 20 ns, indicating system equilibration. Subsequent analyses focused on the stable trajectory segment (20-100 ns). Root Mean Square Fluctuation (RMSF) and B factor analyses (Fig. S10D) demonstrated that most residues in both proteins exhibited low fluctuations (< 0.2 nm), with METTL1 showing lower flexibility than OGT. Notably, higher RMSF values occurred in non-interfacial regions, whereas interfacial residues remained stable, suggesting that OGT-METTL1 binding promotes structural stabilization. Electrostatic surface potential calculations using Chimera X revealed complementary charge distributions at the binding interface (Fig. S10E), providing additional stabilization through long-range electrostatic interactions. Hydrogen bond analysis revealed an average of 20 persistent interfacial bonds (Fig. S11A). Binding free energy calculations via gmx MMPBSA (1,000 frames from the final 10 ns) yielded an average ΔG of -85.36 kcal/mol (GGAS = -980.75 kcal/mol; GSOLV = 895.39 kcal/mol) (Fig. S11B and C). Per-residue energy decomposition (Fig. S11D) identified key contributors of OGT at Arg69, Glu90, Asn223, Asp230, Ser255, Asn298, Asn301, and Asn332 and of METTL1 at Arg6, Val8, Pro14, Arg19, Gln23, Arg24, Ala31, and Arg246. Moreover, a quantitative immunoprecipitation assay was used to detect the interaction between the OGT and METTL1 proteins, and the results revealed that high glucose content can promote the interaction between OGT and METTL1 (Fig. 5J).

To further elucidate the mechanism of decreased protein expression of METTL1 induced by high glucose, we speculated that this process may be related to the regulation of degradation of METTL1 induced by high glucose content. Therefore, we treated AC16 cells with cycloheximide (CHX) to determine the half-life of the METTL1 protein, and the results revealed that high-glucose induction can significantly shorten the half-life of the METTL1 protein (Fig.5L and Fig. S13E), further confirming our speculation. Analysis of the ubiquitinated proteomics of DCM model mice revealed multiple abnormalities in ubiquitination pathways in the heart tissues of diabetic mice (Fig. S12A and B). KEGG analysis and GO analysis shown that the proteins in ubiquitinated proteomics was enrichment for the ATP and NADH metabolism pathway (Fig. S12C and F). To explore the specific mechanism further, we used MG132 and CQ to inhibit the proteasomal and lysosomal systems to explore the specific type of degradation of METTL1 controlled by high glucoses. We found that the inhibition of lysosomal progression by CQ did not rescue the decrease in METTL1 expression induced by high glucose, but MG132 rescued this process (Fig. S13A-D). These findings indicate that the high-sugar-induced degradation of METTL1 is dependent on the proteasome system. After HEK293T cells were transfected with K6, K11, K27, K29, K33, K48, or K63 plasmids, we further determined the levels of METTL1 under high-glucose conditions with the K6, K11, K27, K29, K33, K48, or K63 vectors, and the results revealed that high-sugar conditions promoted K48 polyubiquitination of METTL1 (Fig. 5N). Additionally, by stimulating HL-1 cells, AC16 cells, and NMCM cells with high-glucose content and performing immunoprecipitation, we found that high-glucose content promoted K48 polyubiquitination of the METTL1 protein but significantly affected K63 polyubiquitination (Fig. 5P-R). These results collectively suggest that high glucose promotes K48 polyubiquitination-mediated degradation of METTL1 in cardiac muscle cells.

Additionally, we found that multiple proteins involved in the glucose metabolism were modified via O-GlcNAcylation and ubiquitin (Fig. 5O). And the proteins associated the fatty acid metabolism were enrichment in glycoproteomic (Fig. S14A) and ubiquitinated proteomics (Fig. S14B). GSEA also shown that these proteins were involved in the Lipid metabolic pathway, RNA metabolic process and Heart development (Fig. S14E-G). These interesting indicted the regulation network in heart of diabetic mice was quite complex.

### O-GlcNAcylation at residue T268/T262 inhibits the m7G modification of METTL1 but not affect its K48-linked poly-ubiquitination modification

As the previous research the O-GlcNAcylation and K48-linked poly-ubiquitination modification of METTL1 in high glucose condition. We then continue to clarify the detailed mechanism of O-GlcNAcylation and K48-linked poly-ubiquitination modification of METTL1. How would the O-GlcNAcylation of METTL1 affect itself and if any links between two modifications of METTL1? To further explore this hypothesis, we mutated the T268 site from threonine to alanine (T to A, T268A) in AC16 cells and T262A mutation in mouse for *in vivo* assay (Fig. 6 A) and found by stimulating AC16 cells with puromycin and determining the levels of new peptides at different time points, we revealed that the O-GlcNAcylation mutation of METTL1 can strongly rescue the translational efficiency inhibited by high glucoses, promoting peptide translation and protein synthesis (Fig. 6B). Both luciferase experiments and flow cytometry with new peptide labelling revealed that the inhibition of METTL1 O-GlcNAcylation can significantly rescue the translational efficiency inhibited by high glucose (Fig. 6C-E). Furthermore, through m7G-RIP-qPCR experiments, we found that the reduction in METTL1 O-GlcNAcylation caused by the T268A mutation can increase the m7G modification levels of downstream tRNAs (AlaAGC, LysCTT, LysTTT, ArgTCT, VlaAAC, and GlyGCC) (Fig. 6F and G). All these results represented that inhibition of METTL1 protein O-GlcNAcylation might promote the formation of new peptides.

**Figure 6:**
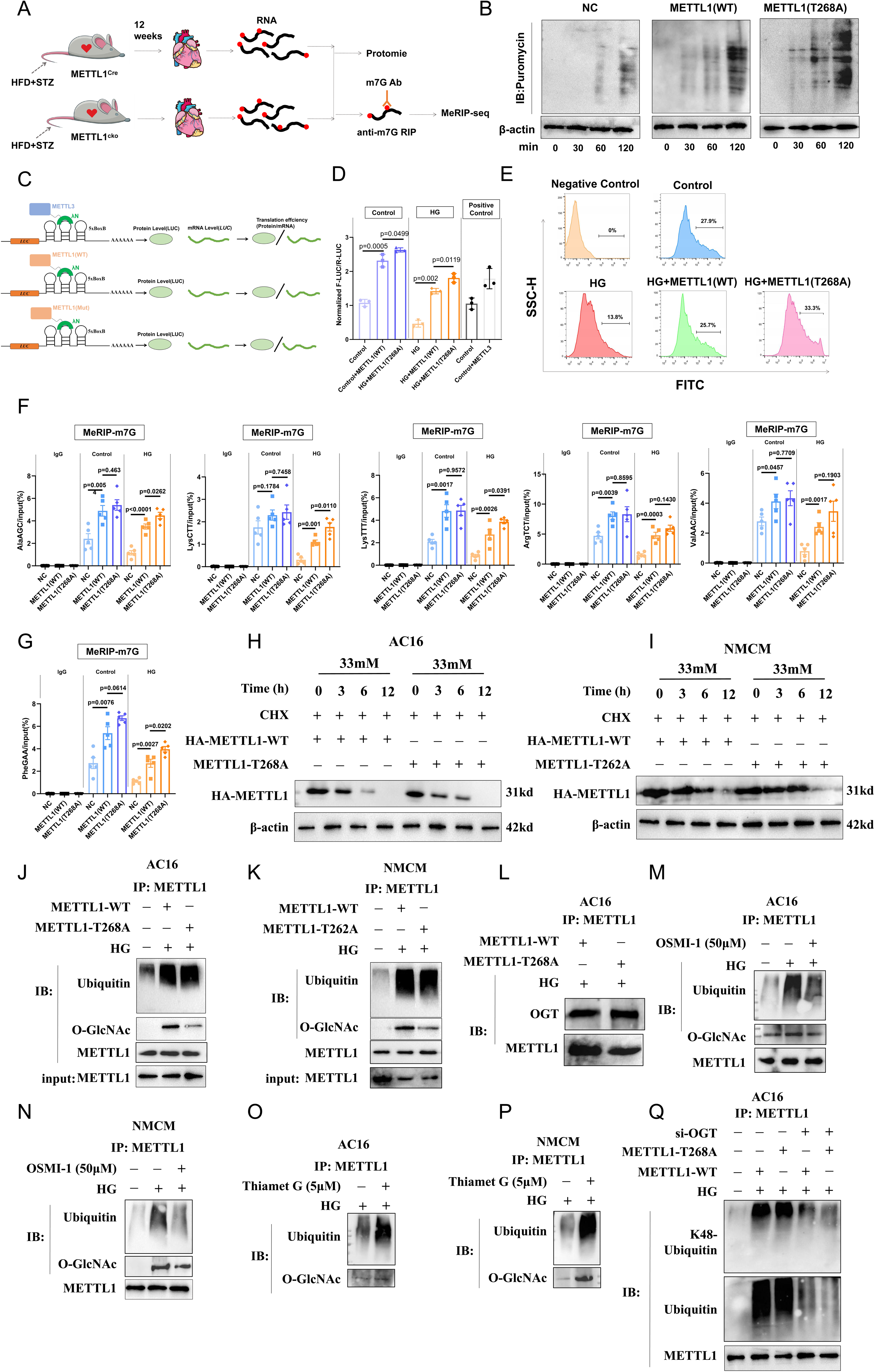
O-GlcNAcylation at residue T268 of METTL1 inhibited its m7G modification activity but not its K48 polyubiquitination. (A) Schematic diagram of heart tissues for m7G-MeRIP-seq and proteomics experiment process. (B) AC16 cells were transfected with vector plasmid, METTL1 (WT) plasmid and METTL1 (T268A) (mutation site of threonine (T) to alanine (A) at 268 of METTL1 protein). Immunoblotting analysis for the AC16 cells treating with Puromycin for different time points. (C-D) Statistical analysis for the luciferase assay of METTL1 activity in regulating the translation efficiency in AC16 cells transfected with vector plasmid, METTL1 (WT) plasmid and METTL1 (T268A) plasmid. (E) Flow cytometric analysis for the detection level of the protein synthesis. (F) AC16 cells were transfected with vector plasmid, METTL1 (WT) plasmid and METTL1 (T268A) plasmid and treated with high glucose (HG) for 24h and m7G-Methylation of RNA were measured via MeRIP-qPCR assay for tRNA-AlaAGC, tRNA-LysCTT, tRNA-lysTTT, tRNA-ArgTCT, tRNA-ValAAC. (G) MeRIP-qPCR assay for tRNA-PheGAA in AC16 cells after treated with high glucose for 24h. (H) AC16 cells were transfected with vector plasmid, METTL1 (WT) plasmid and METTL1 (T268A) plasmid and treated with high glucose for 24h and stimulated with Cycloheximide for different time points. The cells were all collected for further immunoblotting to detect the half-life time of METTL1(WT) and METTL1(T268A). (H) AC16 cells were transfected with vector plasmid, METTL1 (WT) plasmid and METTL1 (T268A) mutation plasmid and treated with high glucose for 24h and stimulated with Cycloheximide for different time points. The cells were all collected for further immunoblotting to detect the half-life time of METTL1 (WT) and METTL1 (T268A). (I) NMCM cells were transfected with vector plasmid, METTL1 (WT) plasmid and METTL1 (T262A) mutation plasmid and treated with high glucose for 24h and stimulated with Cycloheximide for different time points. The cells were all collected for further immunoblotting to detect the half-life time of METTL1 (WT) and METTL1 (T262A). (J) AC16 cells were transfected with FLAG-METTL1 and FLAG-METTL1 (T268A) plasmids and treated HG for 24h and collected the cells for further Co-IP to detected the *in vivo* ubiquitination of METTL1 and METTL1 (T268A). (K) NMCM cells were transfected with FLAG-METTL1 and FLAG-METTL1 (T262A) plasmids and treated HG for 24h and collected the cells for further Co-IP to detected the *in vivo* ubiquitination of METTL1 and METTL1(T262A). (L) AC16 cells were transfected with FLAG-METTL1 and FLAG-METTL1 (T268A) plasmids and treated HG for 24h and collected the cells for further Co-IP assay. (M) AC16 cells were treated with HG and OSMI-1 (50μM) for 24h and collected the cells for further Co-IP assay to detected the *in vivo* ubiquitination of METTL1. (N) NMCM cells were treated with HG and OSMI-1 (50μM) for 24h and collected the cells for further Co-IP assay to detected the *in vivo* ubiquitination of METTL1.(O) AC16 cells were treated with HG and Thiamet G (5μM) for 24h and collected the cells for further Co-IP assay to detected the *in vivo* ubiquitination of METTL1.(P) NMCM cells were treated with HG and Thiamet G (5μM) for 24h and collected the cells for further Co-IP assay to detected the *in vivo* ubiquitination of METTL1. (Q) AC16 cells were transfected with FLAG-METTL1 and FLAG-METTL1 (T268A) plasmids and small interfering RNA targeting for OGT. Cells were then treated with HG for 24h and collected the cells for further Co-IP assay.

Given that protein O-glycosylation can directly affect protein ubiquitination, we first speculated that high-sugar content may directly affect the ubiquitination-mediated degradation of METTL1 through O-GlcNAcylation at the T268 site. By transfecting AC16 cells with the T268 mutation and NMCM cells with the T262 mutation plasmid and stimulating them with high glucose, we found that neither the T268 mutation nor the T262 mutation can change the half-life of the METTL1 protein induced by high glucose (Fig. 6H and I). Furthermore, Co-immunoprecipitation assay revealed that neither the T268 mutation nor the T262 mutation inhibited the ubiquitination of METTL1 under high glucose condition, indicating that the O-GlcNAcylation of METTL1 does not directly affect the ubiquitination-mediated degradation of METTL1 (Fig. 6J-L). However, after the function of OGT was inhibited with OSMI-1, OSMI-1 inhibited the ubiquitination level of METTL1 under high glucose conditions (Fig. 6M and N). After O-GlcNAcylation level was promoted with Thiamet G, the level of ubiquitinated METTL1 under high glucose conditions significantly increased (Fig. 6O and P). Additionally, our findings indicate that OGT knockdown could suppress the ubiquitination of METTL1 under high-glucose conditions (Fig. 6Q). Therefore, we speculate that since O-GlcNAcylation cannot directly affect the ubiquitination of METTL1 itself, there must be a mediator in this process, and the molecular function of this mediator may be influenced by O-GlcNAcylation.

### T262A Mutation of METTL1 Confers Protection Against Diabetic Cardiomyopathy of ob/ob mice *in vivo* and *in vitro*

To further clarify the role of O-GlcNAcylation of METTL1 modification in pathological changes of diabetic cardiomyopathy. We constructed the AAV9 adeno-associated virus carried with METTL1 (WT) and METTL1 (T26A, the same modification with human) and injected the ob/ob mice. Interestingly, the heart tissue of the METTL1 (T26A) mice was notably smaller than that of the METTL1 (WT) mice of ob/ob background (Fig. 7A and B). Decreased BNP levels were found in the peripheral blood serum of the METTL1 (T26A) mice (Fig. 7C). Histological analyses using HE and Masson staining revealed substantial myocardial hypertrophy and fibrosis in the METTL1 (WT) mice compared with those in the METTL1 (T26A) mice. WGA staining further demonstrated that mutation of the METTL1 site in the myocardium inhibited myocardial hypertrophy in diabetic mice (Fig. 7D). Additionally, qPCR analysis revealed significant decrease in the expression levels of heart hypertrophy-related genes (Nppa, Nppb, and β-MHC) following METTL1 site mutation (Fig. 7E). Moreover, the expression of proinflammatory genes (IL-1β, IL-6 and TNF-α) was significantly downregulated (Fig. 7E). The fatty acid decomposition level in cardiomyocytes was assessed through FAO metabolism. The results indicated that mutation of O-GlcNAcylation of METTL1 in NMCM cells contributed to the fatty acid decomposition triggered by stock (Fig.7F). Additionally, oxygen consumption rate (OCR) analysis revealed that mutation of O-GlcNAcylation of METTL1 increased oxidative phosphorylation in cardiomyocytes stimulated with high glucose (Fig.7G). Western blot analysis revealed significant decreases in the expression levels of hypertrophy-related proteins (ANP, BNP), fibrosis-related proteins (collagen I, α-SMA) and inflammatory factors (NLRP3, IL-1β) (Fig. 7H) in the heart tissue of the METTL1 site mutation mice compared with those of the METTL1 (WT) mice with an ob/ob background. Quantitative PCR analysis revealed that key genes involved in cholesterol synthesis were significantly downregulated in site mutation mice compared with WT mice, whereas the expression of fatty acid β-oxidation-related genes was notably upregulated (Fig. 7I).

**Figure 7.**
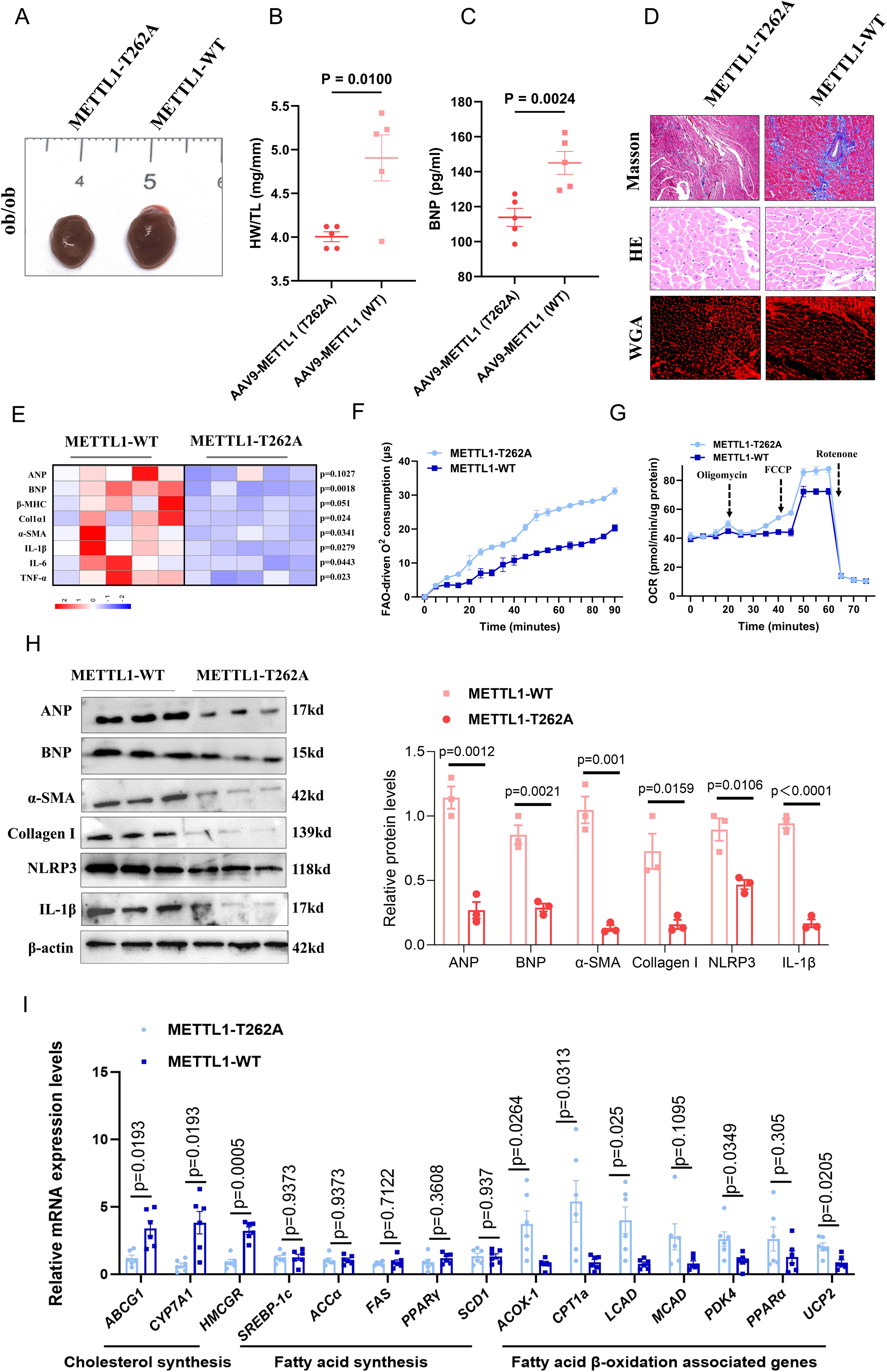
T262A Mutation of METTL1 Confers Protection Against Diabetic Cardiomyopathy of ob/ob mice *in vivo*. 8-week-old ob/ob mice were injected with the adeno-associated virus 9 (AAV9) that carried with the mouse METTL1 (WT) (ob/ob + AAV9-METTL1^WT^) and METTL1 (T262A, mutation site of threonine (T) to alanine (A) at 262 of METTL1 protein) plasmids (ob/ob + AAV9-METTL1^T262A^). The mice were then cultured and sacrificed at age 16 weeks. (A) Representative general photograph of heart tissues. Statistical analysis of ratio of HW/TL (B) and serum BNP levels (C) compared with ob/ob+AAV9-METTL1^WT^ and ob/ob + AAV9-METTL1^T262A^ diabetic mice. Data are expressed as mean ± SEM. (D) Masson, HE and WGA staining of myocardial tissues with ob/ob + AAV9-METTL1^WT^ and ob/ob + AAV9-METTL1^T262A^ diabetic mice. (E) Representative relative mRNA expression levels of genes and heatmap of ANP, BNP, β-MHC, Col1α1, α-SMA, IL-1β, IL-6 and TNF-α were detected by qPCR assay in heart tissues from ob/ob + AAV9-METTL1^WT^ and ob/ob + AAV9-METTL1^T262A^ diabetic mice. NMCM cells were transfected with the METTL1 (WT) and METTL1 (T262A) mutation adenovirus and cultured in HG for 24h. Then, FAO colorimetric assay (F) and Seahorse analysis for OCR (G) for NMCM were conducted. (H) Representative images and statistical analysis of ANP, BNP, α-SMA, Collagen I, NLRP3 and IL-1β were detected by immunoblotting analysis in heart tissues from ob/ob + AAV9-METTL1^WT^ and ob/ob + AAV9-METTL1^T262A^ diabetic mice. (I) Statistical analysis of metabolites of fatty β-oxidation pathway in heart tissues from ob/ob + AAV9-METTL1^WT^ and ob/ob + AAV9-METTL1^T262A^ diabetic mice. Data are expressed as mean ± SEM.

*In vitro* cell assay shown that the expression levels of hypertrophy-related proteins (ANP, BNP), fibrosis-related proteins (collagen I, α-SMA), and inflammatory factors (NLRP3, IL-1β, TNF-α) were inhibited after over-expression level of METTL1(T268A) in AC16 cells and METTL1(T262A) in NMCM cells under high condition (Fig. S15A and B). All these results shown that mutation of O-GlcNAcylation site of METTL1 alleviated the cardiac function of ob/ob mice *in vivo* and *in vitro*.

### High glucose suppresses USP5 deubiquitinase activity to promote METTL1 degradation

These results indicate that high glucose promotes the ubiquitination-mediated degradation of METTL1 in a manner dependent on OGT-mediated glycosylation. To identify this crucial intermediate molecule, we conducted primary screening via ELISAs and secondary screening with western blotting (WB) on HEK293T cells transfected with deubiquitinase siRNA libraries (Fig. S16A and B), as well as mass spectrometric analysis of OGT-bound proteins (Fig. 8B). After library screening and mass spectrometry-based interaction analysis, we discovered that USP5 may regulate METTL1 as a key deubiquitinase (Fig. 8A, S16C and D). The literature also suggests that METTL1, along with the family protein METTL3, is regulated by USP5, leading us to further speculate that the key intermediate molecule induced by high glucose in the ubiquitination of METTL1 could be USP5 (Fig. 8B). Combined the results of deubiquitinase siRNA libraries and OGT-bound proteins, we found that USP5 might be the most potential candidate (Fig. 8B). To validate our hypothesis, we performed immunoprecipitation (Fig. 8C and D, Fig. S17A-C) and fluorescence co-localization experiments, and the results revealed that METTL1 can interact with USP5, with the OGT protein also interacting with USP5 (Fig.8E). Docking of the USP5 and METTL1 proteins using HDOCK revealed that the Asp residue at position 15 and the Ser residue at position 17 of METTL1 can form hydrogen bonds with the Ser residue at position 728 and the Arg residue at position 731 of USP5, with the Tyr residue at position 57 of USP5 forming a hydrogen bond with the Trp residue at position 582 of USP5 and the Met residue at position 112 and the Ile residue at position 146 of USP5 forming hydrogen bonds with the Asn residue at position 230 and the His residue at position 232 of USP5 (Fig. 8F). Analysis of the GEPIA database revealed a positive correlation between the expression levels of USP5 and METTL1 via Spearman analysis (p < 0.0001, R = 0.68) and Pearson analysis (p < 0.0001, R=0.64) (Fig. 8G) and in other organs (Fig. S18A). Furthermore, we analyzed the correlation between USP5 and METTL1 in single-nuclei profiling analysis and the result shown that USP5 was positive correlation in cardiomyocyte (Fig. S19B). Additionally, immunoprecipitation experiments revealed that high glucose can promote O-glycosylation of USP5 and inhibit the interaction between METTL1 and USP5 (Fig. 8H). Immunoprecipitation experiments following mutation of the USP5 binding sites revealed that the binding of METTL1 to USP5 mainly depends on the Asn residue at position 230 and the His residue at position 232 (Fig. 8I and J). Treatment with Thiamet G to activate O-glycosylation of USP5 inhibited the interaction between METTL1 and USP5, as well as the deubiquitination activity of USP5 on METTL1 (Fig. 8K and M, Fig. S17D). We further investigated the deubiquitination activity of USP5 on METTL1 and found that the deubiquitination domain of mutated USP5 significantly weakened its deubiquitination activity towards METTL1 when it was transfected (Fig. 8L). OGT knockdown via siRNA restored the deubiquitination activity of USP5 towards METTL1 after induction by high glucose (Fig. 8N).

**Figure 8:**
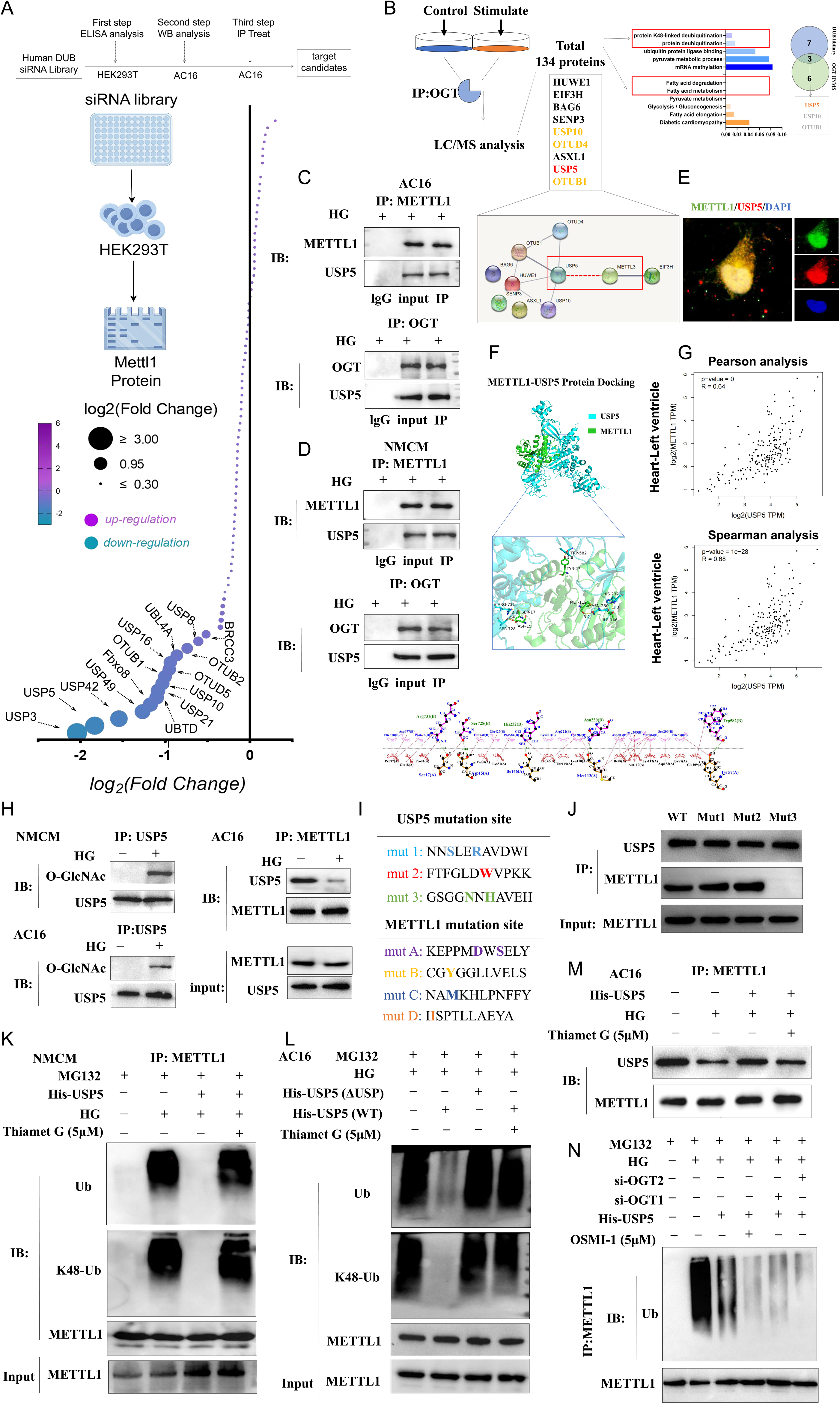
High glucose inhibited the deubiquitination activity of USP5, thus increasing the ubiquitination-mediated degradation of METTL1 via O-GlcNAcylation. (A) HEK293T were transfected with Human DUB siRNA library and collected the cells for ELISA assay and WB assay to detected the protein levels of METTL1 (up). Bubble diagram shows the level of METTL1 expression after different DUB siRNA interference. Fold changes compared with the control (si-NC) group (below). (B) MEFs were cultured and collected for the Co-IP assay and the beads of the assay was detected via mass spectrometry (left). KEGG analysis for the potential proteins that bind with the OGT in MEFs via mass spectrometry analysis (right). (C) AC16 cells were cultured in high glucose for 24h and collected the cells for further Co-IP assay to detect the interaction of USP5 and METTL1 (up). AC16 cells were cultured in high glucose for 24h and collected the cells for further Co-IP assay to detect the interaction of USP5 and OGT (below). (D) NMCM cells were cultured in high glucose for 24h and collected the cells for further Co-IP assay to detect the interaction of USP5 and METTL1 (up). NMCM cells were cultured in high glucose for 24h and collected the cells for further Co-IP assay to detect the interaction of USP5 and OGT (below). (E) AC16 cells were cultured and for the co-location of METTL1 and USP5. DAPI shown the nucleus, green shown the METTL1, red shown the USP5. (F) The structure of METTL1 and USP5 were predicted via AlphaFold 3.0 and for further molecular docking analysis and drawn via PyMoL software. (G) The Spearman analysis and Pearson analysis for the correlation of USP5 and METTL1 in left ventricle of human. Data were download from the GEPIA Database. (H) NMCM and AC16 cells were cultured and treated with HG or not for 24h and cells were collected for further Co-IP assay to detect the O-GlcNAcylation of USP5 (left) and the interaction level of USP5 and METTL1 (right). (I) Design for the mutation of USP5 and METTL1 on the site depended on its own binding site predicted via HDOCK. (J) HEK293T cells were transfected with different mutation site of USP5 and for further Co-IP assay to clarify the potential binding site of USP5 to METTL1. (K) NMCM cells transfected with USP5 over-expression plasmid *in vitro* and cells were cultured under the treatment of HG and Thiamet G (5μM) for 24h and MG132 for 6h. Cells were then collected for further Co-IP assay to detected the *in vivo* ubiquitination of METTL1. (L) AC16 cells transfected with USP5 over-expression plasmid (His-USP5) and USP5 mutation plasmid (lacking the catalytic domain (USP domain). USP5 (ΔUSP)) *in vitro* and cells were cultured under the treatment of HG and Thiamet G (5μM) for 24h and MG132 for 6h. Cells were then collected for further Co-IP assay to detected the *in vivo* ubiquitination of METTL1. (M) AC16 cells were transfected with the His Tag Usp5 plasmid and cultured under the treatment of HG and Thiamet G (5μM) for 24h. Cells were collected for further Co-IP assay to detect the interaction level of USP5 and METTL1. (N) AC16 cells were transfected with the His Tag Usp5 plasmid and two independent small interfering RNAs targeting the OGT. Cells were then cultured under the treatment of HG and Thiamet G (5μM) for 24h and collected for Co-IP assay to detected the *in vivo* ubiquitination of METTL1.

### Small molecule HIT106265621 disrupts OGT-METTL1 interaction and attenuates cardiac pathology in diabetic mice

Given that our above results showed that OGT regulates the O-GlcNAcylation of METTL1, we investigated whether interfering with the OGT-METTL1 interaction could alleviate the disease progression of DCM. The potential binding sites of OGT and METTL1 were subsequently identified by MD simulations. The small molecule compound library in the Chemdiv database was used for computer-aided drug screening (Fig. 9A). Structural visualization of the final MD frame (Fig. 9B) revealed three critical interaction networks: Region C: OGT Asn124-Glu90-Gln33 ↔ METTL1 Glu23-Arg24-His165; Region D: OGT Asn230-Asp221-Asp162-Leu254 ↔ METTL1 Pro221-Arg246-Arg19-Glu93; and Region 3: OGT Arg6-Asn7-Val8 ↔ METTL1 Asn332-Asn301-Asn298. By predicting the presence of a small-molecule binding site (pink) on OGT, we found that this site is important for OGT-METTL1 interaction; thus, virtual screening of this site can yield inhibitors that interfere with the interaction of OGT and METTL1. Through computational virtual screening, we identified 30 small-molecule candidates targeting the OGT-METTL1 interaction interface (Top 5 were shown in Fig. S19). To evaluate their inhibitory efficacy, we developed a luciferase-based protein-protein interaction (PPI) reporter system in HEK293T cells, where the luciferase signal intensity is correlated with the OGT-METTL1 binding status. All the compounds were procured and tested at an initial concentration of 10μM. Quantitative luciferase assays performed 48 hours post-treatment revealed dose-dependent inhibition across three concentrations (10, 20, and 30 μM). Notably, HIT106265621 significantly suppressed the OGT-METTL1 interaction (p < 0.001 vs. vehicle control), maintaining > 50% inhibition even at the lowest concentration (10 μM), outperforming the other candidates in both potency and efficacy (Fig. S20A-C). To further analyse the MD of OGT-HIT106265621 complex stability (Fig. 9C-E), after 100-ns MD simulations, we conducted comprehensive trajectory analyses. The RMSD of both the protein and ligand stabilized below 2.0 Å after 20 ns (Fig. S21A), indicating system equilibration. Subsequent analyses focused on the 20-100-ns trajectory window, with 100 representative conformations extracted at 1ns intervals for structural alignment (Fig. S21B). Fig. S21C shows excellent conformational convergence (average Cα RMSD < 1.5 Å), confirming the maintenance of structural integrity and stable ligand binding within the active site (Fig. S21D). RMSF and B factor analyses revealed low structural flexibility (average RMSF < 0.25 nm) across the protein (Fig. S21D), particularly at the binding pocket (residues 29-94). This rigidity profile suggests strong ligand-induced stabilization, with binding site fluctuations 40% lower than the global protein average. Detailed interaction analysis identified five key interfacial residues: His29, Leu59, Phe66, Glu90, and Asn94 (Fig. 9E and Fig. S21E). These residues maintain persistent contacts (> 80% occupancy) through hydrophobic interactions (Leu59/Phe66), water-mediated hydrogen bonds (Glu90/Asn94) and π-π stacking (His29) (Fig. S21F). Further quantitative binding metrics revealed the following: stable hydrogen bond count: 3.2 ± 0.4 (mean ± SD); ligand positional RMSD: 1.1 ± 0.3 Å; and binding pocket volume conservation: 92.4% (Fig. S21F). Energy decomposition analysis revealed that Glu90 and Asn94 contributed 68% of the total polar interaction energy (-12.3 kcal/mol combined), whereas hydrophobic residues (His29/Leu59/Phe66) accounted for 85% of the nonpolar contributions (-9.8 kcal/mol) (Fig. S21E).

**Figure 9.**
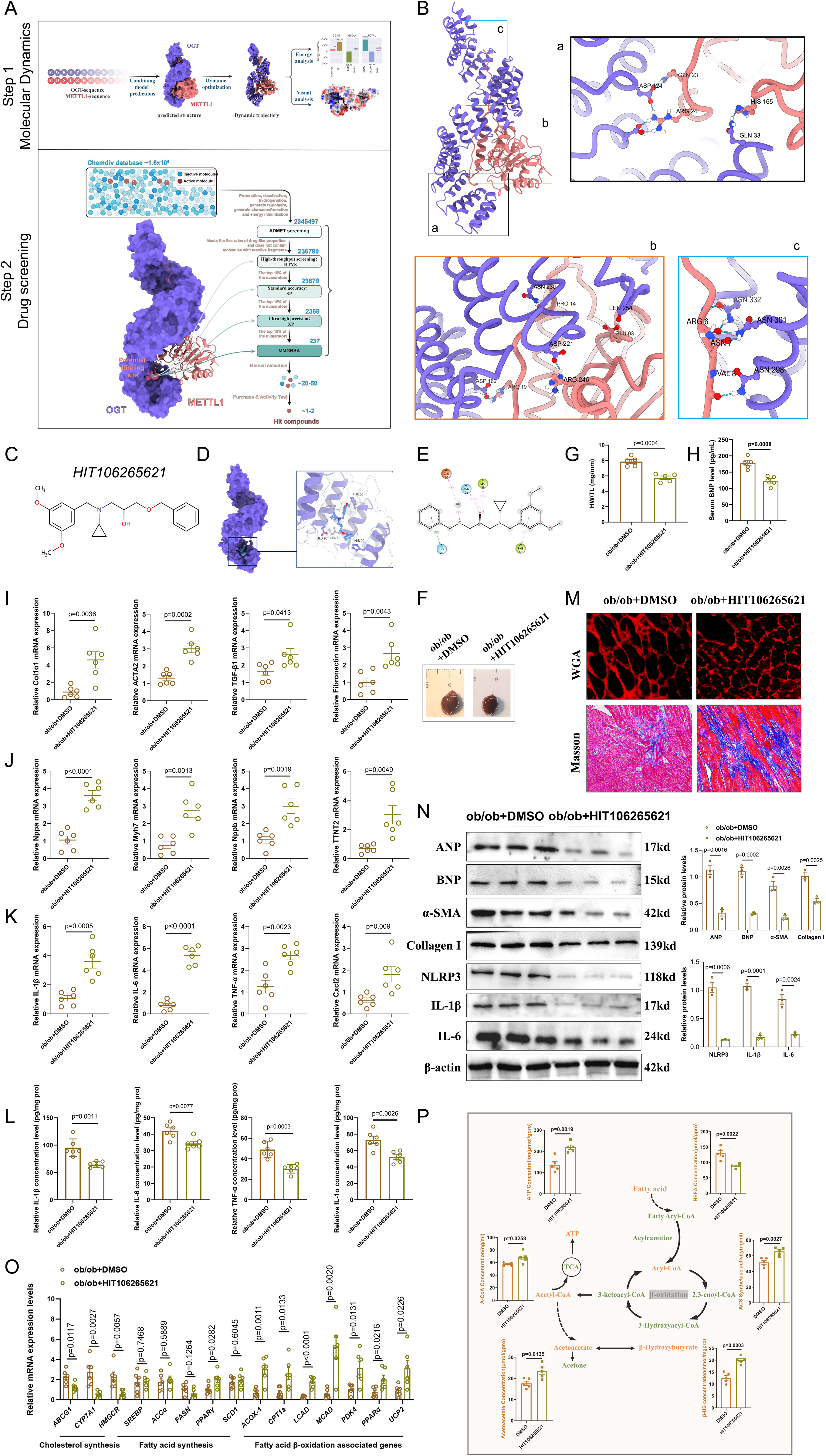
HIT106265621 might be a potential drug to alleviate pathological changes associated with spontaneous ob/ob DCM by targeting the interaction of OGT and METTL1. (A) Working procedures for the screening potential small molecule drug interfering interaction of METTL1 and OGT. (B) Molecular docking and molecular dynamics simulations analysis of OGT and METTL1 (left). The potential binding area and site of OGT and METTL1 (right). (C) Structure of HIT106265621. (D) Molecular docking and molecular dynamics simulations analysis of HIT106265621 and OGT. (E) Potential binding area and site of HIT106265621 and OGT. 8-week-old ob/ob mice were treated with HIT106265621 (2.5mg/kg in DMSO, i.p., twice per week) for 8 weeks and the mice were then sacrifice at age 16 weeks. Representative general photograph of heart tissues (F), statistical analysis of ratio of HW/TL (G) and serum BNP levels (H) compared among ob/ob + DMSO and ob/ob + HIT106265621 mice. Data are expressed as mean ± SEM. (I) Comparison of relative mRNA expression levels of genes associated with cardiac fibrosis (Col1α1, Acta2, TGF-β1 and fibronectin) among ob/ob + DMSO and ob/ob + HIT106265621 mice. (J) Comparison of relative mRNA expression levels of genes associated with cardiac hypertrophy (Nppa, Myh7, Nppb and TINT2) among ob/ob + DMSO and ob/ob + HIT106265621 mice. (K) Comparison of relative mRNA expression levels of genes associated with pro-inflammatory cytokines (IL-1β, IL-6, TNF-α and Cxcl2) among ob/ob + DMSO and ob/ob + HIT106265621 mice. (L) Relative concentration levels of pro-inflammation factors including IL-1β, IL-6, TNF-α and IL-1α in heart tissues among ob/ob + DMSO and ob/ob + HIT106265621 mice via ELISA assay. Data are expressed as mean ± SEM. (M) WGA and Masson staining of myocardial tissues among ob/ob + DMSO and ob/ob + HIT106265621 mice. (N) Representative images and statistical analysis of ANP, BNP, α-SMA, Collagen I, NLRP3, IL-1β and IL-6 were detected by immunoblotting analysis in heart tissues from ob/ob + DMSO and ob/ob + HIT106265621 mice. (O) Relative expression levels of genes associated with cholesterol metabolism, fatty acid synthesis and fatty acid β-oxidation pathways in heart tissues from ob/ob + DMSO and ob/ob + HIT106265621 mice. (P) Statistical analysis of metabolites of fatty β-oxidation pathway in heart tissues from ob/ob + DMSO and ob/ob + HIT106265621 mice. Data are expressed as mean ± SEM.

**Figure 10.**
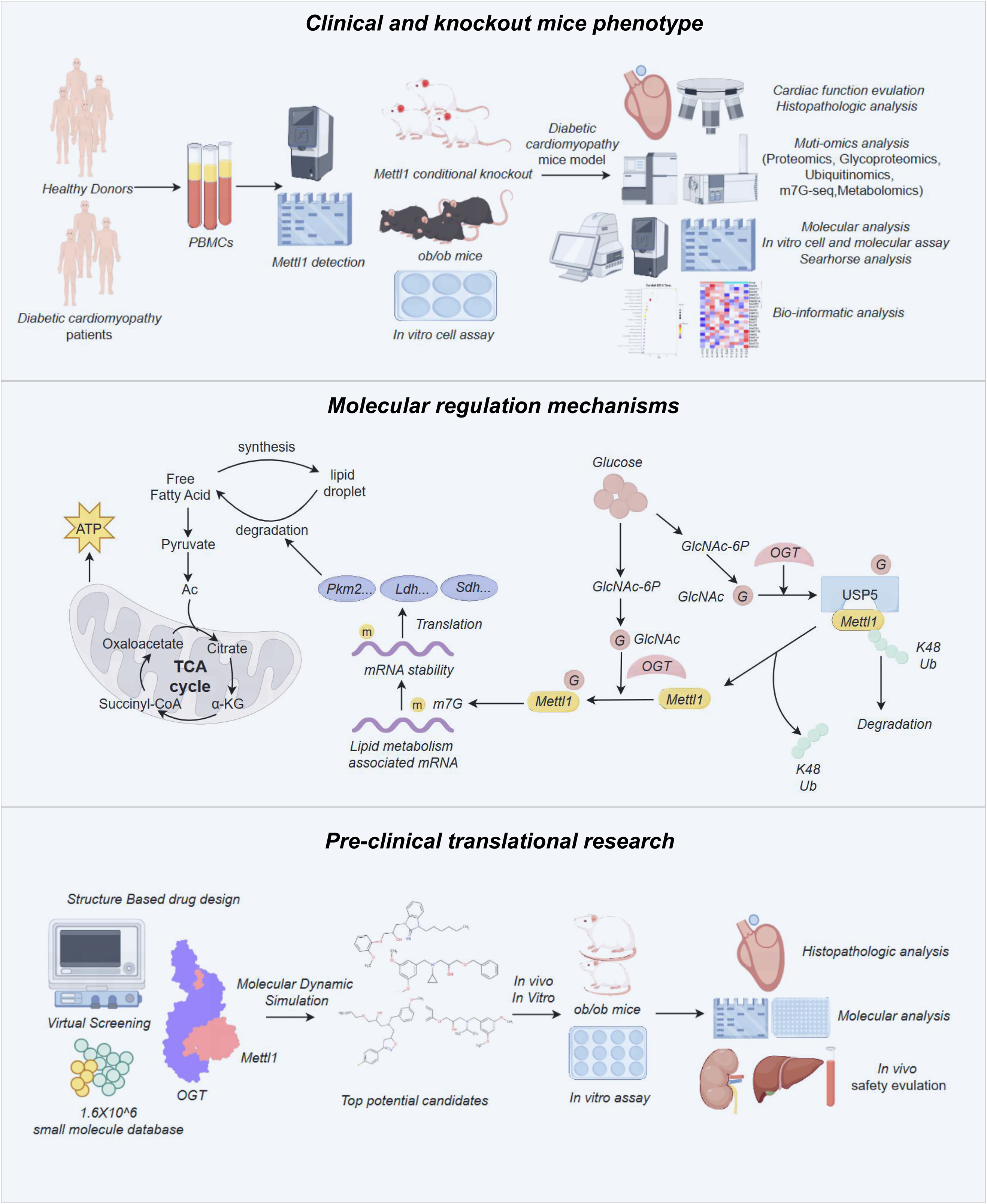
Multiomic analysis and preclinical drug development targeting METTL1 in DCM. This study investigates the role of METTL1 in diabetic cardiomyopathy using clinical samples, knockout mouse models and *in vitro* assays. The research comprises three key sections: (1) Clinical and knockout mice phenotype, where METTL1 expression is analysed in diabetic patients and METTL1 knockout mice, followed by cardiac function assessments, multiomic analysis, and bioinformatics approaches. (2) Molecular regulation mechanism, elucidating METTL1’s role in lipid metabolism, glucose regulation, and protein stability via ubiquitination and glycosylation pathways. (3) Pre-clinical translational research, involving structure-based drug design, molecular simulations, and *in vivo/in vitro* validation of potential METTL1-targeting compounds. This comprehensive approach aims to identify novel therapeutic strategies for diabetic cardiomyopathy.

Subsequent investigations revealed that HIT106265621 relieved heart enlargement induced in ob/ob diabetic mice (Fig. 9F and G) and BNP level of ob/ob mice (Fig. 9H). Additionally, qPCR analysis revealed significant decreases in the expression levels of fibrosis-related genes (Col1α1, Acta2, TGF-β1, and fibronectin) following HIT106265621 treatment (Fig. 9I). Moreover, the expression of heart hypertrophy-related genes (Nppa, Myh7, Nppb, and TTNT2) (Fig. 9J) and proinflammatory genes (IL-1β, IL-6, TNF-α, and Cxcl2) (Fig. 9K) and was significantly downregulated. ELISA results for IL-1β, IL-6, TNF-α, and IL-1α in heart tissue further supported the notion that HIT106265621 could inhibit the proinflammatory response in the heart (Fig. 9L). Histological analyses using Masson staining and WGA staining demonstrated that HIT106265621 alleviated hypertrophy of myocardial cells and severe fibrotic changes (Fig. 9M). Western blot analysis revealed significant decreases in the expression levels of hypertrophic proteins (ANP and BNP), fibrosis-related proteins (collagen I, α-SMA) and inflammatory factors (IL-1β, IL-6, and NLRP3) in the heart tissue of the HIT106265621-treated mice compared with those of the DMSO mice (Fig. 9N).

To investigate the effects of HIT106265621 on fatty acid metabolism in DCM, we performed qPCR analysis, which revealed that key genes involved in cholesterol synthesis were significantly downregulated in the HIT106265621-treated mice compared with the DMSO-treated mice, whereas the expression of fatty acid β-oxidation-related genes was notably upregulated. However, we found no significant differences in the key genes related to fatty acid synthesis (Fig. 9O). Furthermore, in heart tissues from the HIT106265621-treated mice and the DMSO-treated mice, the HIT106265621-treated mice presented significantly lower levels of free fatty acids in cardiomyocytes, along with increased ACS synthetase activity and β-hydroxybutyric acid, ethyl acetate, acetyl-CoA, and ATP levels (Fig. 9P). These results suggest that HIT106265621 can alleviate myocardial injury and inflammation in diabetic mice and promote fatty acid metabolism and oxidative phosphorylation in cardiomyocytes.

To further confirm the in vivo safety of HIT106265621 in mice, 8-week-old C57BL/6 mice were treated with HIT106265621 (2.5 mg/kg in DMSO, i.p., twice per week) until 24 weeks old. We then collected the serum from mice and detected the levels of BUN and AST to reflect the function of liver and kidney *in vivo*. We found that HIT106265621 treatment had no significant changes in the levels of AST, ALT, BUN and Creatinine in serum (Fig. S22A-D). Also, the HIT106265621 treatment had no significant changes on the number of White Blood Cells and level of Hemoglobin *in vivo* (Fig. S22E-G). All these results indicated the *in vivo* safety of HIT106265621 treatment. In conclusion, HIT106265621 might be potential candidate for alleviating pathological changes in ob/ob mice with DCM by targeting the interaction of OGT and METTL1.

## Discussion

Metabolic memory is the key mechanism that causes damage to diabetic target organs. This pathological process is closely related to RNA methylation, DNA methylation and histone modification(H3K9ac^48,49^, H3K4me3^50^, H3K27me3^51^, etc.) are involved in generating diabetic metabolic memory. The role of RNA methylation in diabetic metabolic memory has gradually been recognized in recent years, although many types of RNA modifications have been identified. In this research, we unravel a previously unrecognized post-translational regulatory circuit linking hyperglycaemia to METTL1 destabilization. High glucose induced T268-specific O-GlcNAcylation of METTL1, and facilitated K48-linked polyubiquitination and proteasomal degradation independent of O-GlcNAcylation. Strikingly, this process required USP5 as a critical mediator-glycoproteomics revealed OGT-USP5 interaction, creating a “double-hit” mechanism: O-GlcNAcylation directly impaired METTL1 methyltransferase activity (m7G-RNA, while USP5 inactivation blocked METTL1 stabilization. This dual regulation explains the persistent METTL1 depletion observed in chronic hyperglycemia, establishing a feed forward loop sustaining metabolic memory.

Via two classical diabetic murine models (STZ+HFD and ob/ob mice) and samples from DCM patients, METTL1 downregulation correlated strongly with cardiac dysfunction. Cardiomyocyte-specific METTL1 knockout (METTL1^fl/fl^Myh6^CreERT^) exacerbated diastolic dysfunction and metabolic remodeling, evidenced by long-chain acylcarnitine accumulation and impaired β-oxidation. Crucially, METTL1 deficiency disrupted m7G epitranscriptomic regulation of fatty acid metabolism enzymes, particularly targeting Scd1, Aldoa, Acadm, etc., which collectively drive lipid droplet accumulation. These findings position METTL1-mediated m7G modification as a central gatekeeper of cardiomyocyte metabolic homeostasis under diabetic stress. Also, we showed that high glucose can induce O-GlcNAcylation of USP5, inhibit its interaction with METTL1 and deubiquitination, and promote the ubiquitination-mediated degradation of METTL1. Moreover, tyrosine 268 of METTL1 itself is modified by O-GlcNAcylation, which inhibits the functional activity of m7G modification. The discovery of *HIT106265621* as an OGT-METTL1 interaction inhibitor provides proof-of-concept for targeting this axis. In ob/ob mice, HIT106265621 reversed cardiac hypertrophy, fibrosis, and restored fatty acid oxidation. Clinically, PBMC METTL1 levels showed strong prognostic value for DCM progression, with patients in the lowest METTL1 quartile exhibiting higher risk of heart failure hospitalization.

m7G modification mainly occurs in the 5’ CAP, 5’UTR, and A-G-rich regions of mRNAs^52^. The methyltransferase complex RNMT/RAM catalyses the addition of methyl groups from S-adenosyl methionine to the N7 position of guanine nucleotides, forming the ’cap’ structure of m5G (5’)ppp(7’)X^53,54^. METTL1 and Wdr4 are believed to be key enzymes involved in m7G modification of tRNA, mRNA, and miRNA^55^. The METTL1/WDR4 complex introduces the m7G cap into mRNA molecules, protecting them from degradation and increasing their translation efficiency. METTL1 protein-mediated m7G modification plays important regulatory roles in tumours^56^, cardiac hypertrophy^38,57^, angiogenesis^33,58^, metabolic reprograming^59,60^, and cellular senescence^61^. In our study, we found that METTL1 knockout significantly promoted myocardial hypertrophy and reduced cardiac function in diabetic mice. We attributed this process to its role in remodelling fatty acid metabolism in cardiomyocytes. However, METTL1 has also been reported in previous studies to promote cardiac hypertrophy. Unlike our study, Yu et al. reported that, after intervention with transverse aortic constriction (TAC) and angiotensin II (Ang II) in METTL1 heterozygous mice, METTL1 semi-deletion decreases SRSF9 expression by inducing m7G modification of SRSF9 mRNA, thereby facilitating alternative splicing and stabilizing NFATc4 to inhibit cardiac hypertrophy^36^. While some studies have also demonstrated the adverse effects of METTL1 knockout in fibroblasts on post-myocardial infarction fibrosis^37^. These diverse findings suggest that the role of METTL1 proteins in different diseases is complex. While research into the role of the METTL1 protein in cardiovascular disease is still in its early stages, further exploration is needed to better understand its involvement. Current evidence on the impact of METTL1 remains limited. More studies are necessary to elucidate the molecular pathways connecting this protein to heart conditions. Investigating the precise functions and interactions of METTL1 could provide novel insights into the pathogenesis and potential treatment of cardiovascular problems.

GlcNAcylation, a post-translational modification involving the addition of sugar moieties to serine or threonine residues on proteins, plays a critical role in various cellular processes. Altered O-GlcNAcylation of myofilament proteins can disrupt cardiac contractility, contributing to the diminished heart function observed in heart failure (HF)^62,63^. A previous study indicated that the overexpression of O-GlcNAcylation promoted dilated cardiomyopathy and cardiac hypertrophy^43,44,62^. Studies on the relationship between O-GlcNAcylation and heart failure have identified additional potential therapeutic targets, such as HDAC4^44^, PRMT5^62^, CaMKII^64^, and PKA^65^. The mechanism regulating heart failure development through O-GlcNAcylation modulation is relatively complex. This mechanism involves processes such as myocardial inflammation, cell metabolism, autophagic regulation, cytoskeletal protein function regulation, ion channel regulation, and protein modification modulation. YTHDF2, a key enzyme involved in RNA methylation^66^, has been reported to undergo O-GlcNAcylation^40,67^. This O-GlcNAcylation modification of YTHDF2 promotes its protein stability and oncogenic activity by inhibiting its ubiquitination^67^. We reported that METTL1 can also be modified by O-GlcNAcylation and that its O-GlcNAcylation can affect the m7G modification activity of the METTL1 protein. However, we found that the O-GlcNAcylation of METTL1 did not directly affect its ubiquitination-mediated degradation. Furthermore, through small interference library screening and mass spectrometric identification, we determined that the ubiquitination-mediated degradation of METTL1 can be promoted by USP5 O-GlcNAcylation induced by high glucose.

While global METTL1 deletion during embryogenesis causes developmental cardiomyopathy^38^, our inducible cardiomyocyte-specific knockout model reveals a distinct, metabolism-sensitive regulatory paradigm. The differential outcomes suggest the following: 1) Embryonic METTL1 governs fundamental cardiac morphogenesis via bulk translation control, whereas adult METTL1 fine-tunes metabolic enzyme specificity under nutrient stress through O-GlcNAcylation-coupled epitranscriptomic selection. Our work reveals its distinct, previously unrecognized function in acquired metabolic cardiomyopathy through diabetes-specific post-translational regulation. This finding represents a paradigm shift from developmental biology to dynamic epitranscriptomic adaptation. This dichotomy highlights the critical importance of spatiotemporal context in RNA modification biology.

Although we identified the key mechanism and target of O-GlcNAcylation modification and ubiquitination degradation of the METTL1 protein under high-glucose conditions, our study has several limitations. As previously reported, the role of METTL1 in cardiovascular disease is complex^36,38^, and it remains unclear whether a semi-absent state of METTL1 still has a pro-DCM effect. Moreover, the results of O-GlcNAcylation proteomics revealed that the targets of O-GlcNAcylation exist widely in cardiomyocytes, especially the regulatory proteins involved in glucose metabolism and skeletal proteins, and the relationships between these proteins and METTL1 also involve complex regulatory networks. All of these issues require further exploration.

In summary, our study revealed that hyperglycaemia promoted the O-GlcNAcylation of METTL1, inhibiting the deubiquitination of METTL1 by USP5 and thus reducing the level of the METTL1 protein in cardiomyocytes and the m7G modification. Loss of METTL1 reshaped fatty acid metabolism in cardiomyocytes and exacerbated the inflammatory response and myocardial hypertrophy in DCM. Our study aimed to elucidate the intimate connection between RNA methylation and metabolic reprogramming. METTL1 and USP5 may be new targets for the clinical treatment of DCM.

## Consent for publication

Not applicable.

## Availability of data and materials

All data and materials used in the analysis are available to any researcher for purposes of reproducing or extending the analysis from corresponding authors.

## Authors’ contributions

Conceptualization, X.G., H.M., J.G., Y.D., F.Y., and X.W.; Methodology, X.G., J.L., Y.L., B.L., F.W., Q.L., Z.Z., J.L., X.Z., J.S., Y.D., J.L., F.L., X.W., G.H., S.P., X.H., R.Z., J.Z., L.L., and J.Y.; Software, X.G., H.M., J.L., Y.L., W.X., Y.D., J. Z. and F.Y.; Validation, X.G., H.M., J.L., Y.L., B.L., X.W., Y.D., J.G., F.Y., F.W., and J.Z.; Data Analysis, X.G., B.L., F.W., Q.L., Z.Z., Y.D., W.X., X.Z., Y.L., J.Y., F.Y. and J.G.; Writing-Original Draft, X.G. and Y.D., with help from the other authors; Writing-Review & Editing, X.W., F.Y., J.G. and Y.D., with help from the other authors; Project Administration and Supervision, X.G., Z.Z., B.L. and X.W.; Funding Acquisition, Y.D., X.G., H.M., Y.L., J.L, X.W., F.Y., X.Z. and F.W..

## Acknowledgement

The figure was created using FigDraw scientific illustration platform (www.figdraw.com) with official authorization (License ID: PRAWP8848a; OIARIe744b; ORYRT9ddc2; APWWPa7b74).

