## Supplemental FigureS1-S22 and Materials for "O-GlcNAcylation-Ubiquitin Crosstalk of METTL1 Drives m7G Epitranscriptomic Collapse and Lipid Metabolic Reprogramming in Diabetic Cardiomyopathy"

Figure S1

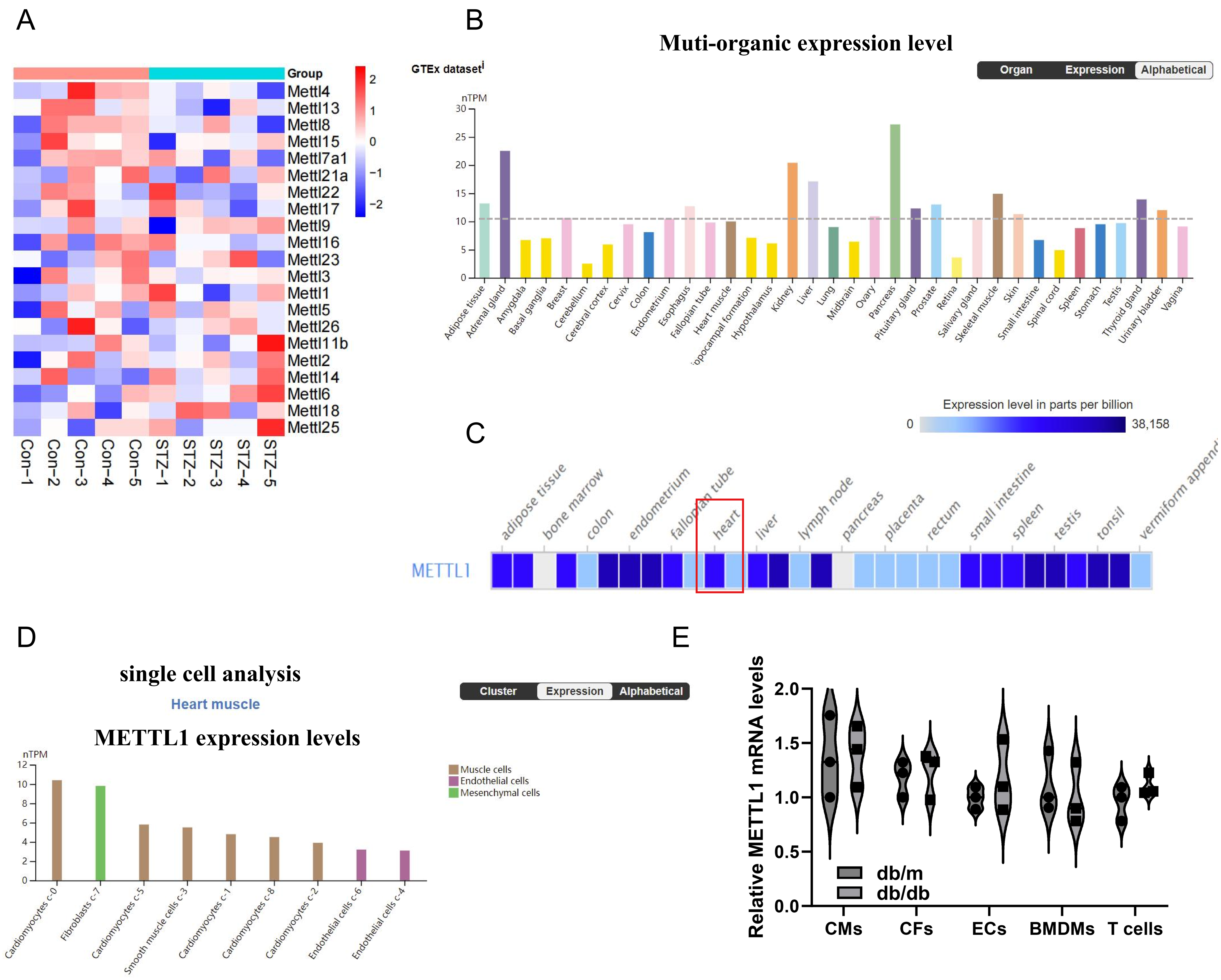

A
Group
Mettl4
Mettl13
Mettl8
Mettl15
Mettl7a1
Mettl21a
Mettl22
Mettl17
Mettl9
Mettl16
Mettl23
Mettl3
Mettl1
Mettl5
Mettl26
Mettl11b
Mettl2
Mettl14
Mettl6
Mettl18
Mettl25
Con−1
Con−2
Con−3
Con−4
Con−5
STZ−1
STZ−2
STZ−3
STZ−4
STZ−5
B
Muti-organic expression level
GTEx dataset
Organ
Expression
Alphabetical
nTPM
Adipose tissue
Adrenal gland
Amygdala
Basal ganglia
Breast
Cerebellum
Cerebral cortex
Cervix
Colon
Endometrium
Esophagus
Fallopian tube
Heart muscle
Hypothalamus
Kidney
Liver
Lung
Midbrain
Ovary
Pancreas
Pituitary gland
Prostate
Retina
Salivary gland
Skeletal muscle
Skin
Small intestine
Spinal cord
Spleen
Stomach
Testis
Thyroid gland
Urinary bladder
Vagina
C
Expression level in parts per billion
0
38,158
adipose tissue
bone marrow
colon
endometrium
fallopian tube
heart
liver
lymph node
pancreas
placenta
rectum
small intestine
spleen
testis
tonsil
METTL1
D
single cell analysis
Heart muscle
METTL1 expression levels
Cluster
Expression
Alphabetical
nTPM
Muscle cells
Endothelial cells
Mesenchymal cells
Cardiomyocytes c-0
Fibroblasts c-7
Cardiomyocytes c-5
Smooth muscle cells c-3
Cardiomyocytes c-1
Cardiomyocytes c-8
Cardiomyocytes c-2
Endothelial cells c-6
Endothelial cells c-4
E
Relative METTL1 mRNA levels
db/m
db/db
CMs
CFs
ECs
BMDMs
T cells

Figure S2

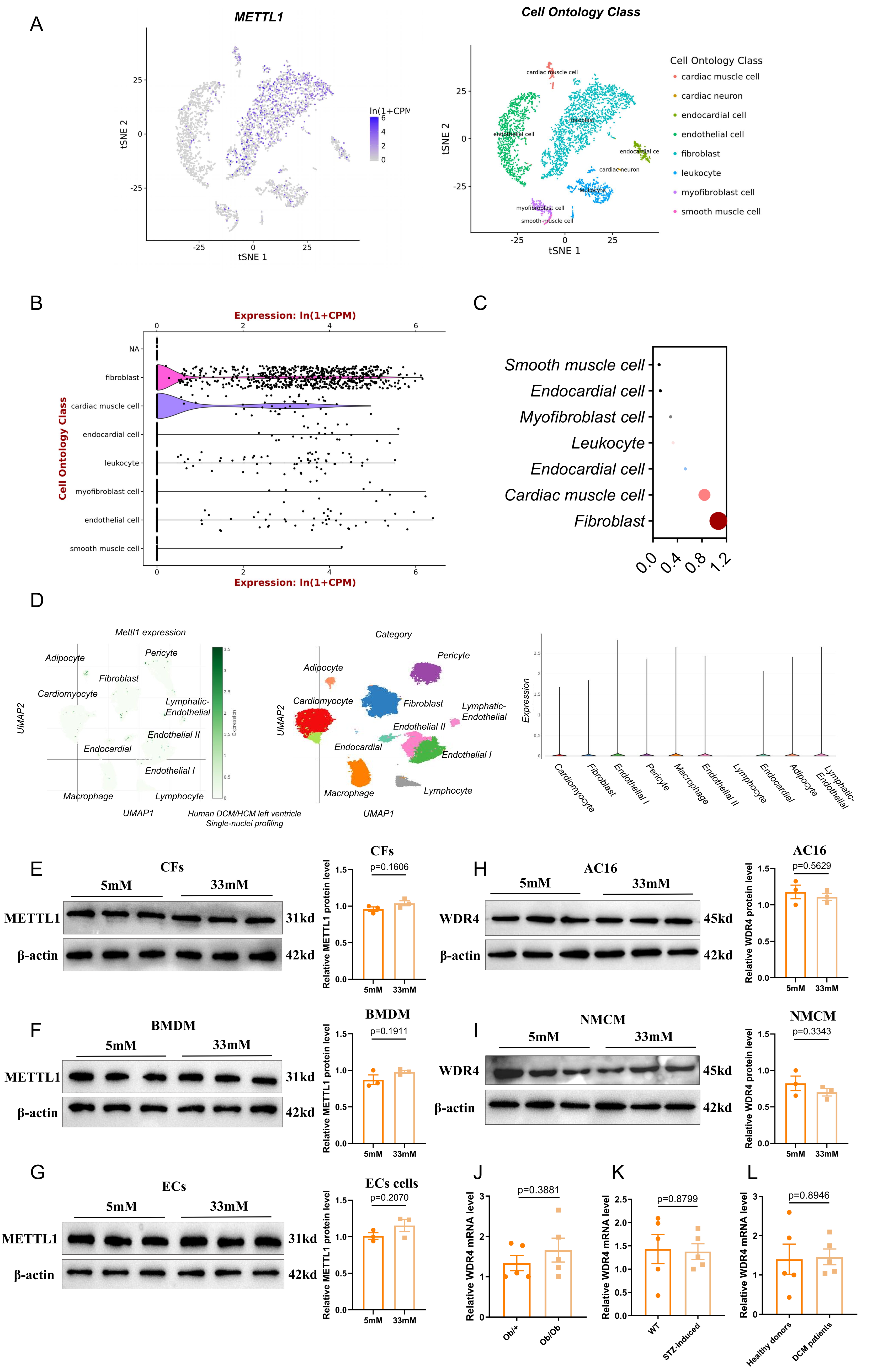

A
METTL1
Cell Ontology Class
tSNE 1
tSNE 2
ln(1+CPM
Cell Ontology Class
cardiac muscle cell
cardiac neuron
endocardial cell
endothelial cell
fibroblast
leukocyte
myofibroblast cell
smooth muscle cell
B
Expression: ln(1+CPM)
Cell Ontology Class
NA
fibroblast
cardiac muscle cell
endocardial cell
leukocyte
myofibroblast cell
endothelial cell
smooth muscle cell
C
Smooth muscle cell
Endocardial cell
Myofibroblast cell
Leukocyte
Endocardial cell
Cardiac muscle cell
Fibroblast
D
Mettl1 expression
Category
Adipocyte
Pericyte
Fibroblast
Cardiomyocyte
Lymphatic-Endothelial
Endothelial II
Endocardial
Endothelial I
Macrophage
Lymphocyte
UMAP1
UMAP2
Human DCM/HCM left ventricle
Single-nuclei profiling
Expression
Cardiomyocyte
Fibroblast
Endothelial I
Pericyte
Macrophage
Endothelial II
Lymphocyte
Endocardial
Adipocyte
Lymphatic-Endothelial
E
CFs
5mM
33mM
METTL1
β-actin
31kd
42kd
Relative METTL1 protein level
p=0.1606
F
BMDM
5mM
33mM
METTL1
β-actin
31kd
42kd
Relative METTL1 protein level
p=0.1911
G
ECs
ECs cells
5mM
33mM
METTL1
β-actin
31kd
42kd
Relative METTL1 protein level
p=0.2070
H
AC16
5mM
33mM
WDR4
β-actin
45kd
42kd
Relative WDR4 protein level
p=0.5629
I
NMCM
5mM
33mM
WDR4
β-actin
45kd
42kd
Relative WDR4 protein level
p=0.3343
J
Relative WDR4 mRNA level
p=0.3881
Ob/+
Ob/Ob
K
Relative WDR4 mRNA level
p=0.8799
WT
STZ-induced
L
Relative WDR4 mRNA level
p=0.8946
Healthy donors
DCM patients

Figure S3

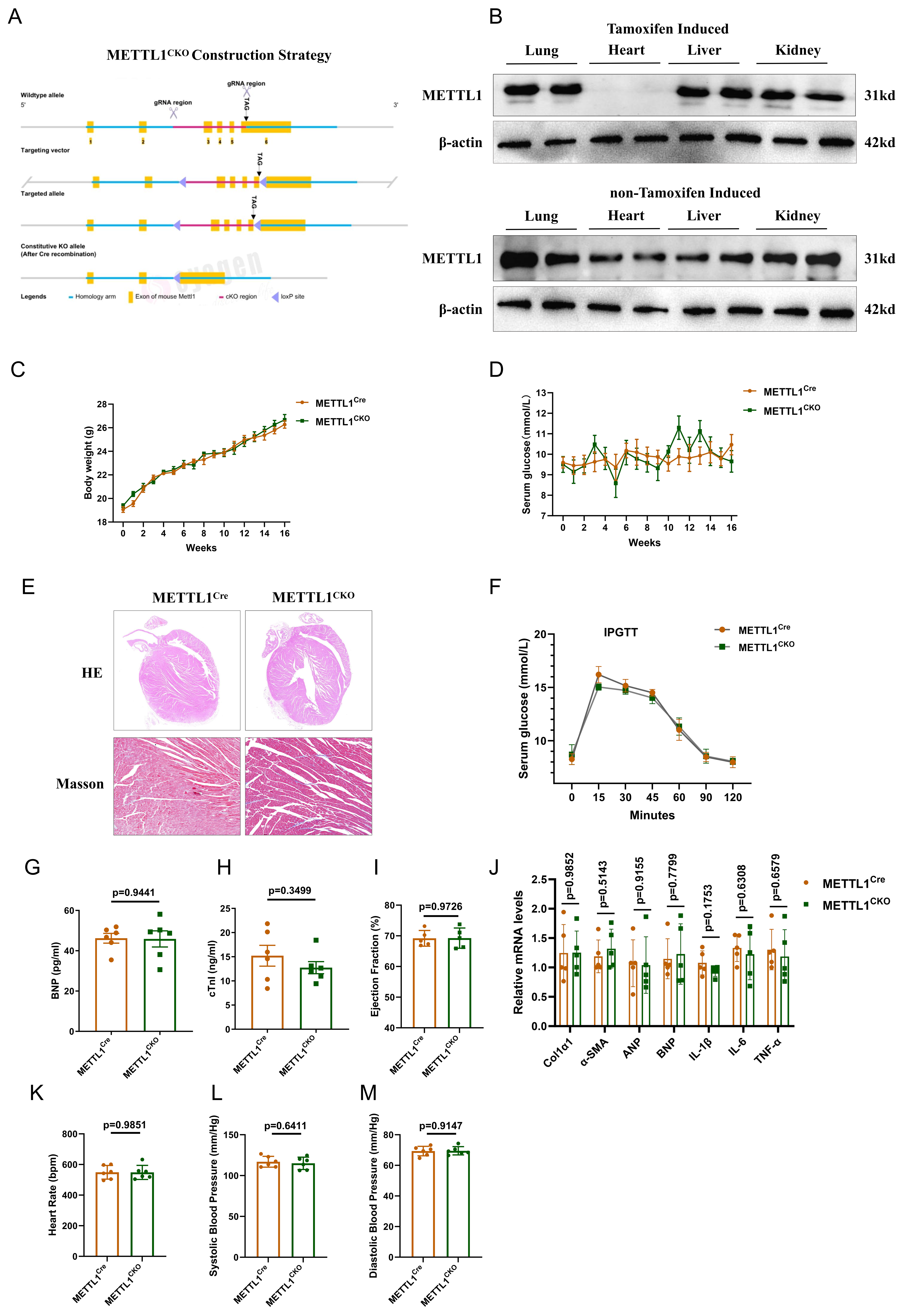

A
METTL1CKO Construction Strategy
Wildtype allele
gRNA region
gRNA region
TAG
Targeting vector
Targeted allele
Constitutive KO allele
(After Cre recombination)
Legends
Homology arm
Exon of mouse Mettl1
cKO region
loxP site
B
Tamoxifen Induced
Lung
Heart
Liver
Kidney
METTL1
β-actin
31kd
42kd
non-Tamoxifen Induced
C
Body weight (g)
Weeks
METTL1Cre
METTL1CKO
D
Serum glucose (mmol/L)
Weeks
E
HE
Masson
F
IPGTT
Serum glucose (mmol/L)
Minutes
G
BNP (pg/ml)
p=0.9441
H
cTnI (ng/ml)
p=0.3499
I
Ejection Fraction (%)
p=0.9726
J
Relative mRNA levels
p=0.9852
p=0.5143
p=0.9155
p=0.7799
p=0.1753
p=0.6308
p=0.6579
Col1α1
α-SMA
ANP
BNP
IL-1β
IL-6
TNF-α
K
Heart Rate (bpm)
p=0.9851
L
Systolic Blood Pressure (mm/Hg)
p=0.6411
M
Diastolic Blood Pressure (mm/Hg)
p=0.9147

Figure S4

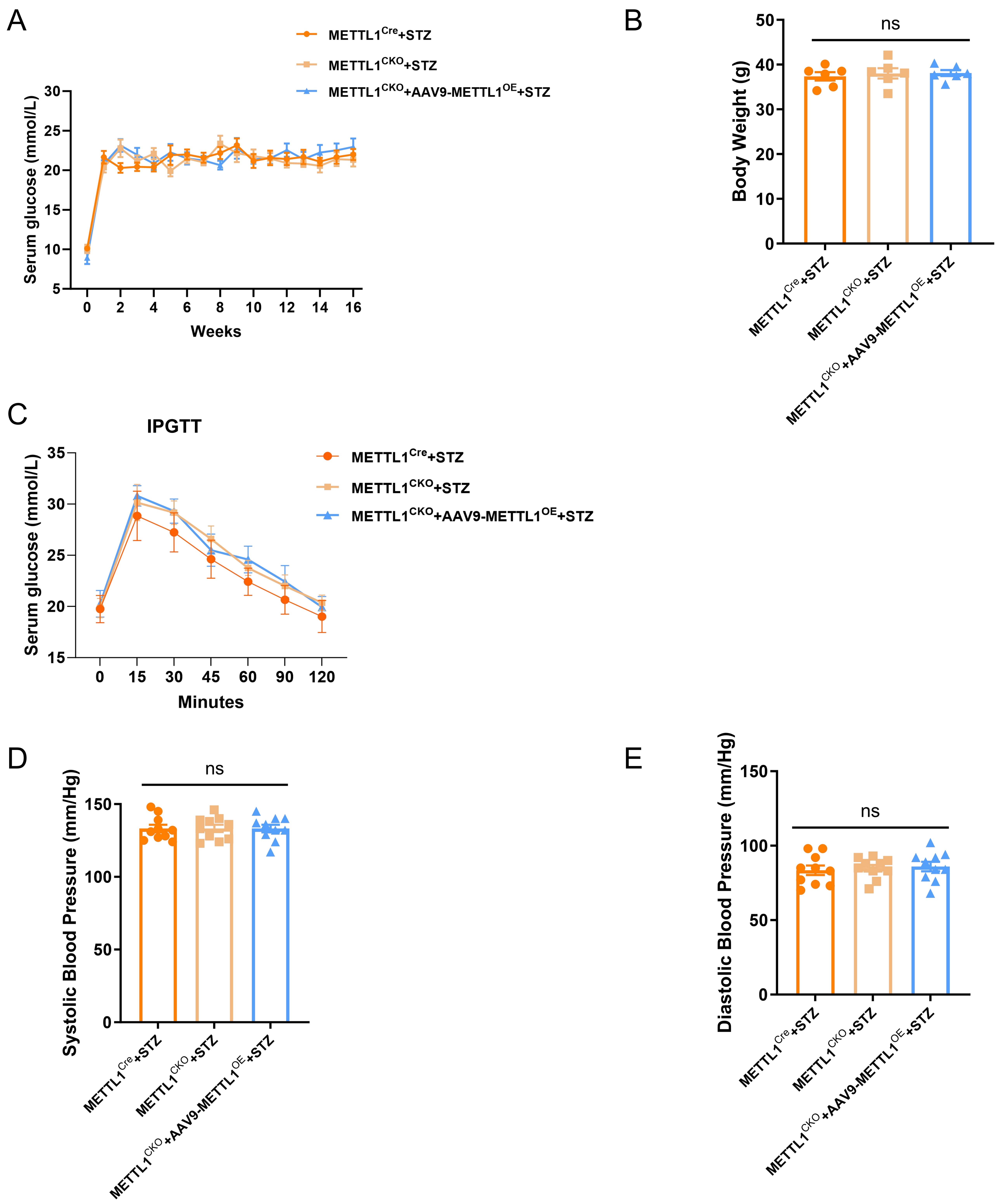

A
METTL1Cre+STZ
METTL1CKO+STZ
METTL1CKO+AAV9-METTL1OE+STZ
Serum glucose (mmol/L)
Weeks
B
ns
Body Weight (g)
METTL1Cre+STZ
METTL1CKO+STZ
METTL1CKO+AAV9-METTL1OE+STZ
C
IPGTT
METTL1Cre+STZ
METTL1CKO+STZ
METTL1CKO+AAV9-METTL1OE+STZ
Serum glucose (mmol/L)
Minutes
D
ns
Systolic Blood Pressure (mm/Hg)
METTL1Cre+STZ
METTL1CKO+STZ
METTL1CKO+AAV9-METTL1OE+STZ
E
ns
Diastolic Blood Pressure (mm/Hg)
METTL1Cre+STZ
METTL1CKO+STZ
METTL1CKO+AAV9-METTL1OE+STZ

Figure S5

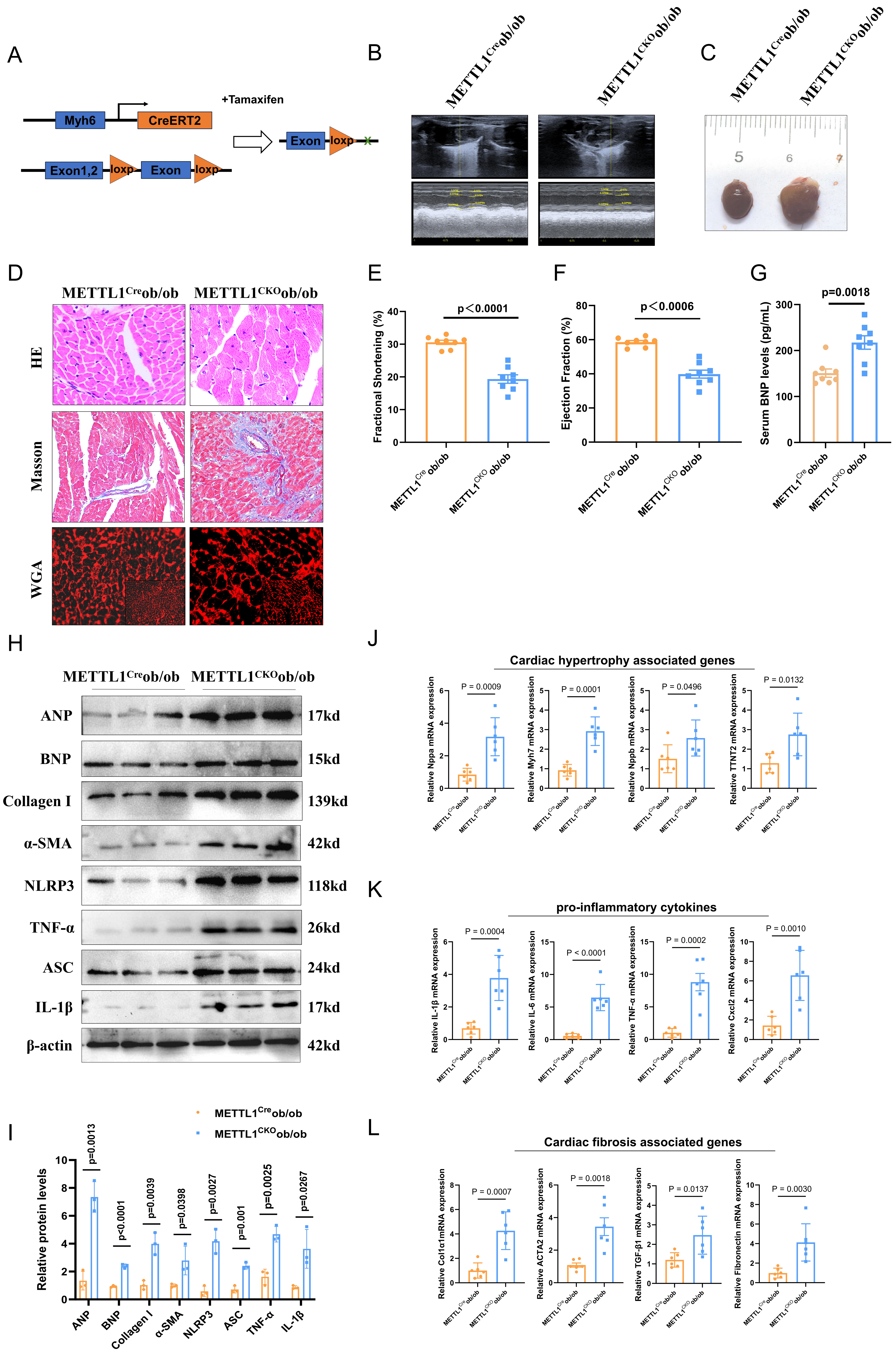
A
B
C
D
E
F
G
H
I
J
K
L
Myh6
CreERT2
Exon1,2
loxp
Exon
loxp
+Tamaxifen
Exon
loxp
METTL1Creob/ob
METTL1CKOob/ob
HE
Masson
WGA
Fractional Shortening (%)
p＜0.0001
Ejection Fraction (%)
p＜0.0006
Serum BNP levels (pg/mL)
p=0.0018
ANP 17kd
BNP 15kd
Collagen I 139kd
α-SMA 42kd
NLRP3 118kd
TNF-α 26kd
ASC 24kd
IL-1β 17kd
β-actin 42kd
Cardiac hypertrophy associated genes
Relative Nppa mRNA expression
P = 0.0009
Relative Myh7 mRNA expression
P = 0.0001
Relative Nppb mRNA expression
P = 0.0496
Relative TNNT2 mRNA expression
P = 0.0132
pro-inflammatory cytokines
Relative IL-1β mRNA expression
P = 0.0004
Relative IL-6 mRNA expression
P < 0.0001
Relative TNF-α mRNA expression
P = 0.0002
Relative Cxcl2 mRNA expression
P = 0.0010
Relative protein levels
p=0.0013
p<0.0001
p=0.0039
p=0.0398
p=0.0027
p=0.001
p=0.0025
p=0.0267
ANP
BNP
Collagen I
α-SMA
NLRP3
ASC
TNF-α
IL-1β
Cardiac fibrosis associated genes
Relative Col1α1mRNA expression
P = 0.0007
Relative ACTA2 mRNA expression
P = 0.0018
Relative TGF-β1 mRNA expression
P = 0.0137
Relative Fibronectin mRNA expression
P = 0.0030

### Figure S6

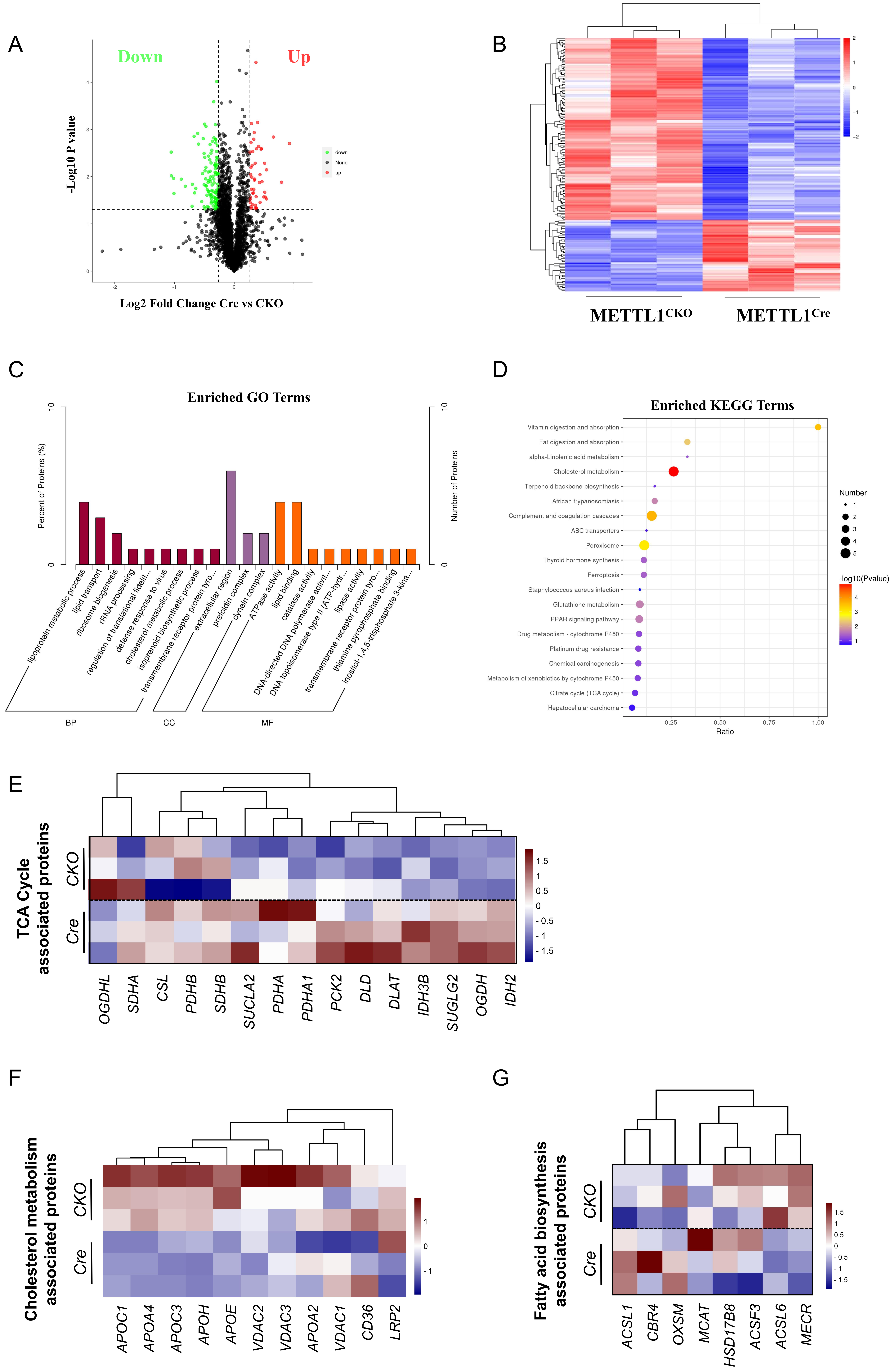

A
Down
Up
-Log10 P value
Log2 Fold Change Cre vs CKO
down
None
up
B
METTL1CKO
METTL1Cre
C
Enriched GO Terms
Percent of Proteins (%)
Number of Proteins
lipoprotein metabolic process
lipid transport
ribosome biogenesis
rRNA processing
regulation of translational fidelit...
defense response to virus
cholesterol metabolic process
isoprenoid biosynthetic process
transmembrane receptor protein tyro...
extracellular region
prefoldin complex
dynein complex
ATPase activity
lipid binding
catalase activity
DNA-directed DNA polymerase activit...
DNA topoisomerase type II (ATP-hydr...
lipase activity
transmembrane receptor protein tyro...
thiamine pyrophosphate binding
inositol-1,4,5-trisphosphate 3-kina...
BP
CC
MF
D
Enriched KEGG Terms
Vitamin digestion and absorption
Fat digestion and absorption
alpha-Linolenic acid metabolism
Cholesterol metabolism
Terpenoid backbone biosynthesis
African trypanosomiasis
Complement and coagulation cascades
ABC transporters
Peroxisome
Thyroid hormone synthesis
Ferroptosis
Staphylococcus aureus infection
Glutathione metabolism
PPAR signaling pathway
Drug metabolism - cytochrome P450
Platinum drug resistance
Chemical carcinogenesis
Metabolism of xenobiotics by cytochrome P450
Citrate cycle (TCA cycle)
Hepatocellular carcinoma
Ratio
Number
-log10(Pvalue)
E
TCA Cycle associated proteins
CKO
Cre
OGDHL
SDHA
CSL
PDHB
SDHB
SUCLA2
PDHA
PDHA1
PCK2
DLD
DLAT
IDH3B
SUGLG2
OGDH
IDH2
F
Cholesterol metabolism associated proteins
CKO
Cre
APOC1
APOA4
APOC3
APOH
APOE
VDAC2
VDAC3
APOA2
VDAC1
CD36
LRP2
G
Fatty acid biosynthesis associated proteins
CKO
Cre
ACSL1
CBR4
OXSM
MCAT
HSD17B8
ACSF3
ACSL6
MECR

Figure S7

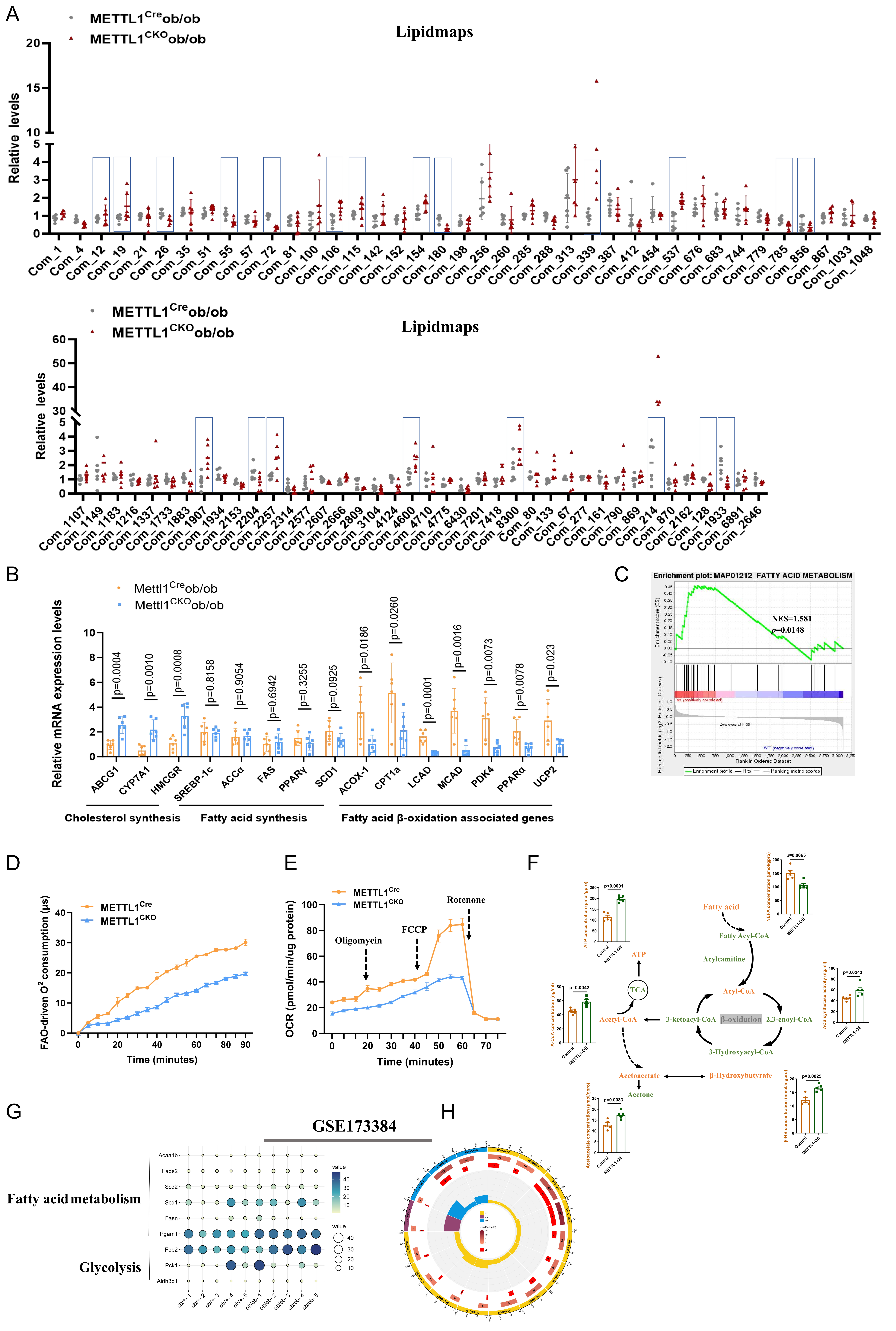
A
METTL1^Cre ob/ob
METTL1^CKO ob/ob
Lipidmaps
Relative levels
Com_1 Com_4 Com_12 Com_19 Com_21 Com_26 Com_35 Com_51 Com_55 Com_57 Com_72 Com_81 Com_100 Com_106 Com_115 Com_142 Com_152 Com_154 Com_180 Com_198 Com_256 Com_260 Com_285 Com_288 Com_313 Com_339 Com_387 Com_412 Com_454 Com_537 Com_676 Com_683 Com_744 Com_779 Com_785 Com_856 Com_867 Com_1033 Com_1048
Com_1107 Com_1149 Com_1183 Com_1216 Com_1337 Com_1733 Com_1883 Com_1907 Com_1934 Com_2153 Com_2204 Com_2257 Com_2314 Com_2577 Com_2607 Com_2666 Com_2809 Com_3104 Com_4124 Com_4600 Com_4710 Com_4775 Com_6430 Com_7201 Com_7418 Com_8300 Com_80 Com_133 Com_67 Com_277 Com_161 Com_790 Com_869 Com_214 Com_870 Com_2162 Com_128 Com_1933 Com_6891 Com_2646
B
Mettl1^Cre ob/ob
Mettl1^CKO ob/ob
Relative mRNA expression levels
p=0.0004 p=0.0010 p=0.0008 p=0.8158 p=0.9054 p=0.6942 p=0.3255 p=0.0925 p=0.0186 p=0.0260 p=0.0001 p=0.0016 p=0.0073 p=0.0078 p=0.023
ABCG1 CYP7A1 HMCGR SREBP-1c ACCα FAS PPARγ SCD1 ACOX-1 CPT1a LCAD MCAD PDK4 PPARα UCP2
Cholesterol synthesis
Fatty acid synthesis
Fatty acid β-oxidation associated genes
C
Enrichment plot: MAP01212_FATTY ACID METABOLISM
NES=1.581
p=0.0148
D
METTL1^Cre
METTL1^CKO
FAO-driven O^2 consumption (μs)
Time (minutes)
E
METTL1^Cre
METTL1^CKO
OCR (pmol/min/ug protein)
Oligomycin
FCCP
Rotenone
Time (minutes)
F
Fatty acid
Fatty Acyl-CoA
Acylcamitine
Acyl-CoA
β-oxidation
2,3-enoyl-CoA
3-Hydroxyacyl-CoA
3-ketoacyl-CoA
Acetyl-CoA
TCA
ATP
Acetoacetate
β-Hydroxybutyrate
Acetone
p<0.0001 p=0.0065 p=0.0042 p=0.0243 p=0.0083 p=0.0025
Control METTL1-OE
G
GSE173384
Fatty acid metabolism
Glycolysis
Acaa1b Fads2 Scd2 Scd1 Fasn Pgam1 Fbp2 Pck1 Aldh3b1
ob/+-1 ob/+-2 ob/+-3 ob/+-4 ob/+-5 ob/ob-1 ob/ob-2 ob/ob-3 ob/ob-4 ob/ob-5
value
H

### Figure S8

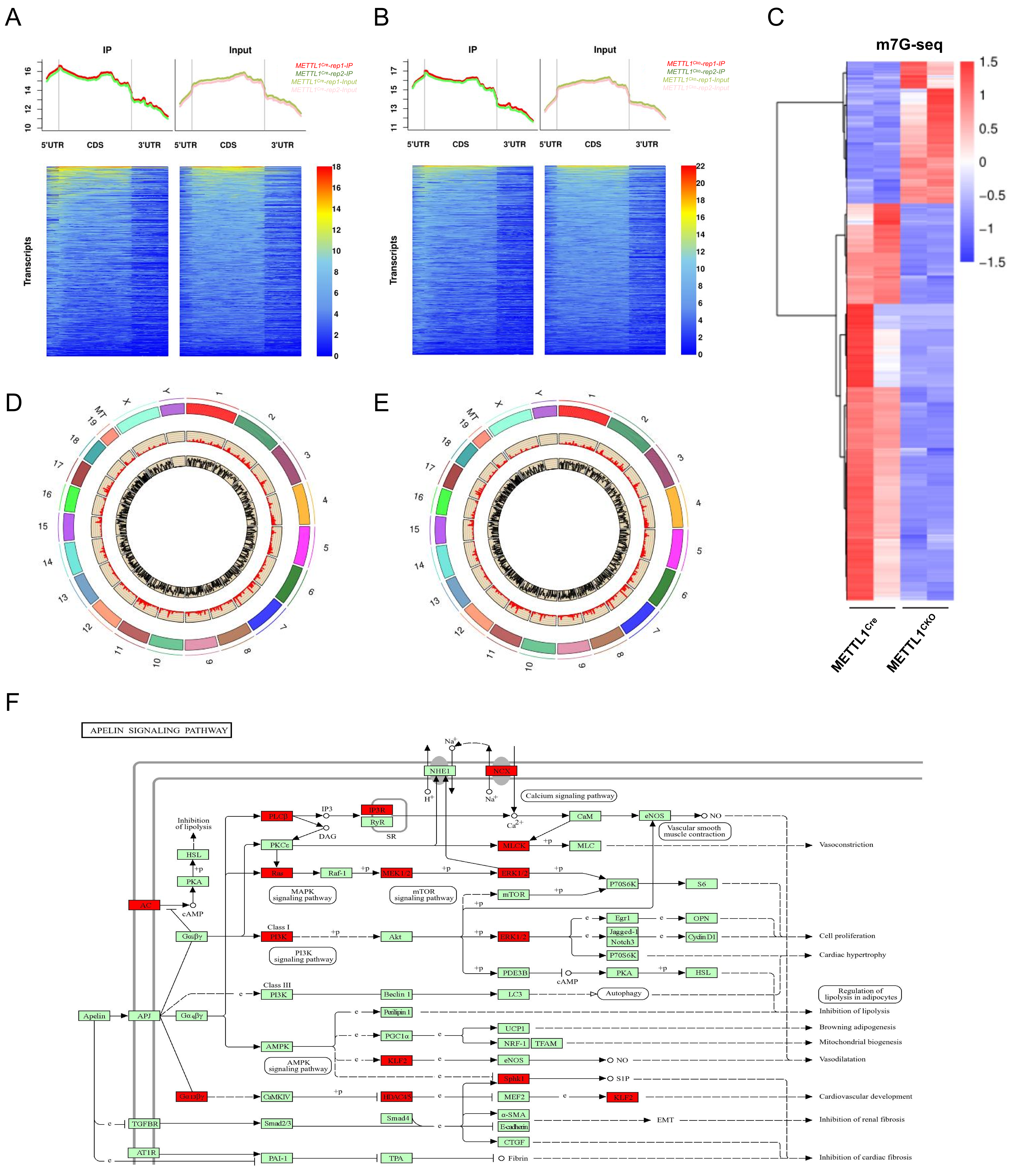

A
B
C
IP
Input
5'UTR
CDS
3'UTR
Transcripts
m7G-seq
METTL1Cre
METTL1CKO
D
E
F
APELIN SIGNALING PATHWAY
Apelin
APJ
Calcium signaling pathway
MAPK signaling pathway
mTOR signaling pathway
PI3K signaling pathway
AMPK signaling pathway
Inhibition of lipolysis
Vascular smooth muscle contraction
Autophagy
Vasoconstriction
Cell proliferation
Cardiac hypertrophy
Regulation of lipolysis in adipocytes
Inhibition of lipolysis
Browning adipogenesis
Mitochondrial biogenesis
Vasodilatation
Cardiovascular development
Inhibition of renal fibrosis
Inhibition of cardiac fibrosis

### Figure S9

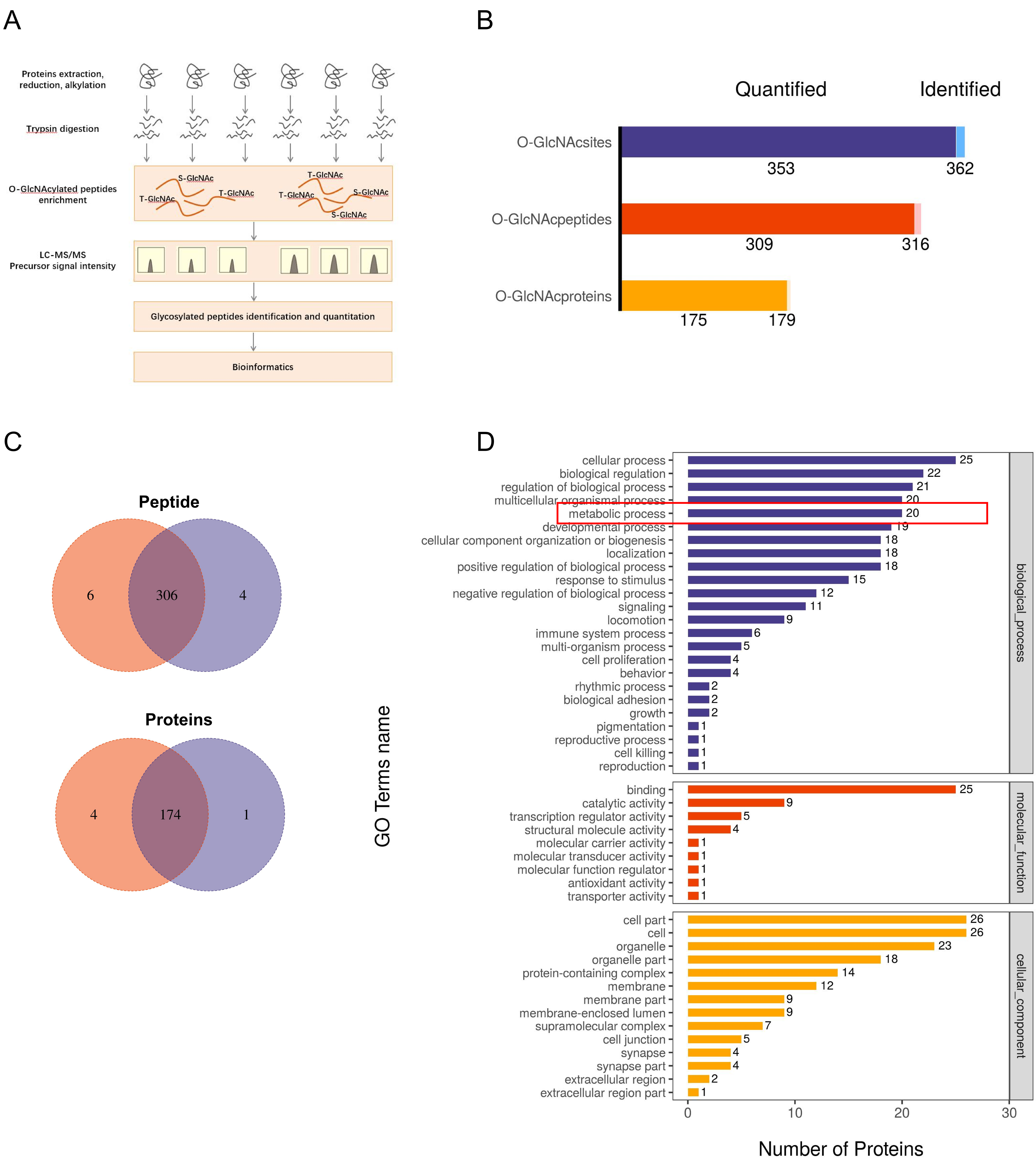
A
Proteins extraction, reduction, alkylation
Trypsin digestion
O-GlcNAcylated peptides enrichment
S-GlcNAc
T-GlcNAc
LC-MS/MS
Precursor signal intensity
Glycosylated peptides identification and quantitation
Bioinformatics
B
Quantified
Identified
O-GlcNAcsites
O-GlcNAcpeptides
O-GlcNAcproteins
353
362
309
316
175
179
C
Peptide
6
306
4
Proteins
4
174
1
D
GO Terms name
cellular process
biological regulation
regulation of biological process
multicellular organismal process
metabolic process
developmental process
cellular component organization or biogenesis
localization
positive regulation of biological process
response to stimulus
negative regulation of biological process
signaling
locomotion
immune system process
multi-organism process
cell proliferation
behavior
rhythmic process
biological adhesion
growth
pigmentation
reproductive process
cell killing
reproduction
binding
catalytic activity
transcription regulator activity
structural molecule activity
molecular carrier activity
molecular transducer activity
molecular function regulator
antioxidant activity
transporter activity
cell part
cell
organelle
organelle part
protein-containing complex
membrane
membrane part
membrane-enclosed lumen
supramolecular complex
cell junction
synapse
synapse part
extracellular region
extracellular region part
biological_process
molecular_function
cellular_component
Number of Proteins
0
10
20
30

Figure S10

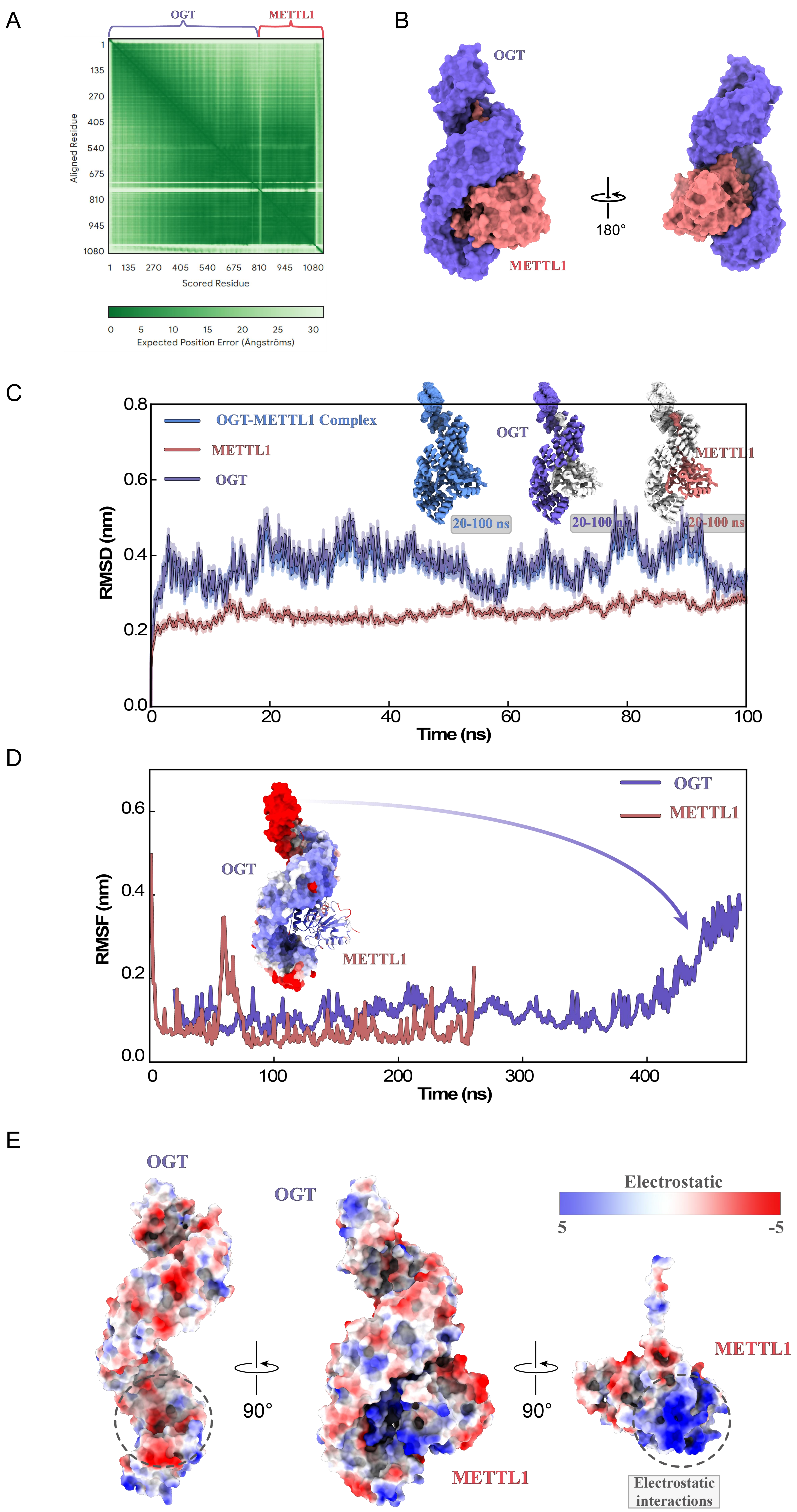
A
OGT
METTL1
Aligned Residue
Scored Residue
Expected Position Error (Ångströms)
B
OGT
METTL1
180°
C
OGT-METTL1 Complex
METTL1
OGT
20-100 ns
RMSD (nm)
Time (ns)
D
OGT
METTL1
RMSF (nm)
Time (ns)
E
OGT
METTL1
Electrostatic
5
-5
90°
Electrostatic interactions

Figure S11

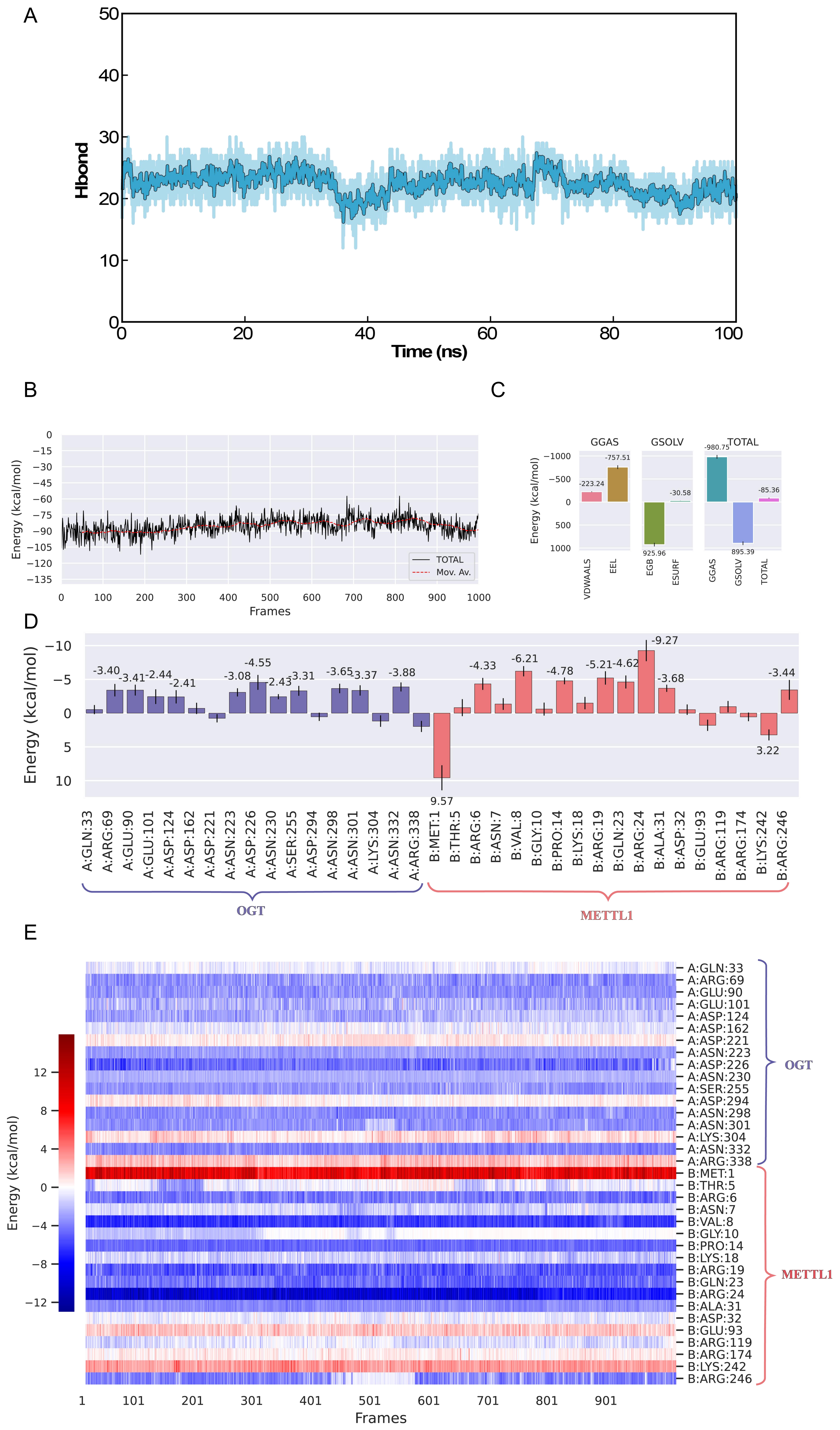

A
Hbond
Time (ns)
B
Energy (kcal/mol)
Frames
TOTAL
Mov. Av.
C
GGAS
GSOLV
TOTAL
VDWAALS
EEL
EGB
ESURF
GGAS
GSOLV
TOTAL
-223.24
-757.51
925.96
-30.58
-980.75
895.39
-85.36
D
Energy (kcal/mol)
A:GLN:33
A:ARG:69
A:GLU:90
A:GLU:101
A:ASP:124
A:ASP:162
A:ASP:221
A:ASN:223
A:ASP:226
A:ASN:230
A:SER:255
A:ASP:294
A:ASN:298
A:ASN:301
A:LYS:304
A:ASN:332
A:ARG:338
B:MET:1
B:THR:5
B:ARG:6
B:ASN:7
B:VAL:8
B:GLY:10
B:PRO:14
B:LYS:18
B:ARG:19
B:GLN:23
B:ARG:24
B:ALA:31
B:ASP:32
B:GLU:93
B:ARG:119
B:ARG:174
B:LYS:242
B:ARG:246
OGT
METTL1
E
Energy (kcal/mol)
Frames
OGT
METTL1

### Figure S12

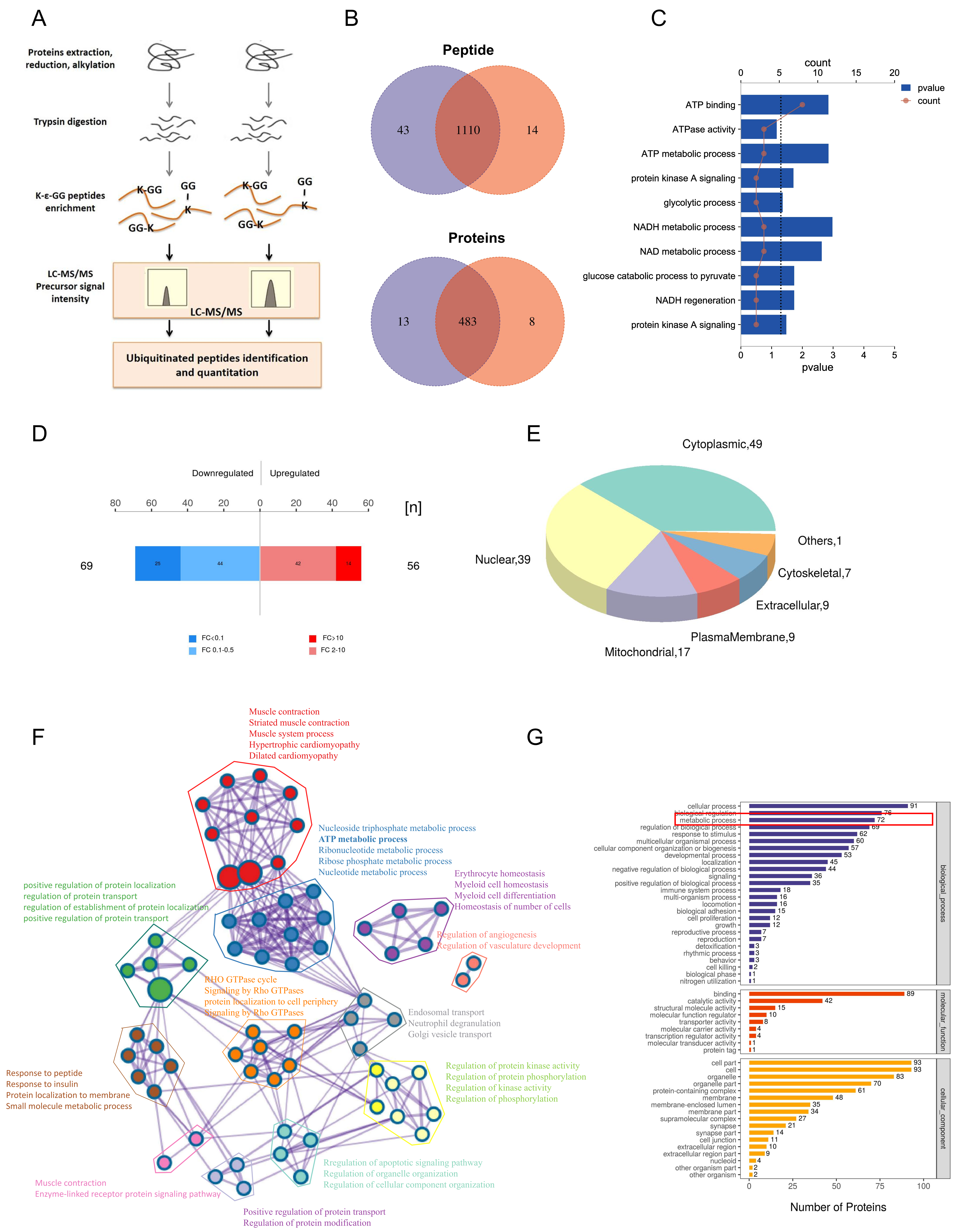

A
Proteins extraction, reduction, alkylation
Trypsin digestion
K-ε-GG peptides enrichment
K-GG
GG
K
GG-K
LC-MS/MS Precursor signal intensity
LC-MS/MS
Ubiquitinated peptides identification and quantitation
B
Peptide
43
1110
14
Proteins
13
483
8
C
count
0
5
10
15
20
pvalue
count
ATP binding
ATPase activity
ATP metabolic process
protein kinase A signaling
glycolytic process
NADH metabolic process
NAD metabolic process
glucose catabolic process to pyruvate
NADH regeneration
protein kinase A signaling
0
1
2
3
4
5
pvalue
D
Downregulated
Upregulated
80
60
40
20
0
20
40
60
[n]
69
25
44
42
14
56
FC<0.1
FC 0.1-0.5
FC>10
FC 2-10
E
Cytoplasmic,49
Others,1
Cytoskeletal,7
Extracellular,9
PlasmaMembrane,9
Mitochondrial,17
Nuclear,39
F
Muscle contraction
Striated muscle contraction
Muscle system process
Hypertrophic cardiomyopathy
Dilated cardiomyopathy
Nucleoside triphosphate metabolic process
ATP metabolic process
Ribonucleotide metabolic process
Ribose phosphate metabolic process
Nucleotide metabolic process
positive regulation of protein localization
regulation of protein transport
regulation of establishment of protein localization
positive regulation of protein transport
Erythrocyte homeostasis
Myeloid cell homeostasis
Myeloid cell differentiation
Homeostasis of number of cells
Regulation of angiogenesis
Regulation of vasculature development
RHO GTPase cycle
Signaling by Rho GTPases
protein localization to cell periphery
Signaling by Rho GTPases
Endosomal transport
Neutrophil degranulation
Golgi vesicle transport
Response to peptide
Response to insulin
Protein localization to membrane
Small molecule metabolic process
Regulation of protein kinase activity
Regulation of protein phosphorylation
Regulation of kinase activity
Regulation of phosphorylation
Regulation of apoptotic signaling pathway
Regulation of organelle organization
Regulation of cellular component organization
Muscle contraction
Enzyme-linked receptor protein signaling pathway
Positive regulation of protein transport
Regulation of protein modification
G
cellular process 91
biological regulation 76
metabolic process 72
regulation of biological process 69
response to stimulus 62
multicellular organismal process 60
cellular component organization or biogenesis 57
developmental process 53
localization 45
negative regulation of biological process 44
signaling 36
positive regulation of biological process 35
immune system process 18
multi-organism process 16
locomotion 16
biological adhesion 15
cell proliferation 12
growth 12
reproductive process 7
reproduction 7
detoxification 3
rhythmic process 3
behavior 3
cell killing 2
biological phase 1
nitrogen utilization 1
biological_process
binding 89
catalytic activity 42
structural molecule activity 15
molecular function regulator 10
transporter activity 8
molecular carrier activity 4
transcription regulator activity 4
molecular transducer activity 1
protein tag 1
molecular_function
cell part 93
cell 93
organelle 83
organelle part 70
protein-containing complex 61
membrane 48
membrane-enclosed lumen 35
membrane part 34
supramolecular complex 27
synapse 21
synapse part 14
cell junction 11
extracellular region 10
extracellular region part 9
nucleoid 4
other organism part 2
other organism 2
cellular_component
0
25
50
75
100
Number of Proteins

Figure S13

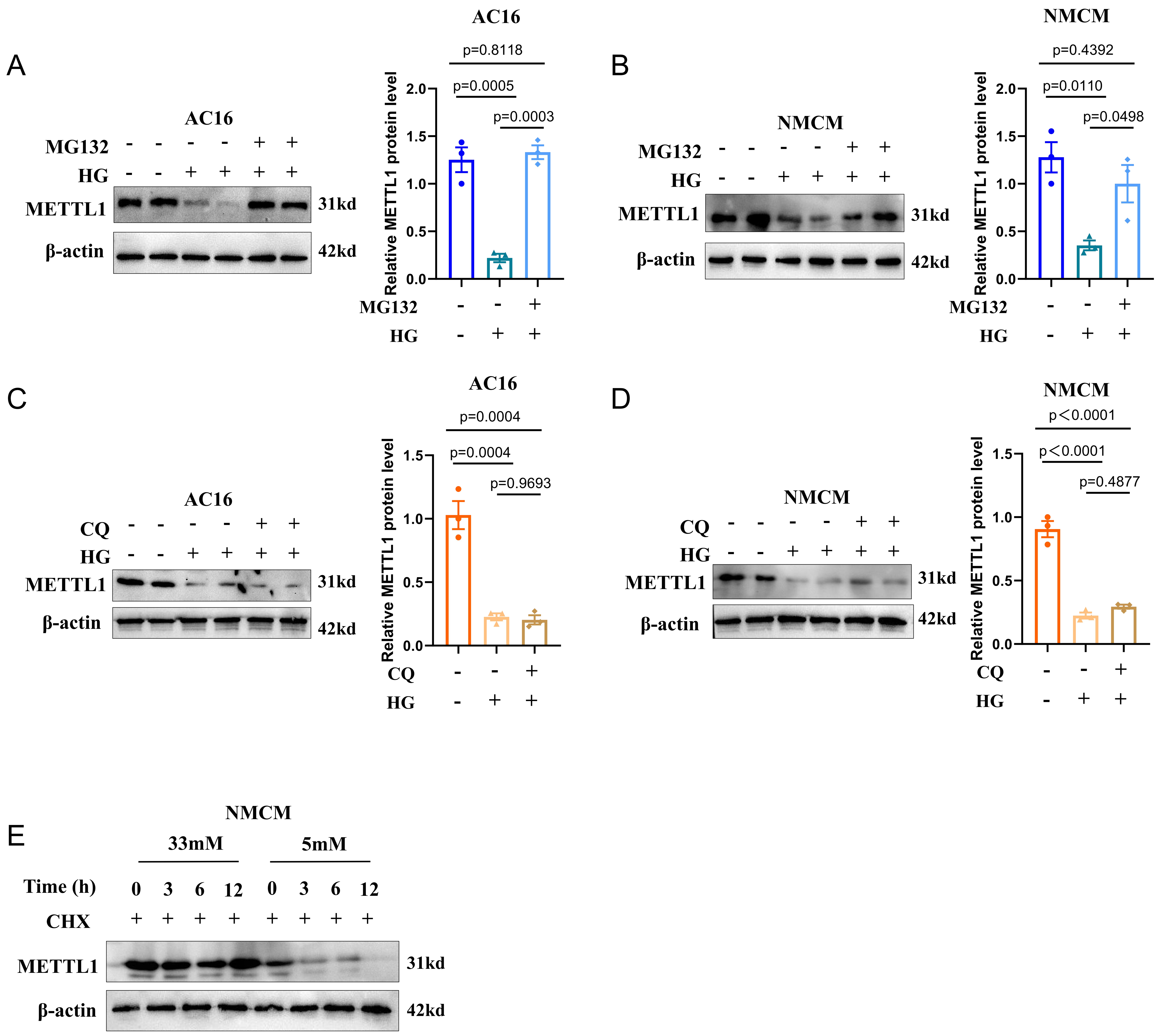
A
AC16
MG132
HG
METTL1
31kd
β-actin
42kd
AC16
p=0.8118
p=0.0005
p=0.0003
Relative METTL1 protein level
B
NMCM
p=0.4392
p=0.0110
p=0.0498
C
CQ
p=0.0004
p=0.0004
p=0.9693
D
p<0.0001
p<0.0001
p=0.4877
E
NMCM
33mM
5mM
Time (h)
0 3 6 12 0 3 6 12
CHX

Figure S14

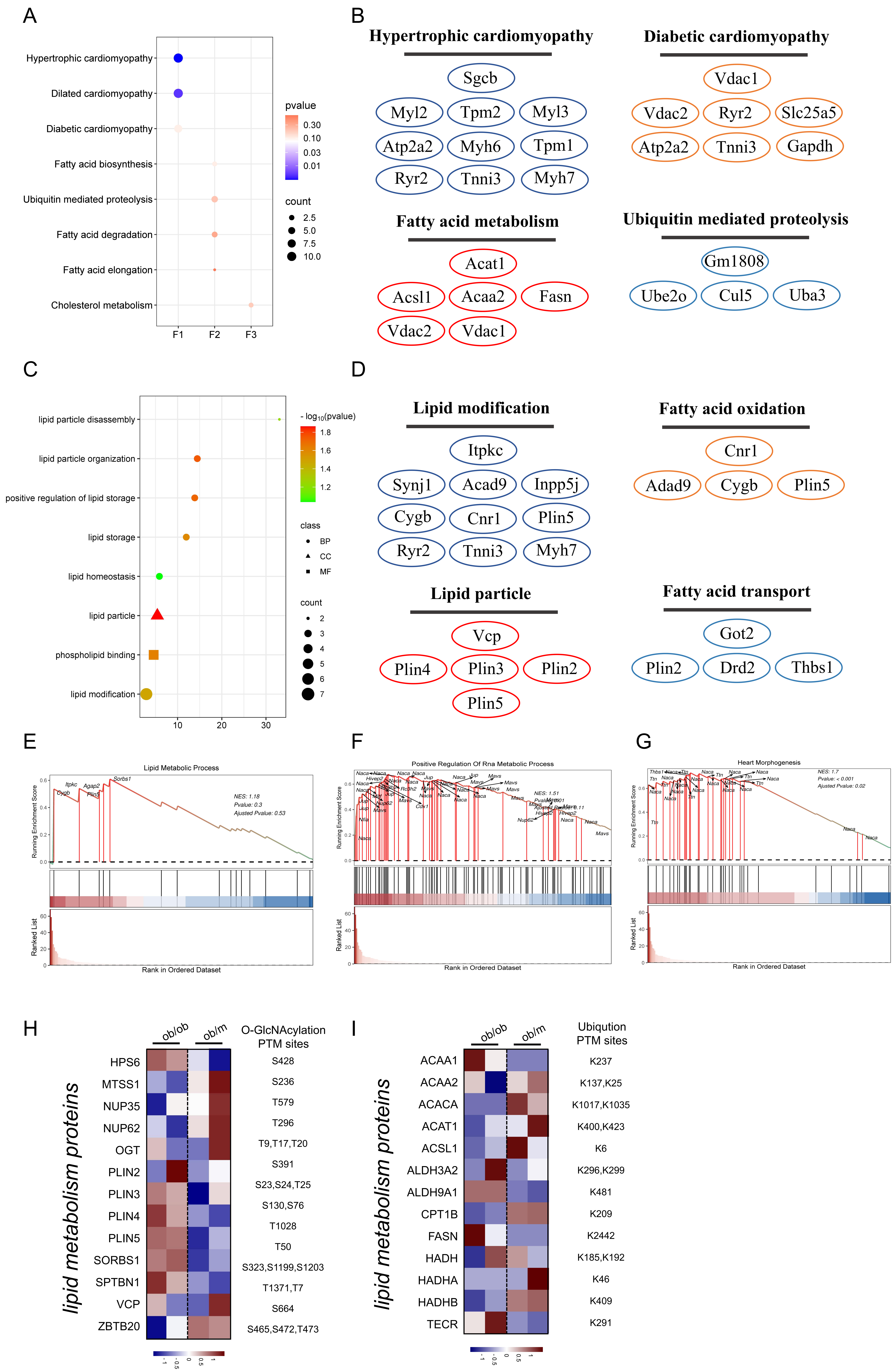
A
Hypertrophic cardiomyopathy
Dilated cardiomyopathy
Diabetic cardiomyopathy
Fatty acid biosynthesis
Ubiquitin mediated proteolysis
Fatty acid degradation
Fatty acid elongation
Cholesterol metabolism
F1
F2
F3
pvalue
0.30
0.10
0.03
0.01
count
2.5
5.0
7.5
10.0
B
Hypertrophic cardiomyopathy
Sgcb
Myl2
Tpm2
Myl3
Atp2a2
Myh6
Tpm1
Ryr2
Tnni3
Myh7
Diabetic cardiomyopathy
Vdac1
Vdac2
Ryr2
Slc25a5
Atp2a2
Tnni3
Gapdh
Fatty acid metabolism
Acat1
Acsl1
Acaa2
Fasn
Vdac2
Vdac1
Ubiquitin mediated proteolysis
Gm1808
Ube2o
Cul5
Uba3
C
lipid particle disassembly
lipid particle organization
positive regulation of lipid storage
lipid storage
lipid homeostasis
lipid particle
phospholipid binding
lipid modification
10
20
30
- log10(pvalue)
1.8
1.6
1.4
1.2
class
BP
CC
MF
count
2
3
4
5
6
7
D
Lipid modification
Itpkc
Synj1
Acad9
Inpp5j
Cygb
Cnr1
Plin5
Ryr2
Tnni3
Myh7
Fatty acid oxidation
Cnr1
Adad9
Cygb
Plin5
Lipid particle
Vcp
Plin4
Plin3
Plin2
Plin5
Fatty acid transport
Got2
Plin2
Drd2
Thbs1
E
Lipid Metabolic Process
NES: 1.18
Pvalue: 0.3
Ajusted Pvalue: 0.53
Running Enrichment Score
Ranked List
Rank in Ordered Dataset
F
Positive Regulation Of Rna Metabolic Process
NES: 1.51
Running Enrichment Score
Ranked List
Rank in Ordered Dataset
G
Heart Morphogenesis
NES: 1.7
Pvalue: < 0.001
Ajusted Pvalue: 0.02
Running Enrichment Score
Ranked List
Rank in Ordered Dataset
H
ob/ob
ob/m
O-GlcNAcylation PTM sites
lipid metabolism proteins
HPS6 S428
MTSS1 S236
NUP35 T579
NUP62 T296
OGT T9,T17,T20
PLIN2 S391
PLIN3 S23,S24,T25
PLIN4 S130,S76
PLIN5 T1028
SORBS1 T50
SPTBN1 S323,S1199,S1203
VCP T1371,T7
ZBTB20 S664
S465,S472,T473
-1
-0.5
0
0.5
1
I
ob/ob
ob/m
Ubiqution PTM sites
lipid metabolism proteins
ACAA1 K237
ACAA2 K137,K25
ACACA K1017,K1035
ACAT1 K400,K423
ACSL1 K6
ALDH3A2 K296,K299
ALDH9A1 K481
CPT1B K209
FASN K2442
HADH K185,K192
HADHA K46
HADHB K409
TECR K291
-1
-0.5
0
0.5
1

Figure S15

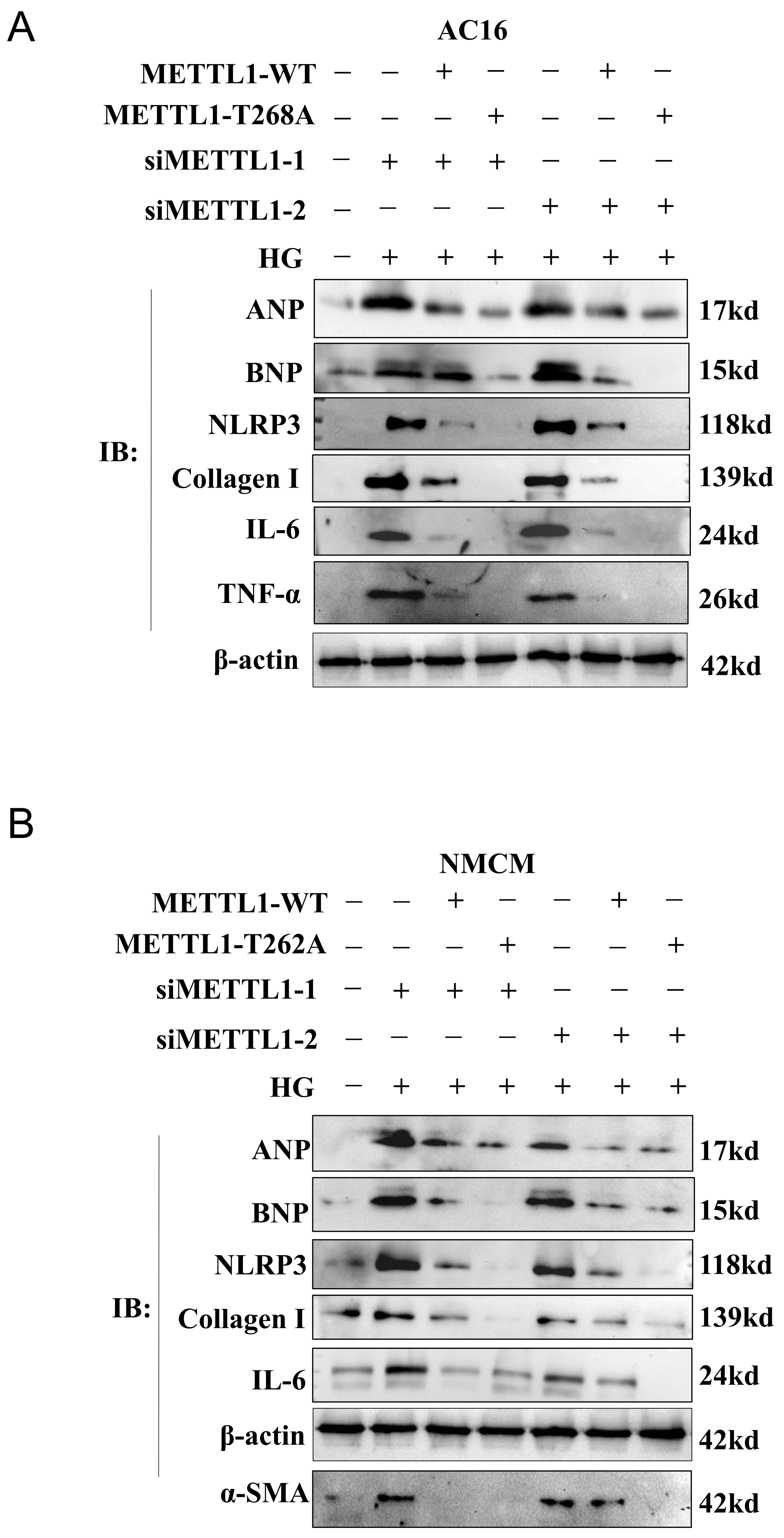
A
AC16
METTL1-WT
METTL1-T268A
siMETTL1-1
siMETTL1-2
HG
IB:
ANP
BNP
NLRP3
Collagen I
IL-6
TNF-α
β-actin
17kd
15kd
118kd
139kd
24kd
26kd
42kd
B
NMCM
METTL1-WT
METTL1-T262A
siMETTL1-1
siMETTL1-2
HG
IB:
ANP
BNP
NLRP3
Collagen I
IL-6
β-actin
α-SMA
17kd
15kd
118kd
139kd
24kd
42kd
42kd

Figure S16

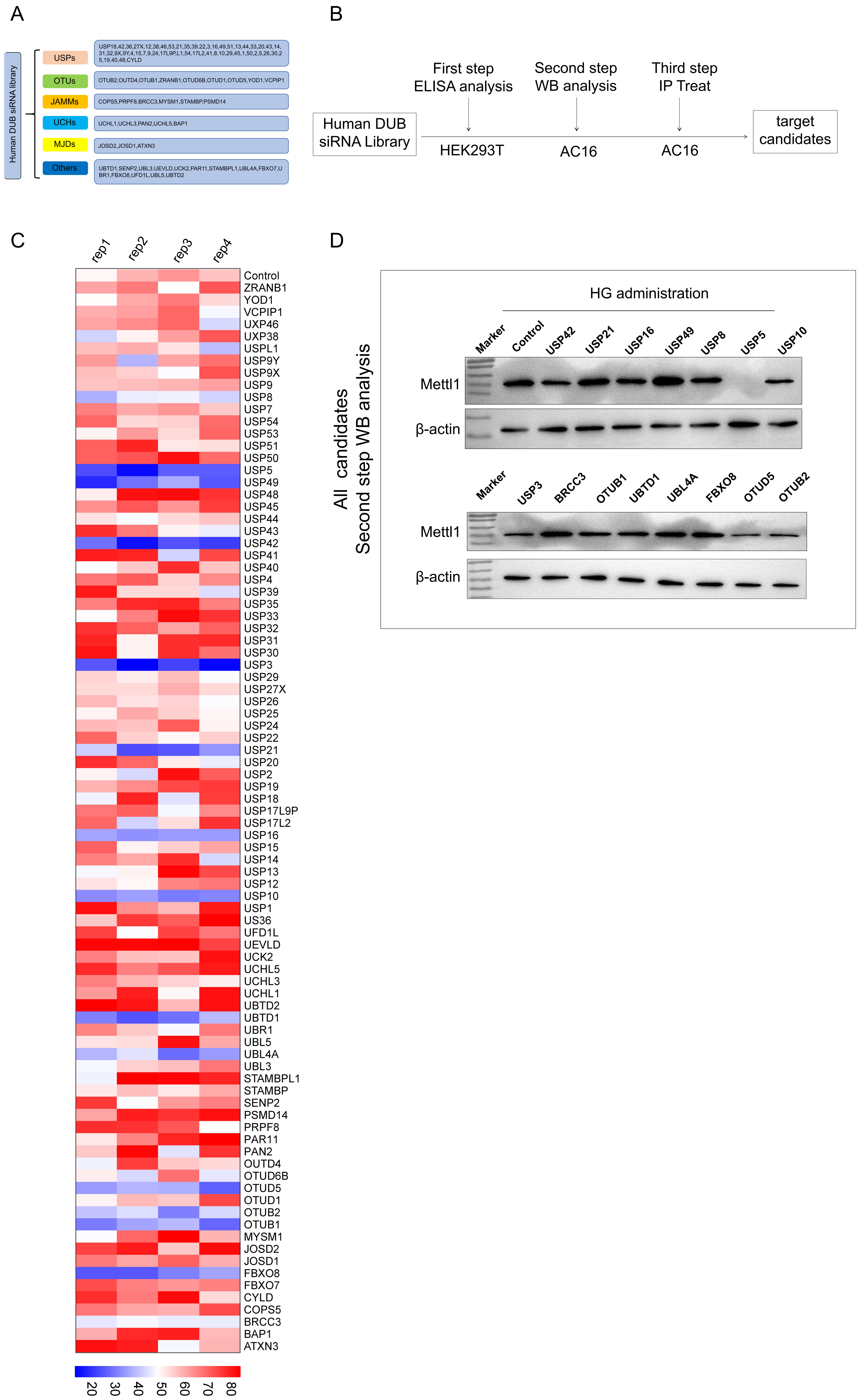
A
Human DUB siRNA library
USPs
USP18,42,36,27X,12,38,46,53,21,35,39.22,3,16,49,51,13,44,33,20,43,14,31,32,9X,9Y,4,15,7,9,24,17L9P,L1,54,17L2,41,8,10,29,45,1,50,2,5,26,30,25,19,40,48,CYLD
OTUs
OTUB2,OUTD4,OTUB1,ZRANB1,OTUD6B,OTUD1,OTUD5,YOD1,VCPIP1
JAMMs
COPS5,PRPF8,BRCC3,MYSM1,STAMBP,PSMD14
UCHs
UCHL1,UCHL3,PAN2,UCHL5,BAP1
MJDs
JOSD2,JOSD1,ATXN3
Others
UBTD1,SENP2,UBL3,UEVLD,UCK2,PAR11,STAMBPL1,UBL4A,FBXO7,UBR1,FBXO8,UFD1L,UBL5,UBTD2
B
First step
ELISA analysis
Second step
WB analysis
Third step
IP Treat
Human DUB siRNA Library
HEK293T
AC16
AC16
target candidates
C
rep1
rep2
rep3
rep4
Control
ZRANB1
YOD1
VCPIP1
UXP46
UXP38
USPL1
USP9Y
USP9X
USP9
USP8
USP7
USP54
USP53
USP51
USP50
USP5
USP49
USP48
USP45
USP44
USP43
USP42
USP41
USP40
USP4
USP39
USP35
USP33
USP32
USP31
USP30
USP3
USP29
USP27X
USP26
USP25
USP24
USP22
USP21
USP20
USP2
USP19
USP18
USP17L9P
USP17L2
USP16
USP15
USP14
USP13
USP12
USP10
USP1
US36
UFD1L
UEVLD
UCK2
UCHL5
UCHL3
UCHL1
UBTD2
UBTD1
UBR1
UBL5
UBL4A
UBL3
STAMBPL1
STAMBP
SENP2
PSMD14
PRPF8
PAR11
PAN2
OUTD4
OTUD6B
OTUD5
OTUD1
OTUB2
OTUB1
MYSM1
JOSD2
JOSD1
FBXO8
FBXO7
CYLD
COPS5
BRCC3
BAP1
ATXN3
20 30 40 50 60 70 80
D
All candidates
Second step WB analysis
HG administration
Marker
Control
USP42
USP21
USP16
USP49
USP8
USP5
USP10
Mettl1
β-actin
Marker
USP3
BRCC3
OTUB1
UBTD1
UBL4A
FBXO8
OTUD5
OTUB2
Mettl1
β-actin

Figure S17

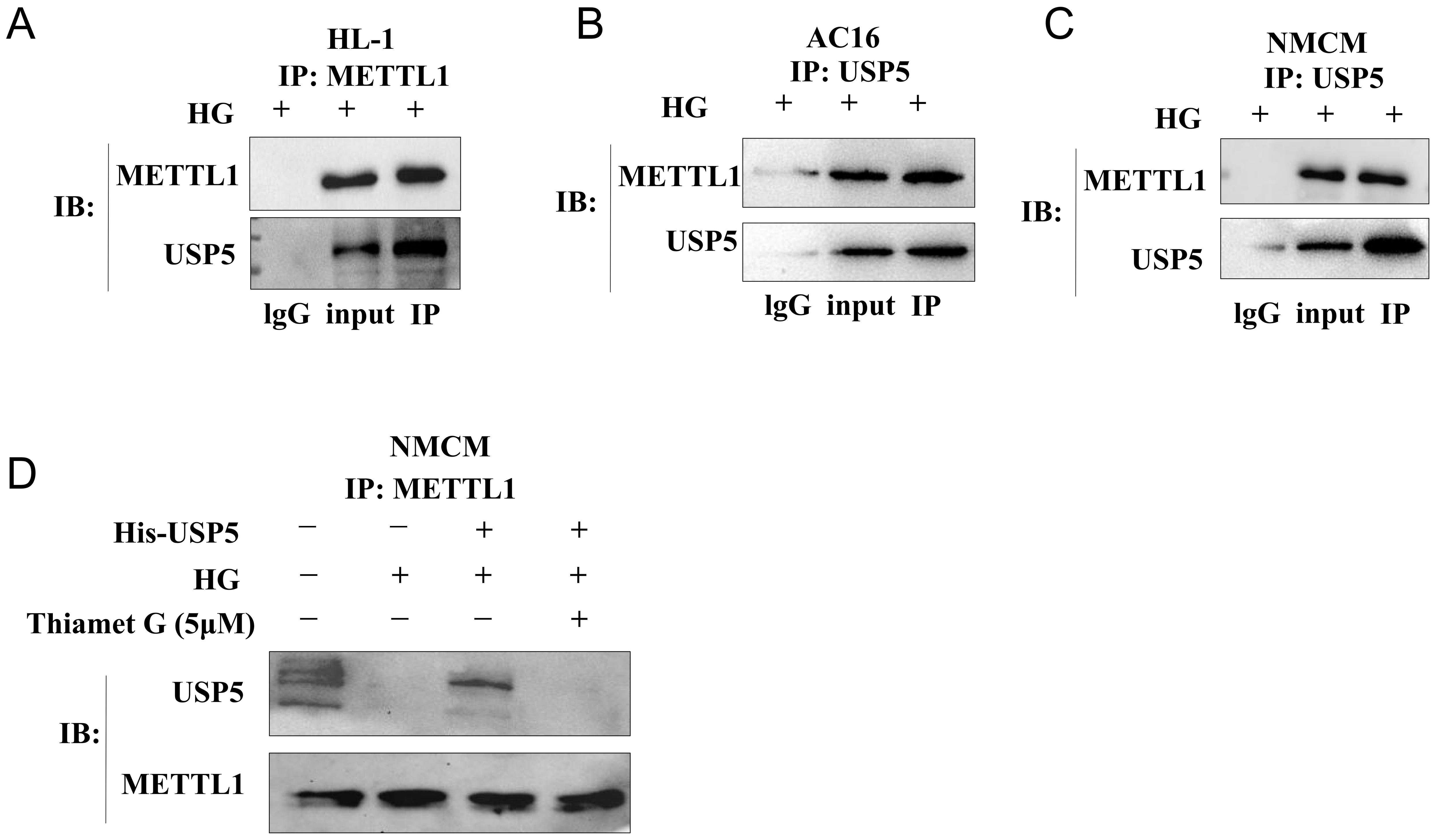
A
HL-1
IP: METTL1
HG + + +
IB:
METTL1
USP5
lgG input IP
B
AC16
IP: USP5
HG + + +
IB:
METTL1
USP5
lgG input IP
C
NMCM
IP: USP5
HG + + +
IB:
METTL1
USP5
lgG input IP
D
NMCM
IP: METTL1
His-USP5 − − + +
HG − + + +
Thiamet G (5μM) − − − +
IB:
USP5
METTL1

Figure S18

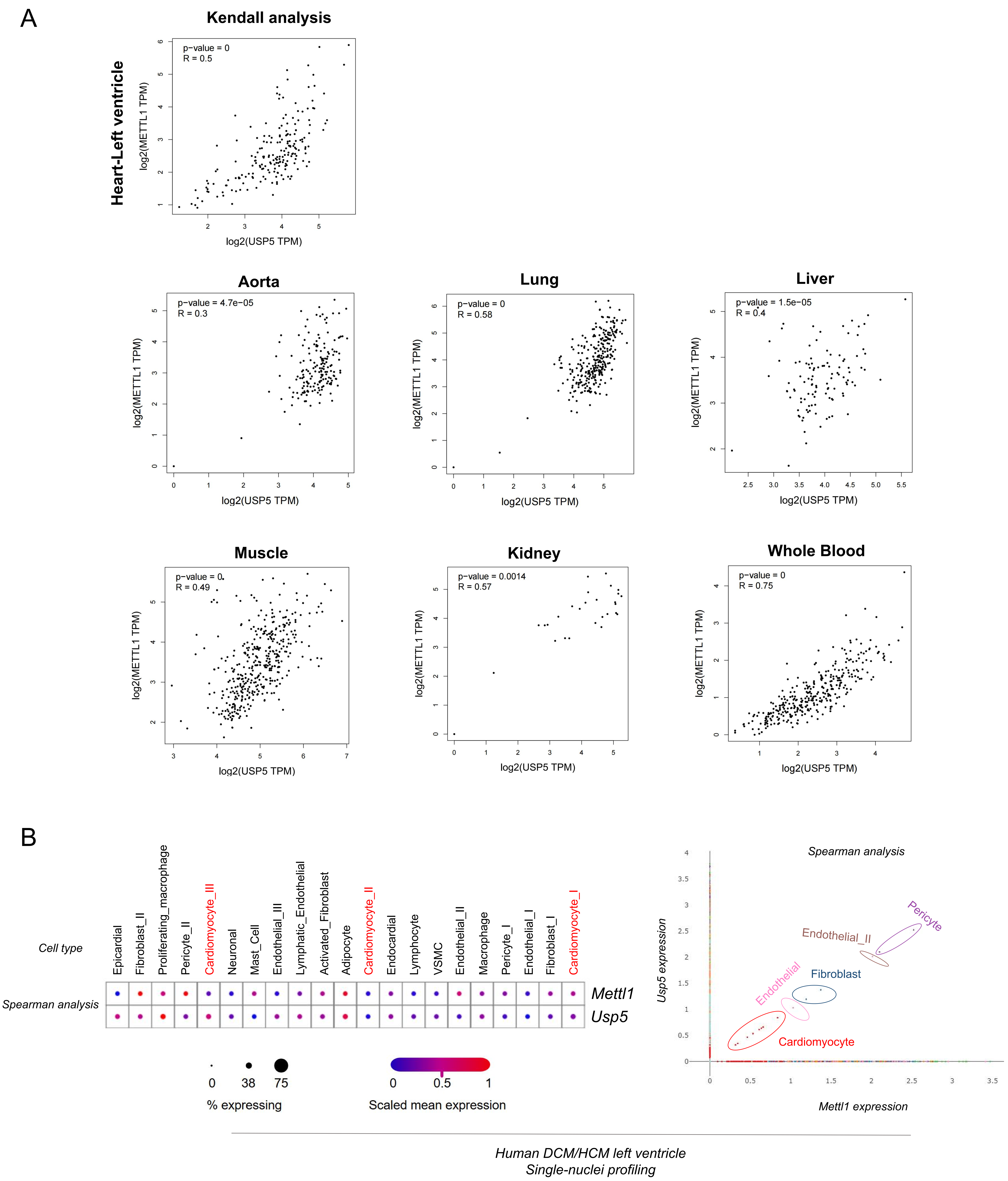
A
Kendall analysis
Heart-Left ventricle
p−value = 0
R = 0.5
log2(METTL1 TPM)
log2(USP5 TPM)
Aorta
p−value = 4.7e−05
R = 0.3
Lung
p−value = 0
R = 0.58
Liver
p−value = 1.5e−05
R = 0.4
Muscle
p−value = 0
R = 0.49
Kidney
p−value = 0.0014
R = 0.57
Whole Blood
p−value = 0
R = 0.75
B
Cell type
Spearman analysis
Epicardial
Fibroblast_II
Proliferating_macrophage
Pericyte_II
Cardiomyocyte_III
Neuronal
Mast_Cell
Endothelial_III
Lymphatic_Endothelial
Activated_Fibroblast
Adipocyte
Cardiomyocyte_II
Endocardial
Lymphocyte
VSMC
Endothelial_II
Macrophage
Pericyte_I
Endothelial_I
Fibroblast_I
Cardiomyocyte_I
Mettl1
Usp5
0
38
75
% expressing
0
0.5
1
Scaled mean expression
Spearman analysis
Usp5 expression
Mettl1 expression
Pericyte
Endothelial_II
Endothelial
Fibroblast
Cardiomyocyte
Human DCM/HCM left ventricle
Single-nuclei profiling

### Figure S19

A *HIT104371060*

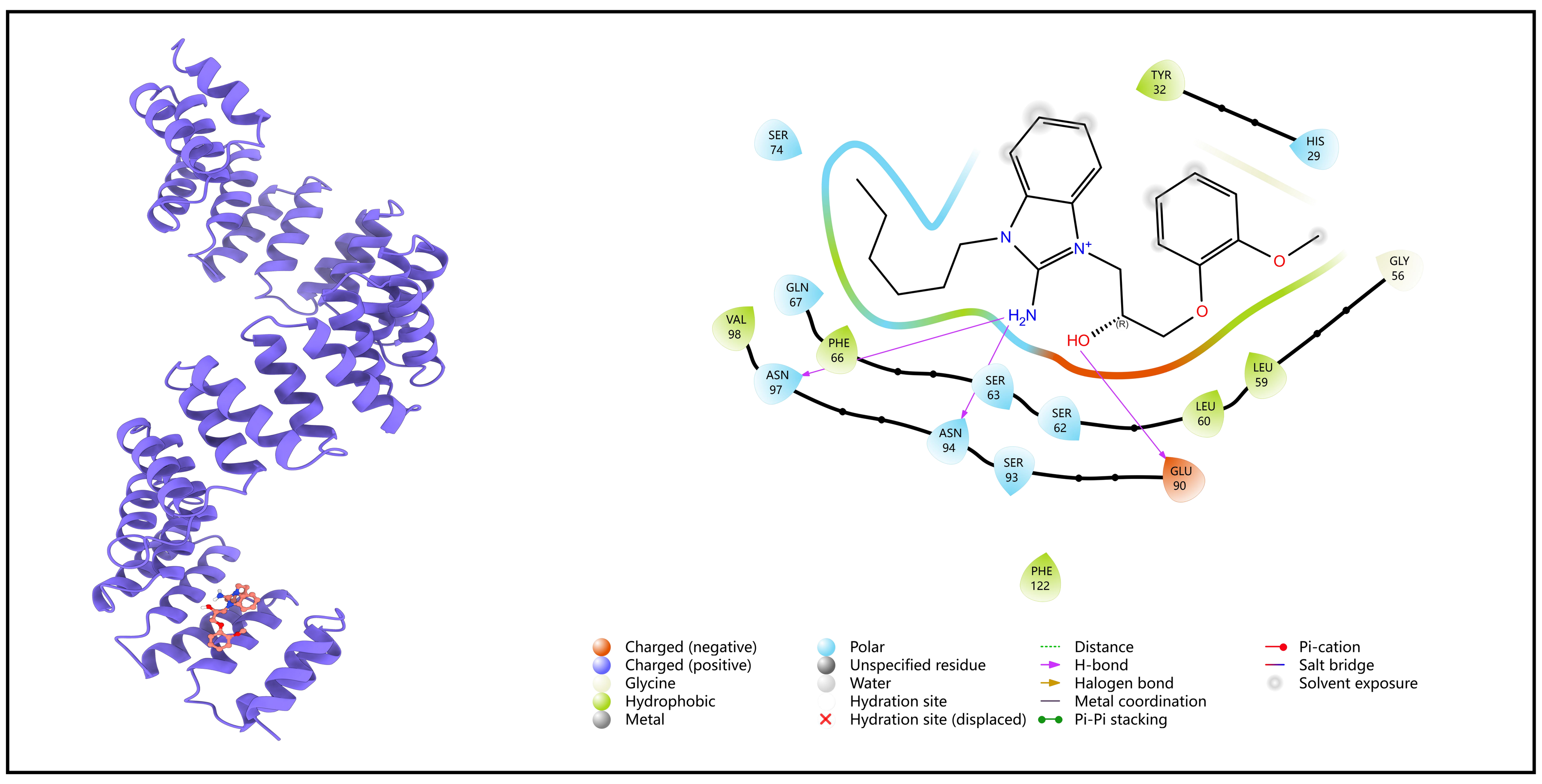

B *HIT106265621*

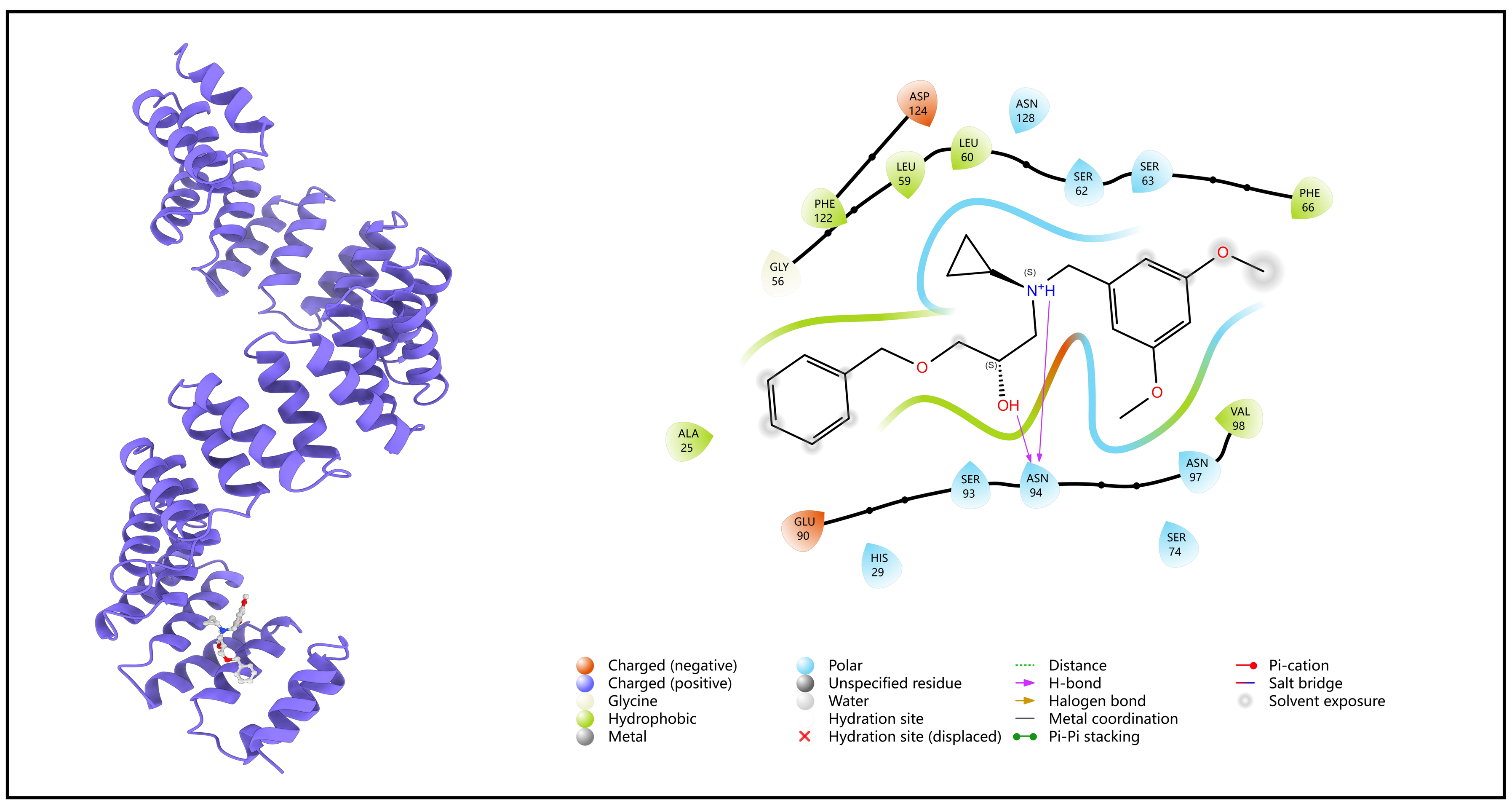

C *HIT101296070*

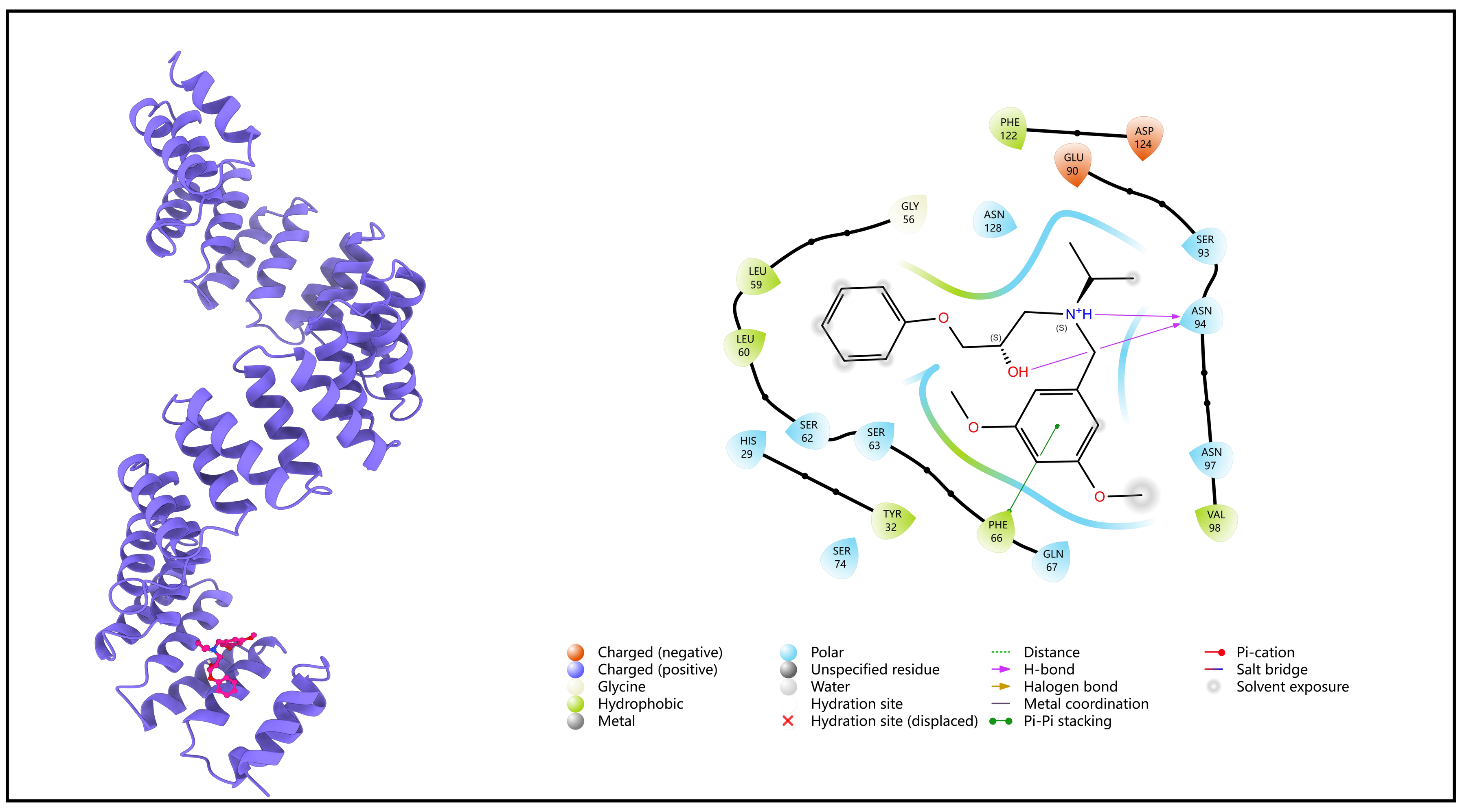

D *HIT103099087*

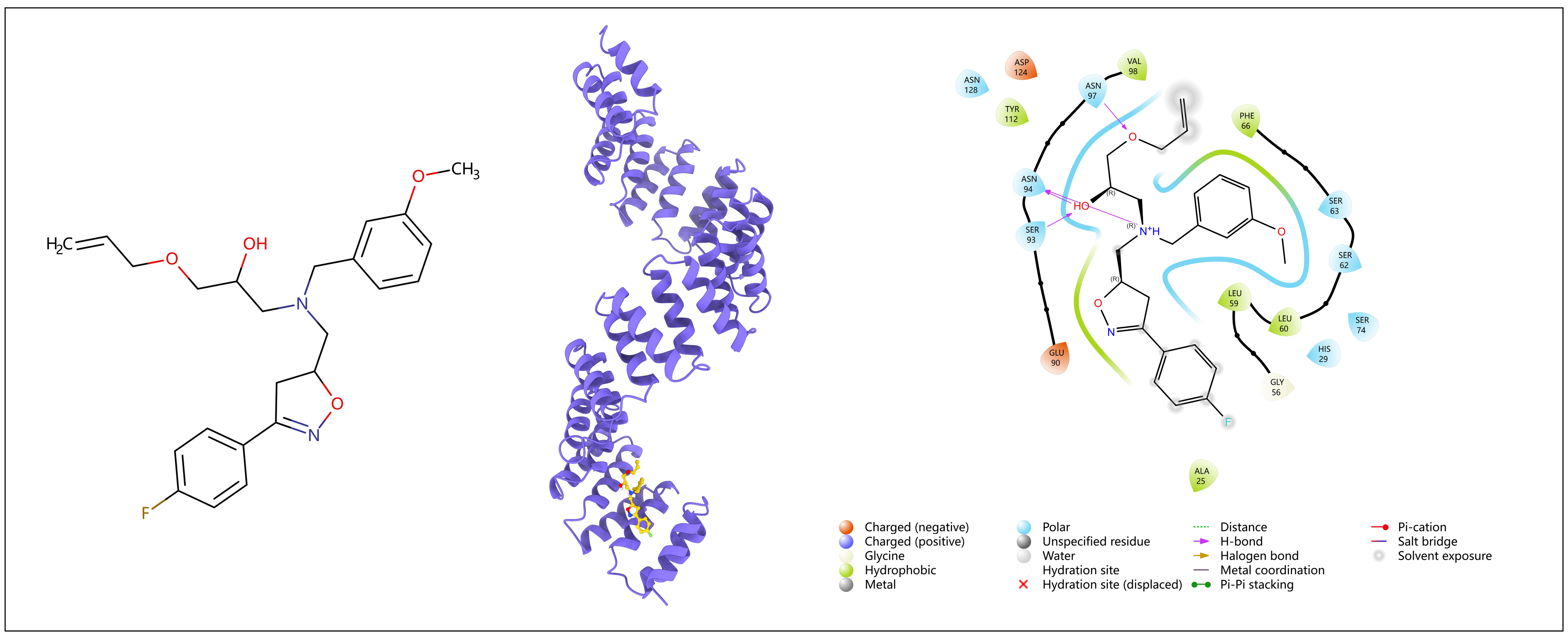

E *HIT104933928*

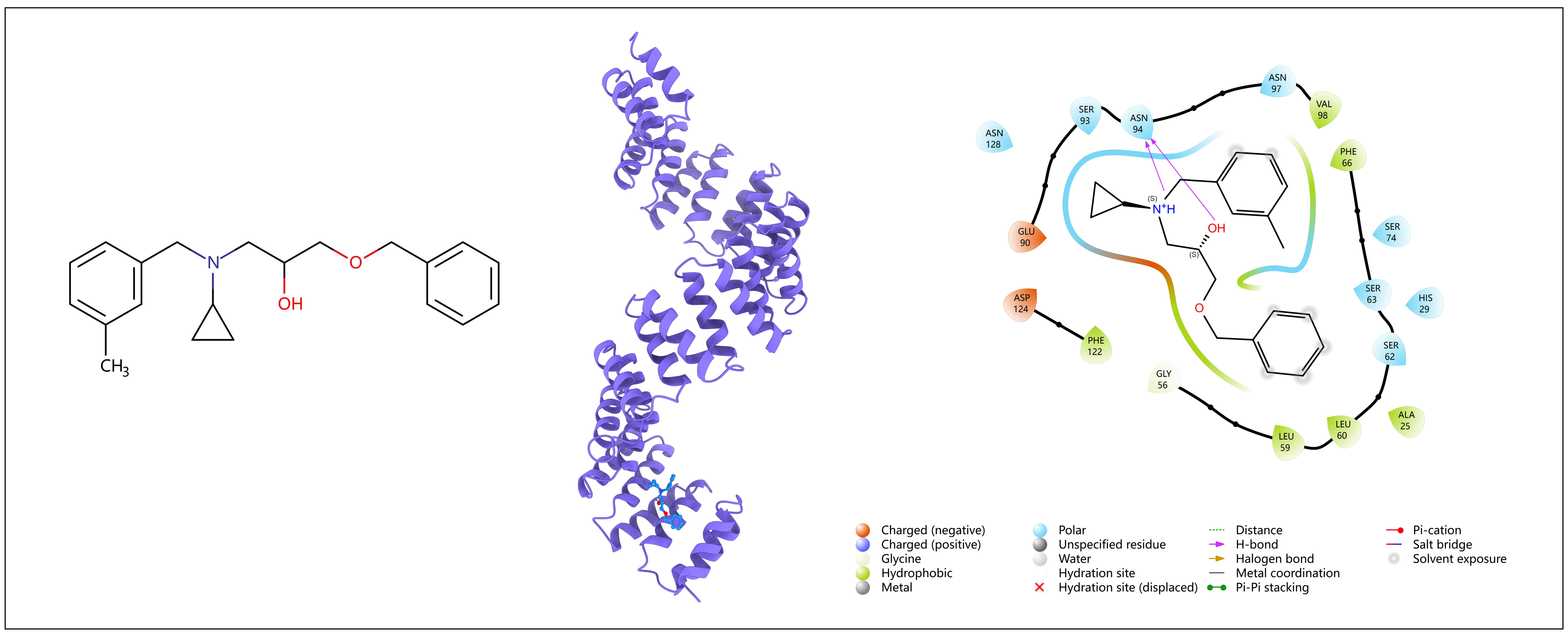

### Figure S20

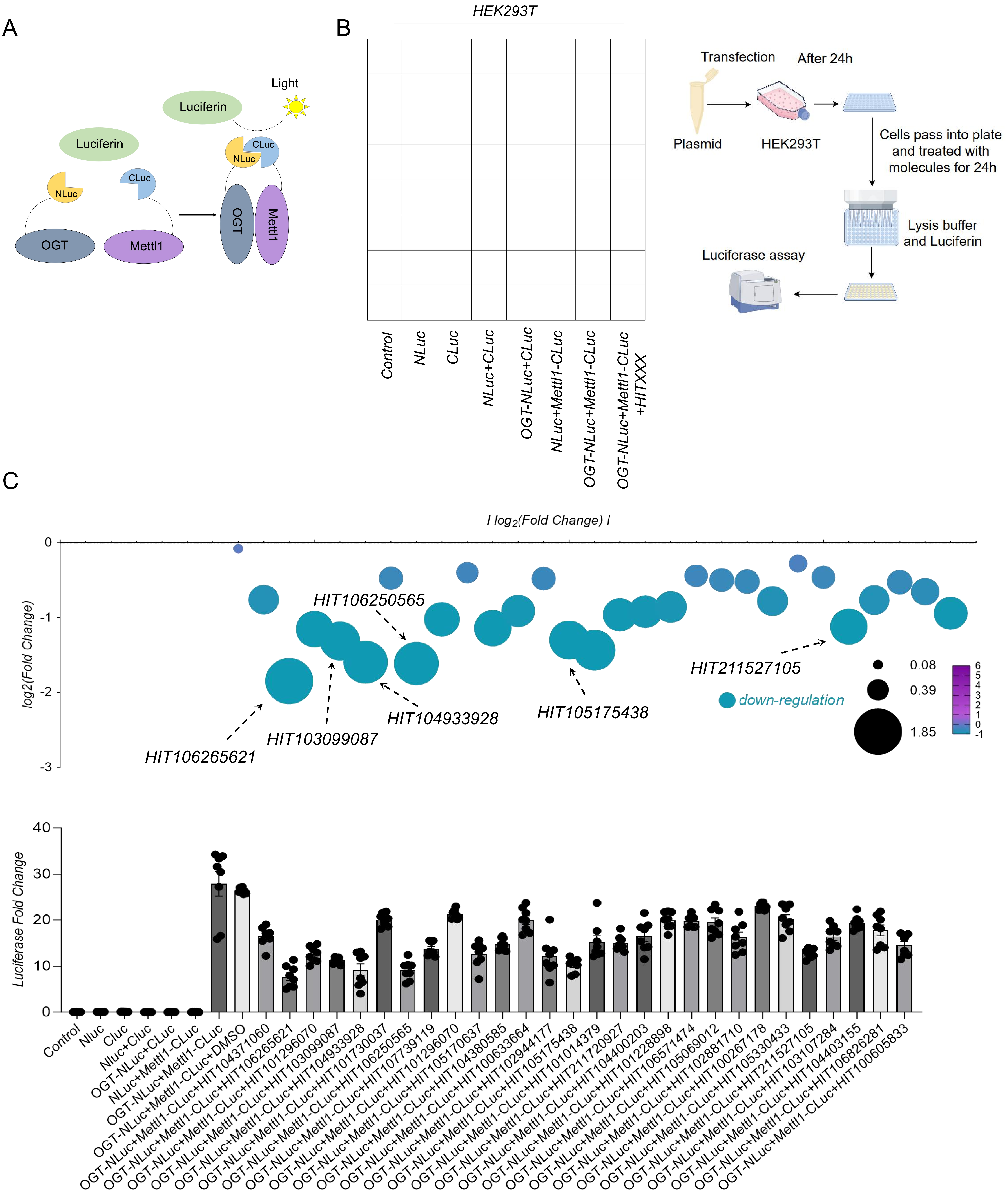

A
Luciferin
Light
NLuc
CLuc
OGT
Mettl1
B
HEK293T
Control
NLuc
CLuc
NLuc+CLuc
OGT-NLuc+CLuc
NLuc+Mettl1-CLuc
OGT-NLuc+Mettl1-CLuc
OGT-NLuc+Mettl1-CLuc +HITXXX
Transfection
After 24h
Plasmid
HEK293T
Cells pass into plate and treated with molecules for 24h
Lysis buffer and Luciferin
Luciferase assay
C
I log2(Fold Change) I
log2(Fold Change)
0
-1
-2
-3
HIT106250565
HIT106265621
HIT103099087
HIT104933928
HIT105175438
HIT211527105
down-regulation
0.08
0.39
1.85
6
5
4
3
2
1
0
-1
Luciferase Fold Change
40
30
20
10
0
Control
Nluc
Cluc
Nluc+Cluc
OGT-NLuc+CLuc
NLuc+Mettl1-CLuc
OGT-NLuc+Mettl1-CLuc
OGT-NLuc+Mettl1-CLuc+DMSO
OGT-NLuc+Mettl1-CLuc+HIT104371060
OGT-NLuc+Mettl1-CLuc+HIT106265621
OGT-NLuc+Mettl1-CLuc+HIT101296070
OGT-NLuc+Mettl1-CLuc+HIT103099087
OGT-NLuc+Mettl1-CLuc+HIT104933928
OGT-NLuc+Mettl1-CLuc+HIT101730037
OGT-NLuc+Mettl1-CLuc+HIT106250565
OGT-NLuc+Mettl1-CLuc+HIT107739119
OGT-NLuc+Mettl1-CLuc+HIT101296070
OGT-NLuc+Mettl1-CLuc+HIT105170637
OGT-NLuc+Mettl1-CLuc+HIT104380585
OGT-NLuc+Mettl1-CLuc+HIT100633664
OGT-NLuc+Mettl1-CLuc+HIT102944177
OGT-NLuc+Mettl1-CLuc+HIT105175438
OGT-NLuc+Mettl1-CLuc+HIT101014379
OGT-NLuc+Mettl1-CLuc+HIT211720927
OGT-NLuc+Mettl1-CLuc+HIT104400203
OGT-NLuc+Mettl1-CLuc+HIT101228898
OGT-NLuc+Mettl1-CLuc+HIT106571474
OGT-NLuc+Mettl1-CLuc+HIT105069012
OGT-NLuc+Mettl1-CLuc+HIT102881710
OGT-NLuc+Mettl1-CLuc+HIT100267178
OGT-NLuc+Mettl1-CLuc+HIT105330433
OGT-NLuc+Mettl1-CLuc+HIT211527105
OGT-NLuc+Mettl1-CLuc+HIT103107284
OGT-NLuc+Mettl1-CLuc+HIT104403155
OGT-NLuc+Mettl1-CLuc+HIT106826281
OGT-NLuc+Mettl1-CLuc+HIT100605833

Figure S21

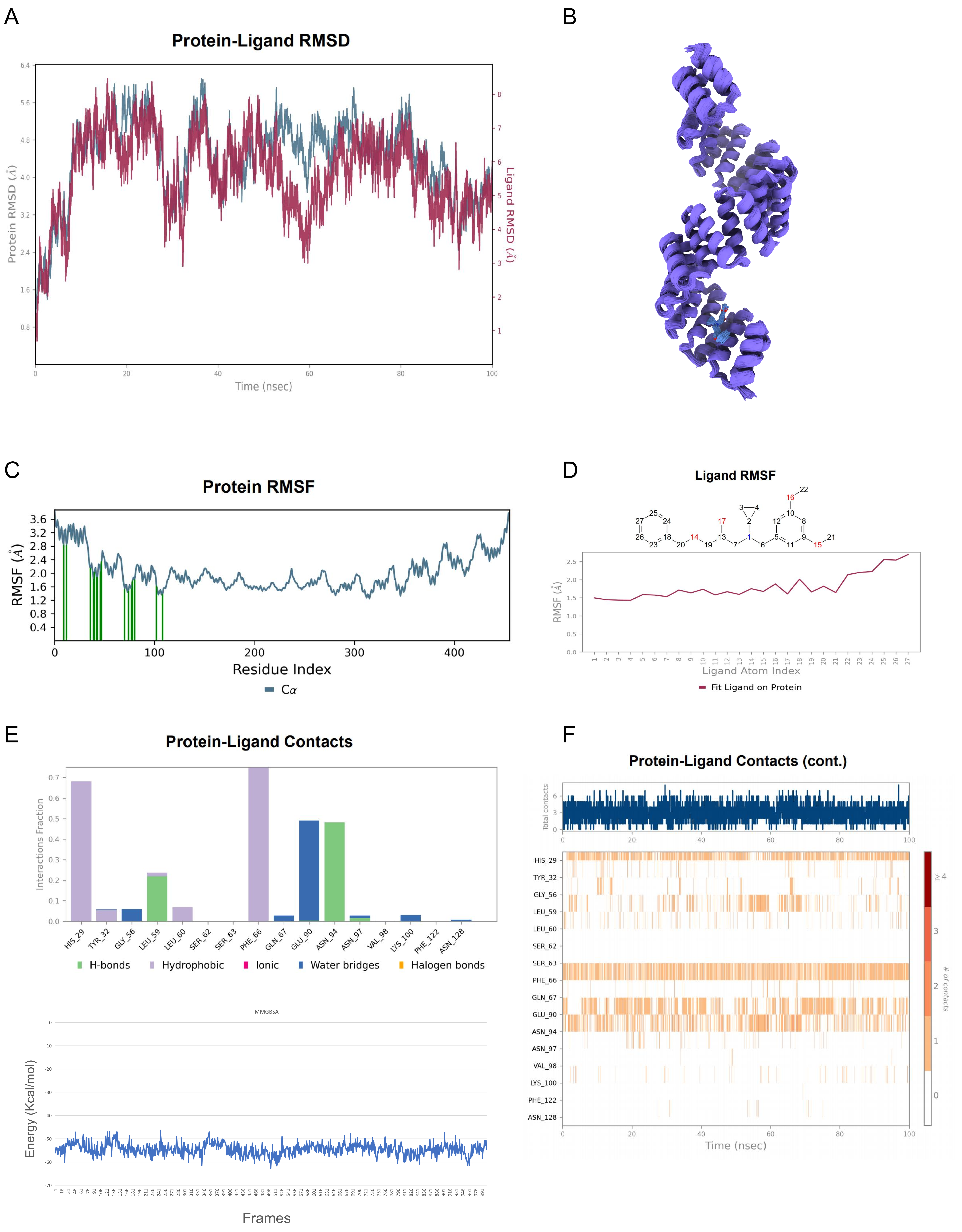

A
Protein-Ligand RMSD
Protein RMSD (Å)
Ligand RMSD (Å)
Time (nsec)
B
C
Protein RMSF
RMSF (Å)
Residue Index
Cα
D
Ligand RMSF
RMSF (Å)
Ligand Atom Index
Fit Ligand on Protein
E
Protein-Ligand Contacts
Interactions Fraction
HIS_29
TYR_32
GLY_56
LEU_59
LEU_60
SER_62
SER_63
PHE_66
GLN_67
GLU_90
ASN_94
ASN_97
VAL_98
LYS_100
PHE_122
ASN_128
H-bonds
Hydrophobic
Ionic
Water bridges
Halogen bonds
MMGBSA
Energy (Kcal/mol)
Frames
F
Protein-Ligand Contacts (cont.)
Total contacts
### of contacts
Time (nsec)

Figure S22

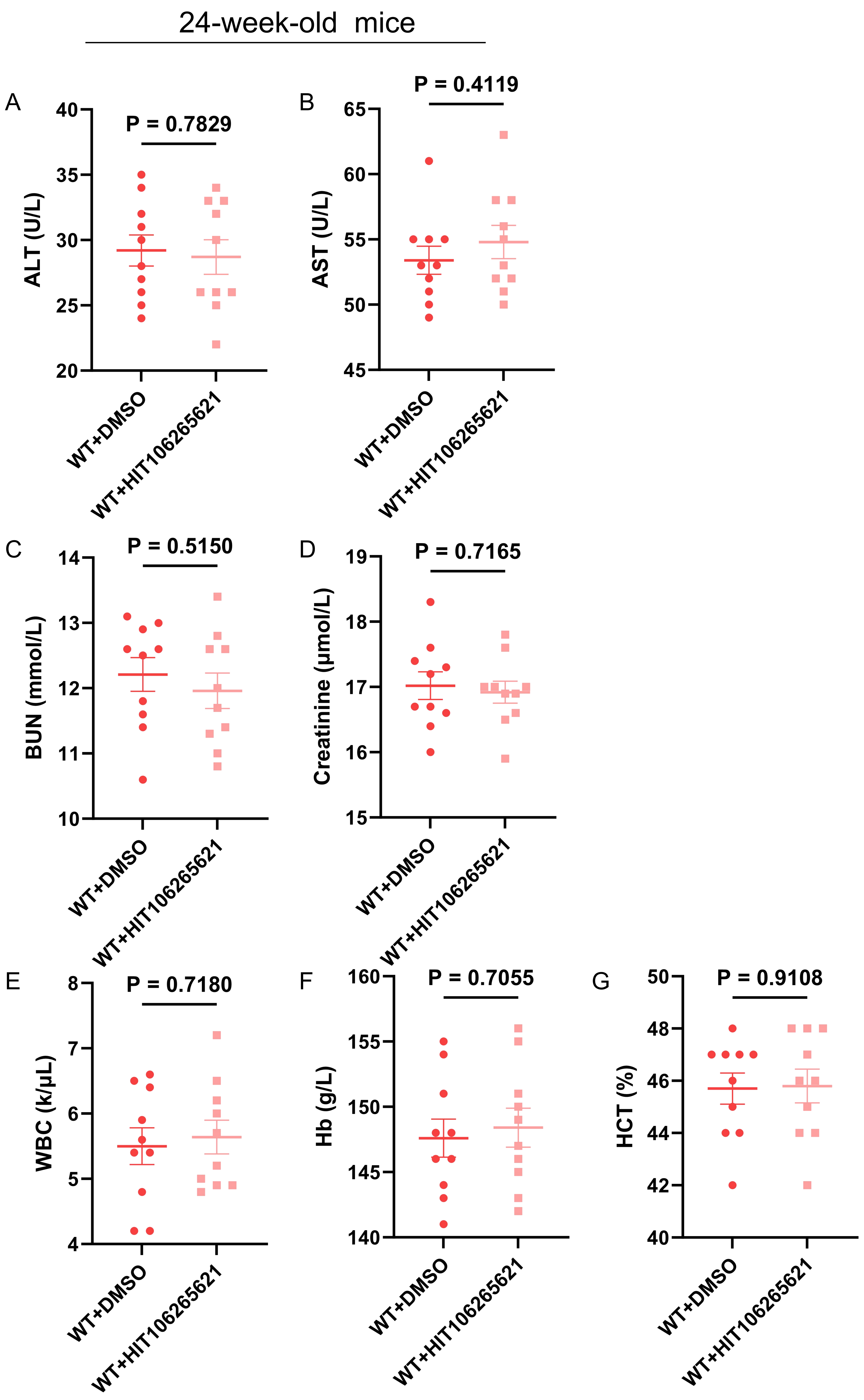

24-week-old mice
A
ALT (U/L)
P = 0.7829
B
AST (U/L)
P = 0.4119
C
BUN (mmol/L)
P = 0.5150
D
Creatinine (μmol/L)
P = 0.7165
E
WBC (k/μL)
P = 0.7180
F
Hb (g/L)
P = 0.7055
G
HCT (%)
P = 0.9108
WT+DMSO
WT+HIT10626562l

#### Materials and Methods

##### Mice

Mice were maintained with free access to food and water and housed in the specific pathogen-free (SPF) condition or germ-free (GF) condition. The conventionally-fed SPF mice were housed at 4-6 mice per cage in a room maintained at 20-25°C, 50% relative humidity, and a 12-h light/12-h dark cycle with sterilized food and autoclaved tap water *ad libitum.* All mice were checked twice for sterility tests before use. The $Myh6^{CreERT2}$ mouse strain was obtained from the Cyagen company (Suzhou, China). METTL1-flox mice (Strain S-CKO-03718) at the background of B6 were constructed via Cyagen (Suzhou, China). B6-ob mice (Strain NO. T001461) and B6-db (Strain NO. T002407) were purchased from GemPharmatech (Nanjing, China). Corn oil containing tamoxifen at a concentration of 50 mg/kg was administered to conditional CKO mice via intraperitoneal injection for seven consecutive days to induce recombination. B6-ob/m mice was crossed with $METTL1^{fl/fl}Myh6^{CreERT}$ and induced with tamoxifen to get the strain of $METTL1^{fl/fl}Myh6^{CreERT}$ ob/ob mice. Wild type (WT) C57BL/6J mice were bred and maintained and purchased from GemPharmatech (Nanjing, China). All animal experiments were carried out according to the guidelines for animal care set by the Institute for Laboratory Animal Research of Nanjing Medical University. This study was approved by the Institutional Animal Care and Use Committee of Nanjing Medical University (Approval No. IACUC-2305044). Primer for the mice genotype were listed as below:

METTL1-flox:

F:5′-CATGTCAAGCCACTGGTCTATGTT-3′;

R:5′-GGAAACTGAGGGTCAGAAATGCT-3′;

ob/ob mice:

RFLP-F 5′-TGAGTTTGTCCAAGATGGACC-3′;

RFLP-R 5′-GCCATCCAGGCTCTCTGG-3′;

WtLep-F 5′-AATGACCTGGAGAATCTCC-3′;

Lepob-R 5′-GCAGATGGAGGAGGTCTCA-3′

$Myh6^{CreERT}$:

F:5’-TCTATTGCACACAGCAATCCA-3’

R:5’-CCAGCATTGTGAGAACAAGG-3’

**Clinical subject enrolment and blood sampling**

The study population consisted of 80 patients with diabetic cardiomyopathy and 80 healthy donors. The patients were recruited from inpatients in the Department of cardiology of Affiliated Hospital of Jiangnan University according to the following inclusion and exclusion criteria. Inclusion criteria included diagnosis of diabetes and had either symptoms of heart failure (New York Heart Association functional class II-IV) or cardiac systolic or diastolic dysfunction on echocardiography or NT-proBNP ≥ 125pg/ml. Systolic dysfunction was defined as left ventricular ejection fraction (LVEF) < 50% or left ventricular fraction shortening (LVFS) < 25% or left ventricular end diastolic dimension (LVEDD) > 55mm in men and > 50mm in women. Diastolic dysfunction was defined as E/A < 0.8 or E/e′> 13 or left atrial volume index (LAVI) > 34mL/$m^2$. Exclusion criteria: younger than 18, pregnant or breastfeeding women, tumour, blood disease, thyroid gland disorders, mental and cognitive dysfunction, dementia, serious kidney dysfunction (eGFR < 45 mL/min/1.73$m^2$), body mass index ⩾ 45 kg/$m^2$, uncontrolled hypertension (systolic blood pressure > 140 mm Hg or diastolic blood pressure > 90 mm Hg), clinically significant arrhythmia, hypertensive heart disease, valvular diseases, coronary atherosclerotic heart disease, congenital heart diseases, infiltrative (e.g., amyloidosis, sarcoidosis, or hemochromatosis) cardiomyopathy, post-partum cardiomyopathy, or hypertrophic cardiomyopathy, pulmonary arterial hypertension, autoimmune diseases, chronic infectious or inflammatory diseases, history of toxic and unable to cooperate with follow-up. Besides, 80 healthy donors of the same age were enrolled as controls. Baseline clinical characteristics of patients and healthy donors were listed in **Table S1** in the *Supplemental Material*. Peripheral blood samples from patients and healthy donors were collected when they were eligible for enrollment. Then, serum and PBMCs were separated from peripheral blood for further ELISA detection. After one-year follow-up, peripheral blood samples were collected again to separate serum and PBMCs to repeat above detection.

The sample size after one-year follow-up is calculated by G*power. Using an effect size of 0.8, α = 0.05, and an independent sample t-test, the power analysis indicated that a sample size of 64 patients or healthy donors per group would be necessary to achieve 80% power. Considering that the mortality and drop-out rate of patients with heart failure at one-year follow-up is about 20%, 80 patients with diabetic cardiomyopathy and 80 healthy donors were recruited at the beginning of the study. Actually, after one-year follow-up, 66 patients with diabetic cardiomyopathy and 73 healthy donors were still at follow-up and accepted blood collection. This study was approved by Ethics Committee of Affiliated Hospital of Jiangnan University (IEC approval WXSY-YXLL-AF/SC-11/02.0; No. LS2024267) and all subjects signed informed consent forms.

**Diabetic cardiomyopathy mice model**

We constructed the diabetic cardiomyopathy mice model as the previous research[1]. Briefly, 6-week-old METTL1$^{fl/fl}$Myh6$^{CreERT}$ and METTL1$^{fl/fl}$Myh6$^{CreERT}$ ob/ob were injected with tamoxifen seven consecutive days to get the METTL1$^{CKO}$ mice. Then 8-week-old METTL1$^{CKO}$ and METTL1$^{Cre}$ mice were used to established the diabetic cardiomyopathy mice model. Streptozotocin (STZ, Sigma-Aldrich, Shanghai, China) was dissolved in sodium citrate buffer (pH = 4.5) to obtain a final concentration of 8 mg/ml 15 minutes before use. Male C57BL/6J mice were starved overnight and then injected intraperitoneally with 40 mg/kg STZ for five consecutive days. An equivalent volume of sodium citrate buffer was administered into the control animals as suggested. Blood glucose levels in the tail-veil of all experimental mice were assessed 1 week after STZ treatment, using a blood glucometer (OneTouch Verio Vue, OneTouch, Shanghai, China), with a range of ≥ 13.3 mmol/L to be considered as diabetic. After the STZ inducing, the mice were fed with the high fat diet to maintain. The body weight and blood glucose levels in the tail-veil were measured once a week after the successful model establishment.

**Cell culture and transfection**

Human cardiomyocyte-like AC16 cells and HEK-293T cells were purchased from ATCC. Cell lines were maintained in H-DMEM supplemented with 10% FBS at 37 °C in a 95% air and 5% CO2 atmosphere. For high glucose treatment, AC16 cells were incubated with normal (5 mM) or high (33 mM) glucose for different time points and then collected. siRNAs (si-OGT, si-METTL1) and relative controls were performed using Lipofectamine 2000 (Invitrogen, Carlsbad, CA), according to the manufacturer's recommendations. For siRNA transfection, complexes containing an RNA (pmol) to Lipofectamine 2000 (μl) ratio of 100:3 were prepared. The medium was then changed 6 hours after transfection.

**Preparation of isolated cardiac myocytes**

Primary cardiomyocytes were isolated from adult mice as described previously[2]. Briefly, 8-week-old mice fresh hearts were immersed into Tyrode's buffer containing NaCl 125mmol/L, KCl 5mmol/L, hydroxyethyl piperazine ethanesulfonic acid (HEPES) 15 mmol/L, $MgCl_2$ 1.2mmol/L, and glucose 10 mmol/L, pH 7.35-7.38. The aorta was then quickly mounted on the cannula. The hearts were perfused with perfusion buffer (Tyrode buffer supplemented with taurine 10 mmol/L and 2,3-Butanedione monoamine 10mmol/L) for 5 min at 37 °C. The perfusion buffer was then switched to enzymatic buffer (perfusion buffer supplemented with 1 mg/ml collagenase II). After perfusion and digestion, left ventricles were harvested and dissected into small pieces to dissociate in a transfer buffer. The digestion was stopped using perfusion buffer containing 1% BSA. The suspension was centrifuged at 500g (3 min) for cardiomyocytes precipitation. The cell pellet was checked for typical rod-shaped morphology.

For isolation of neonatal mouse cardiomyocytes (NMCM), we performed as previous described[3]. Neonatal mice (P1-3) were anesthetized on ice, and hearts were excised. Ventricles were transferred to a C-tube (Miltenyi Biotec) without mechanical disruption and tissues were incubated in a 37°C water bath with intermittent agitation (every 15-20 min, 3 cycles) if the dissociator was unavailable. Digestion was terminated immediately with pre-warmed DMEM supplemented with 10% fetal bovine serum (FBS) to prevent over-digestion. The cell suspension was filtered through a 100-μm strainer, centrifuged (300 ×g, 5 min), treated with red blood cell lysis buffer (3-5 min), and gently resuspended in DMEM + 10% FBS to minimize mechanical damage. To purify cardiomyocytes (CMs), the cell mixture was plated in culture flasks (37°C, 5% $CO_2$, 90 min) to allow non-CM attachment. CM-enriched suspensions were collected, while adherent non-CMs were retained for further analysis. Critical precautions included avoiding vigorous pipetting or vortexing and scaling reagents proportionally (max 20 hearts per 2.5 mL enzyme mixture).

**Mouse cardiac fibroblasts culture**

Mice were euthanized with pentobarbital sodium (100 mg $kg^{-1}$) intraperitoneally, and their hearts were separated with blunt dissection aseptically and rinsed three times in PBS (0.01 mM $PO_4^{3-}$, pH7.4) containing 200 U $ml^{-1}$ penicillin and 200 μg $ml^{-1}$ streptomycin (Gibco, Grand Island, NY, USA). Left ventricle of heart was minced and digested for 30 min in Dulbecco's Modified Eagle Medium (DMEM) (Gibco) containing 1 mg $ml^{-1}$ collagenase D (0.30 U $mg^{-1}$ lyophilisate, Roche Diagnostics GmbH, Mannheim, BW, Germany) and 2% fetal bovine serum (FBS) (v/v) (Gibco) on a constant temperature shaker at 37°C at 210 rpm. The post-digestive product was centrifuged at 1,500 rpm for 5 min. Supernatants were discarded and pellets were washed with PBS three times and centrifuged. Pellets of cardiomyocytes were resuspended in 3ml DMEM containing 15% (v/v) FBS, 100 U $ml^{-1}$ penicillin and 100 μg $ml^{-1}$ streptomycin and placed in six-well plates and kept in a humidified 5% $CO_2$ incubator at 37°C. The adherent fibroblast-shaped cells were mouse cardiac fibroblasts (MCFB).

**Color doppler echocardiography**

Mice were anesthetized by inhaling isoflurane at a 1:1 concentration with oxygen (4% induction concentration, 1.5% maintenance concentration) and detected by the high-frequency ultrasound imaging system (Vevo 2100, Visual Sonics, Toronto,

Canada) equipped with a 30-MHz transducer as described previously[4]. Left ventricular ejection fraction (LVEF) and left ventricular shortening fraction (LVFS) of each mouse were calculated as previously described[4].

**Adeno associated virus system**

For overexpressing different subcellular forms of METTL1 in mouse heart, the adeno associated virus (AAV9) delivery system was used to deliver AAV9-cTNT-METTL1, AAV9-cTNT-METTL1(T262A) and control viruses (Vigene Biosciences Co., Ltd., Shandong, China). In brief, mice were administrated 100 μL of virus containing 1 × $10^{11}$ AAV9 vector genomes containing AAV9-cTNT-METTL1 or AAV9-cTNT-METTL1 (T262A) via tail vein once a month.

**Wheat germ agglutinin (WGA) staining**

Mice were killed to determine the parameters of hypertrophic growth and cardiac remodeling. The hearts of the mice were excised and embedded in OCT compound according to standard histological protocols. The hearts were sectioned transversely at 5μm. Cardiomyocytes cross-sectional areas were detected via Alexa Fluor 594-conjugated wheat germ agglutinin (WGA, Invitrogen) staining. 4, 6-Diamidino-2-phenylindole (DAPI) was used to label the nuclei. The cardiac myocyte cross-sectional areas were measured using Image-Pro Plus software (version 6.0) with captured images.

**Histology and HE staining**

Mice were euthanized with pentobarbital sodium (100 mg $kg^{-1}$) intraperitoneally. Heart samples from mice or humans were fixed in periodate-lysine-paraformaldehyde (PLP) solution overnight at 4°C. Mouse heart samples were embedded in paraffin, and cut into 5-μm samples along the coronal plane using a rotary microtome (Leica Biosystems Nussloch GmbH, Nussloch, Germany). Sections were deparaffinized and rehydrated in a series of descent graded ethanol solutions and $ddH_2O$ for the following staining. Human or mouse frozen heart samples were embedded in

Tissue-Tek O.C.T. Compound (#4583, Sakura Finetek Japan Co., Ltd., CA, USA) and cut into 8-μm samples using a freezing microtome (Thermo Scientific Cryotome FSE Cryostats, Loughborough, Leicestershire) and then fixed in PLP solution overnight at 4℃. For HE staining, the slides of heart sections of mice were subjected to hematoxylin and eosin staining for morphological analyses.

**Masson's trichrome staining**

After deparaffinization and rehydration, the paraffin sections were stained according to instructions from the kit supplier (#KGMST-8003, KeyGen Biotech. Co. Ltd., Nanjing, Jiangsu, China). Fibrosis was measured as a positively stained area and was expressed as a percentage of the total area using Image-Pro Plus software (version 6.0).

**Western blotting**

Cells and mouse left ventricular tissues were lysed on ice with RIPA lysis buffer, protease and phosphatase inhibitors added. Lysates were centrifuged for 12,000 rpm, 15 minutes at 4℃; and the supernatants were used for western blotting. Proteins were separated by SDS-PAGE on 8%-15% acrylamide gels and transferred to polyvinyl difluoride membranes. Primary antibodies against the following proteins were used: β-actin, NPPA, NPPB, IL-1β, IL-6, TNF-α, METTL1, Collagen I, α-SMA, NLRP3, Puromycin, O-GlcNAc, OGT, Ubiquitin, Ubiquitin (linkage-specific K48), Ubiquitin (linkage-specific K63), DYKDDDDK Tag, HA Tag, Myc Tag and His Tag. Secondary antibodies were anti-rabbit and anti-mouse IgG HRP-conjugated antibodies. Chemiluminescence was generated and detected by ChemiDoc+(Bio-RAD). Protein levels were quantified using ImageJ software (version 1.44). All information about the primary antibodies used in the study were listed in **Table S2** in the *Supplemental Material*.

**Quantitative Real-Time PCR analysis**

Quantitative Real-Time PCR analysis was performed as described[5]. Total RNA was extracted using RNAiso Plus (Takara, 9109), followed by synthesizing cDNA with HiScript® II Q RT SuperMix (Vazyme). Gene expression levels were detected using target primers with ChamQTM SYBR® qPCR Master Mix (Vazyme). The expression of target genes was normalized to U6 or β-actin mRNA. The sequences of the primers are listed in **Table S3** in the *Supplemental Material.*

**Co-Immunoprecipitation (Co-IP)**

Co-IP was performed with Pierce® Co-Immunoprecipitation Kits (Pierce Biotechnology, Rockford, IL, USA) as previously described[6]. Cell lysates from AC16, NMCM or 293T cells were incubated with the corresponding IgG control or precipitation antibodies at 4°C overnight, including METTL1 (ATLAS, USA), USP3 (Cell Signaling Technology, USA), HA (Cell Signaling Technology, USA), DYKDDDDK Tag, His Tag for Co-IP analyses. Co-Immunoprecipitation or total cytoplasm lysates were detected in western blots.

**Plasmid or adenovirus construction and transfection**

Adenovirus for overexpression of Flag-METTL1 or HA-METTL1 (human or mouse) was synthesized by TranSheep Bio, Shanghai, China. According to the structural features, METTL1 overexpression plasmids containing a full-length fragment and His Tag USP5 were constructed with pcDNA3.1/ His-tag (Clontech, USA). Plasmids were transfected into HEK293T or AC16 cells using Lipofectamine 2000 (Invitrogen Inc. USA) as previously described[6].

According to the O-GlyNac threonine of human METTL1 or mouse METTL1, we generated adenovirus for overexpression of three mutation plasmids: T to A mutation Flag-METTL1 adenovirus at threonine 58, 268 and 269 and T to A mutation Flag-METTL1 adenovirus at threonine 262 for mouse.

***In vivo* ubiquitin detection**

Plasmids for overexpression of Flag-Tag METTL1, HA-Ubiquitin, HA-K6-Ubiquitin, HA-K11-Ubiquitin, HA-K27-Ubiquitin, HA-K29-Ubiquitin, HA-K33-Ubiquitin,

HA-K48-Ubiquitin, HA-K63-Ubiquitin were transfected into 293T cells. The 293T cells were incubated in complete medium after transfection mixture for 6h, and were harvested after 24h. MG132 (50μM) was incubated in complete medium for 6h. The cell lysis was used for Co-IP assay to detect the ubiquitin of METTL1. For AC16 and NMCM cells, the cells were treated with the high glucose (33mM) for 24h and induced the cells with or without MG132 for 6h, cells were then collected for the Co-Immunoprecipitation (Co-IP) assay to detect the *in vivo* ubiquitin level. Production of the above plasmids was performed by TranSheep Bio, Shanghai, China.

**Detection of half-life of METTL1**

AC16 or NMCM cells were transfected with the HA-tag METTL1 (WT) and METTL1 (T268A) for human METTL1 and METTL1 (T262A) for mouse METTL1. Then we treated with Cycloheximide (CHX, MCE) with final concentration 50μM for 0h, 3h, 6h and 12h. AC16 or NMCM cells were collected for protein extraction for further western blot.

**Atomic modeling analysis**

**3D structure prediction of proteins**

The structures of OGT in complex with METTL1 were predicted using the AlphaFold3 multimer programs. The conformation with the highest score was used for dynamic simulation or analysis in AlphaFold3 according to previous research[7]. Accurate structure prediction of biomolecular interactions with AlphaFold 3. Nature (2024).

**Molecular dynamics simulation**

GROMACS package (version 2023.03) was applied to run conventional MD simulations to investigate the changes in conformation of OGT-METTL1 complexes. The force field amber14sb were employed to parameterize proteins, respectively. The TIP3P was used for the waters. This OGT-METTL1 complexes was solvated in an octahedral water box, and then the charge of the system was neutralized by adding

0.150 M chloride and sodium ions. First, the steepest descent minimization method was used to minimize the energy of the system by 50,000 steps. In the next step, we restricted the position of heavy atoms to run both NVT equilibration and NPT equilibration by 50,000 steps. The system temperature was maintained at 300 K, and the system pressure was maintained at 1 bar. Upon completion of the two equilibration phases, the system was now well-equilibrated at the desired temperature and pressure. A 100-ns unrestrained simulation was carried out. Every 20 ps, the energy and coordinate system of the trajectory was saved. In the simulation trajectory, ChimeraX were used to map interaction patterns and animate kinetic trajectories.
GROMACS: https://www.sciencedirect.com/science/article/pii/S2352711015000059
amber14sb: https://pubs.acs.org/doi/full/10.1021/ja5032776
tip3p: http://refhub.elsevier.com/S0223-5234(20)30971-5/sref37
ChimeraX: https://onlinelibrary.wiley.com/doi/full/10.1002/pro.3235

**Free energy calculations and residue decomposition**

MM-GBSA method has been widely adopted in the estimation of binding free energy in drug research. In our work, the MM-GBSA calculation was performed using the gmx_MMPBSA, a tool of GROMACS for MM-PB (GB) SA calculations. To understand the binding of OGT and METTL1 at the molecular level, we used gmx_MMPBSA to decompose the free energy of binding to the contribution of each residue to the free energy of binding.
MM-GBSA: https://www.tandfonline.com/doi/full/10.1517/17460441.2015.1032936
gmx_MMPBSA: https://pubs.acs.org/doi/abs/10.1021/acs.jctc.1c00645
The contributions are further broken for the complex, receptor and ligand into GGAS and GSOLV. GGAS is the interaction energy and is obtained after sum the internal(bonded) components (BOND + ANGLE + DIHED) and the non-bonded (VDWAALS + EEL) components. For GSOLV, the polar and non-polar contributions are EGB (or EPB) and ESURF (or ENPOLAR + EDISPER), respectively for GB (or PB) calculations. A single trajectory protocol does not produce any differences between bond lengths, angles, dihedrals or 1-4 interactions between the complex and

receptor/ligand structures. Thus, when subtracted they cancel completely. If not, these values are displayed and inconsistency warnings are printed. When this occurs, the results are generally useless. Of course, this does not hold for the multiple trajectory protocol as independent trajectories are used for the complex, receptor and ligand. Two approaches are used when calculating the standard deviation, and the standard error of the mean. The SD and SEM values are calculated using a sample (array) of values. On the other hand, SD (Prop.) and SEM (Prop.) are obtained with the propagation of uncertainty formula for f = A - B. Check this thread for more details on MM/PB(GB)SA statistics.

**High-throughput virtual screening of a commercial compound library for novel OGT-METTL1 inhibitors**

**Protein structure preparation and preprocessing**

The protein structures were first preprocessed using the Protein Preparation Wizard module in Schrödinger. Potential binding sites of OGT and METTL1 were then predicted using Schrödinger's SiteMap module. Subsequently, docking grids were generated using the Glide Grid module in Schrödinger, which served as the basis for subsequent molecular docking studies.

**Small molecule database preparation**

The screening database was sourced from ChemDiv (https://www.chemdiv.com/) and processed using Schrödinger's LigPrep module. This preprocessing included protonation, desalting, hydrogen addition, generation of tautomers and stereoisomers, and energy minimization.

**Virtual screening**

Initial ADMET filtering of the database molecules was performed using Schrödinger's QuickProp module, retaining only compounds that satisfied Lipinski's Rule of Five and excluded reactive fragments. High-throughput virtual screening (HTVS) was then conducted using Schrödinger's Glide module. The top 10% of molecules based on

HTVS scores were retained for subsequent standard precision (SP) docking. Similarly, the top 10% of molecules from SP docking were subjected to extra precision (XP) docking. The top 10% of molecules from XP docking were further analyzed using MM-GBSA to calculate binding free energies. Based on MM-GBSA scores, five molecules were selected for binding mode analysis.

**Molecular dynamics (MD) simulation analysis**

To further optimize the binding mode of OGT with the top-scoring small molecule, molecular dynamics (MD) simulations were performed using the Desmond program. The OPLS2005 force field was applied for parameterization of the protein and small molecule, while the TIP3P model was used for water molecules. The protein-ligand complex was placed in a cubic water box and solvated, with 0.150 M sodium and chloride ions added to neutralize the system. Energy minimization was performed using the steepest descent method for 50,000 steps, followed by 50,000 steps of NVT and NPT equilibration with heavy atom positional restraints. The system temperature and pressure were maintained at 300 K and 1 bar, respectively. After the equilibration phases, an unconstrained 100 ns simulation was conducted, with energy and coordinates saved every 10 ps.

**DUB siRNA library**

Human DUB siRNA library was purchased from Dharmacon as previous research[8]. We screened a human DUB siRNA library consisting of siRNAs targeting each of the 98 DUB genes according to previous research[8]. Pooled siRNAs (4 siRNAs per gene) were transfected into HEK293T cells, and protein lysates were collected 48h later for further detection of METTL1 in cells via METTL1 ELISA kit (RayBiotech). The fold change of the METTL1 expression level in siRNA groups and control group were calculated. AC16 were then transfected with the top 10 siRNA in HEK293T for 24h and treated for HG for 24h, the AC16 cells were then collected for the detection the level of METTL1 via WB assay.

**m7G methylated RNA immunoprecipitation (MeRIP)-qPCR**

The utilized protocol for m7G-MeRIP was previously outlined[9]. Total RNA was isolated from cells or their corresponding controls under HG stimulation. The total RNA was sonicated into 100-nt to 150-nt fragments RNA and then incubated with m7A antibody to confirm m7A enrichment of the RNA by performing qRT-PCR or high-throughput assays. Briefly, fragmented RNA and 5 μg of m7A antibody or rabbit IgG were combined in 1 mL RIP immunoprecipitation buffer (100 μL of the supernatant, 860 μL of RIP Wash Buffer, 35 μL of 0.5 M EDTA, and 5 μL RNase inhibitor for each reaction) and incubated with shaking at 4°C overnight. m6A IP protease was incubated in proteinase K buffer at 55°C for 30 min with shaking. Trizol was used to extract RNA, and m7G immunopurification of mRNA was performed using the sample for detection using RT-PCR.

**RNA stability assay**

AC16 Cells were cultured in 6-well plates and transfected with the specified constructs as outlined above. Following a 24h transfection period, the cells were exposed to actinomycin D (Act D, 10 μg/mL, Cat# GC16866, GLPBIO) for 0, 3, or 6 min before being collected. Total RNA was then isolated for qRT-PCR analysis.

**Seahorse analysis**

According to previous research[10], cells were seeded in a 96-well Seahorse XF Cell Culture Microplate ($2 \times 10^4$ cells/well, Agilent, USA). Following overnight incubation, the culture medium was replaced as per the instructions provided in the Agilent Seahorse XF Cell Mito Stress Test Kit. Subsequently, the microplate was incubated in a non-CO2 incubator at 37 °C for 1 h prior to measuring the oxygen consumption rate (OCR) in the XF Analyzer, following the procedural guidelines.

**Fatty acid oxidation (FAO) complete assay**

The utilized protocol for FAO assay was previously outlined[11]. FAO was determined using a fatty acid complete oxidation assay kit from Abcam (ab222944). The assay was performed according to the manufacturer's protocol. This assay uses FAO substrate, Oleate conjugated with BSA, and FAO modulator FCCP. In the presence or the absence of FAO substrate, Oleic acid, FAO can be determined. An equal number of cells or an equal amount of isolated mitochondria were incubated in a fatty acid measurement medium that contained 150μM Oleate conjugated with BSA and 0.5mM L-carnitine in a standard ELISA plate. Extracellular oxygen reagent was added to the mitochondria before sealing each well with highly sensitive mineral oil to limit the back diffusion of oxygen. This assay was designed based on the principle that molecular oxygen quenches the oxygen consumption reagent. Therefore, the increase in fatty acid-driven oxygen consumption is monitored for one hour as an increase in fluorescence at oxygen absorption spectra (Excitation and Emission at 380-650 nM). Basal FAO in mitochondria was measured in the presence of carnitine without Oleate.

**Protein Synthesis Assay**

The Click-iT HPG Alexa Fluor 488 Protein Synthesis Assay Kit was purchased from Thermo Fisher Scientific(C10428). The cells were cultured and treated with HG for 24h and follow the instructions to detect the Protein Synthesis level.

**Immunofluorescence assay**

For the immunofluorescence assay as the previous described[12], AC16 cells subjected to different treatments were washed three times with PBS, fixed in 4% paraformaldehyde (pre-cooled at 4 °C, Solarbio) for 20 min, and blocked with 5% bovine serum albumin (BSA, KeyGEN BioTECH) containing 0.1% TritonX-100 (Sigma) for 30 min at room temperature. Then, the resultants were incubated with anti-METTL1 and anti-O-GlyNac or anti-USP5 antibodies. After washing with 1 × PBS again, and then incubated with fluorescently conjugated secondary antibodies (1:200, Thermo Fisher) diluted in 5% BSA for about 1h at room temperature. Finally, the cells were washed with 1 × PBS and mounted with the DAPI-containing mounting medium (Solarbio). The images of cells were taken with a laser scanning confocal microscope.

**Luciferase reporter assay for drug screening**

HEK293T cells in 24-well plates were transfected with luciferase reporter vector fusing OGT-NLuc and METTL1-CLuc. All cells were treated with different small molecules and collected 48 h after treatment, and the luciferase activities in each well were calculated by the luciferase reporter gene analysis system (Promega). The relative ratio of luciferase activity was determined. Each experiment was repeated three times.

**Molecular docking prediction**

In this study, the three-dimensional structures of both OGT and USP5 proteins were derived from predictive models generated by the AlphaFold 3 online tool (https://alphafoldserver.com). Protein–protein molecular docking was performed using the online tool HDOCK SERVER, with detailed parameters referring to the provided example tutorials[13]. The Protein Interfaces, Surfaces and Assemblies (PISA) service, available at the European Bioinformatics Institute, can be accessed via the following link: http://www.ebi.ac.uk/pdbe/prot_int/pistart.html. The final results were analyzed and visualized using PyMOL version 2.6 software.

**Untargeted metabolomics analysis**

Untargeted metabolomics analysis was performed on an ultra-performance liquid chromatography tandem quadrupole time-of-flight mass spectrometer (UPLC-Q-TOF/MS) (Waters, London, UK), and the large volume of data was processed using MetaX software. All samples were acquired by the LC-MS system. First, all chromatographic separation was applied using an UPLC system (Waters). An ACQUITY UPLC BEH C18 column (100mm × 2.1 mm, 1.7µm, Waters) was used for reversed-phase separation. The column oven was maintained at 50°C. The flow-rate was 0.4 ml/min and the mobile phase consisted of solvent A (water + 0.1% formic acid) and solvent B (acetonitrile + 0.1% formic acid). Gradient elution conditions were set as follows: 0-2 min, 100% phase A; 2-11 min, 0-100% B; 11-13 min, 100%

B; 13-15min, 0-100% A. The injection volume for each sample was 10 ml. A high-resolution tandem mass spectrometer (Synapt G2-Si Q-TOF, Waters) was used to detect the metabolites eluted from the column. The Q-TOF was operated in positive ion mode. The capillary and sampling cone voltages were 1 kV and 40 V, respectively. The mass spectrometry data were acquired in centroid MSE mode. The TOF mass range was from 50 to 1200 Da and the scan time was 0.2 s. For the MS/MS detection, all precursors were fragmented using 20-40 eV, and the scan time was 0.2 s. During acquisition, the low-energy signal was acquired every 3 s to calibrate the mass accuracy. Furthermore, in order to evaluate the stability of the LC-MS during the whole acquisition, a quality control sample (pool of all samples) was acquired after every 10 samples. The raw data from the mass spectrometry analysis were imported into the commercial software Progenesis QI (version 2.2) for peak extraction to acquisite metabolite related mass-to-charge ratios, retention times, and ion areas. Statistical analysis of the mass spectrometry data was performed using the metabolomics R package meta X, with metabolite identification based on the KEGG database. Serum metabolites passing the criteria of variable importance in the projection values R 1.0, fold-change R 2 or % 0.5, and false discovery rate-adjusted P Values ＜ 0.05 were selected as significantly different between groups as described elsewhere.

**HPLC fractionation and enrichment of O-GlcNAcylated peptides and Ubiquitylated peptides**

**Protein extraction and digestion**

Protein extraction : Urea (8M Urea，100mM Tris/HCl，pH 8.5) buffer was used for sample lysis and protein extraction heart tissues from 22-week-old ob/ob and ob/m mice. The amount of protein was quantified with the BCA Protein Assay Kit; SDS-PAGE: A number of 15 μg protein for each sample were mixed with 5X loading buffer respectively and boiled for 5 min. The proteins were separated on SDS-PAGE gel (4-20% precast gels, constant voltage 200V, 35 min). Protein bands were visualized by Coomassie Blue R-250 staining; In-solution Digestion: DTT (with the final concentration of 10 mM) was added to each sample, respectively, mixed at 600 rpm for 1.5 h (37 °C), then cooled to room temperature. Added IAA with the final concentration of 20 mM into the mixture， incubated in the dark for 30min. The concentrate of UA was diluted to 1.5M with 4 times volume of 50 mM Tris HCl (pH 8.0). Trypsin was added to the samples, the trypsin: protein (wt/wt) ratio was 1:50, incubated at 37 °C for 15-18 h (overnight). After overnight digestion, adjusted the pH to pH ≤ 3 with TFA. The digest peptides of each sample were desalted on C18 SPE Cartridges, and lyophilized for further use.

**O-GlcNAcylated peptides enrichment**

The samples were reconstituted in 1.4 mL of precooled IAP Buffer, added pretreated Anti GlcNAc-S/T antibody beads (PTMScan® O-GlcNAc [GlcNAc-S/T] Motif Kit，Cell Signaling Technology, then incubated at 4 ºC overnight，centrifuged at 2,000 ×g for 30s, then discarded the supernatant. Anti GlcNAc-S/T antibody beads were washed with 1mL precooled IAP Buffer for 3 times，than washed with 1mL precooled water for 3 times. 40 μL 0.15% TFA was added to the washed beads, incubated for 10 min at room temperature (mixed gently every 2-3 min), then added 0.15% TFA twice, centrifuged at 2,000 ×g for 30s, the supernatant was desalted by Zip Tips.

**Ubiquitylated peptides enrichment**

The samples were reconstituted in 1.4 mL of precooled IAP Buffer, added pretreated Anti-K-ε-GG antibody beads (PTMScan Ubiquitin Remnant Motif (K-ε-GG) Kit，Cell Signal Technology), then incubated at 4 ºC for 1.5 h，centrifuged at 2,000 ×g for 30s, then discarded the supernatant. Anti-K-ε-GG antibody beads were washed with 1mL precooled IAP Buffer for 3 times，than washed with precooled water for 3 times. Forty μl 0.15% TFA was added to the washed beads, incubated for 10 min at room temperature, then added 0.15% TFA again, centrifuged at 2,000 ×g for 30s, the supernatant was desalted by C18 STAGE Tips.

**LC-MS/MS analysis**

Each sample was separated by a NanoElute HPLC system at a nanoliter flow rate. The chromatographic column was balanced with 95% liquid A (0.1% Formic acid), and the samples were separated on automatic injector (AUR2-25075C18A-CSI, 25 cm x 75 μm ID, 1.6 μm C18 (IonOpticks, Australia) with a linear gradient of buffer B (84% acetonitrile and 0.1% Formic acid) at a flow rate of 300 nl/min. LC-MS/MS analysis was performed on a timsTOF Pro mass spectrometer (Bruker) that was coupled to Nanoelute (Bruker Daltonics) for 60 min. The mass spectrometer was operated in positive ion mode. The mass spectrometer collected ion mobility MS spectra over a mass range of m/z 100-1700 and 1/k0 of 1.16, and then performed 10 cycles of PASEF MS/MS with a target intensity of 1.5kV. Active exclusion was enabled with a release time of 0.4 minutes.

**Bioinformatic analysis**

Cluster 3.0 (http://bonsai.hgc.jp/~mdehoon/software/cluster/software.htm) and Java Treeview software (http://jtreeview.sourceforge.net) were used to performing hierarchical clustering analysis. Euclidean distance algorithm for similarity measure and average linkage clustering algorithm (clustering uses the centroids of the observations) for clustering were selected when performing hierarchical clustering. A heat map was often presented as a visual aid in addition to the dendrogram. For GO annotation, the protein sequences of the selected differentially expressed proteins were locally searched using the NCBI BLAST+ client software (ncbi-blast-2.2.28+-win32.exe) and InterProScan to find homologue sequences, then gene ontology (GO) terms were mapped and sequences were annotated using the software program Blast2GO. For KEGG analysis, the studied proteins were blasted against the online Kyoto Encyclopedia of Genes and Genomes (KEGG) database (http://geneontology.org/) to retrieve their KEGG orthology identifications and were subsequently mapped to pathways in KEGG. Enrichment analysis was applied based on the Fisher' exact test, considering the whole quantified proteins as background dataset. Benjamini- Hochberg correction for multiple testing was further applied to adjust derived p-values. And only functional categories and pathways with p-values under a threshold of 0.05 were considered as significant.

**Quantitative proteomics and data analysis**

Quantitative proteomics was performed as previous[14], according to the manufacturer's protocol. Firstly, total proteins were extracted from heart tissue of METTL1$^{Cre}$ob/ob and METTL1$^{CKO}$ob/ob mice and digested with trypsin. Then the protein samples labelled with TMT reagent, combined, fractionated, and split for quantitative analysis using UHPLC-MS/MS on an EASY-nLCTM 1200 UHPLC system (Thermo Fisher, Germany) coupled with an Q ExactiveTM HF-X (Thermo Fisher, Germany). Raw files are directly imported into Proteome Discoverer 2.5 software for database retrieval, peptide spectrum and protein quantification. The proteins whose quantitation significantly different between CKO and Cre groups, ($p < 0.05$ and $FC < 0.83$ or $FC > 1.5$), were defined as differentially expressed proteins (DEP). DEPs were used for Gene Ontology (GO) enrichment analysis using the inter pro scan program against the non-redundant protein database (including Pfam, PRINTS, ProDom, SMART, ProSite, PANTHER).

**m7G methylated RNA immunoprecipitation sequencing (m7G-MeRIP-seq)**

Total RNA was extracted from heart tissue from METTL1$^{Cre}$ob/ob and METTL1$^{CKO}$ob/ob mice using TRIzol reagent, followed by DNase I treatment to eliminate genomic DNA contamination. RNA integrity (RIN ≥ 8.0) and concentration were verified using an Agilent 2100 Bioanalyzer and Qubit 4.0 fluorometer, respectively. Purified RNA (1-5 μg) was fragmented into 100–300 nt fragments via incubation in $Mg^{2+}$-containing fragmentation buffer at 94°C for 15 min, followed by ethanol precipitation. For m7G-MeRIP, fragmented RNA underwent pre-clearing with Protein A/G magnetic beads (Thermo Fisher) to reduce nonspecific binding. Immunoprecipitation was performed overnight at 4°C with anti-m7G-specific antibody (MBL International, Cat# RN017M; 5ug/per sample) or control IgG, followed by 2-hour incubation with Protein A/G beads. Beads were washed four times with high-salt buffer (0.1% SDS, 1% Triton X-100), and bound RNA was eluted using

SDS/proteinase K buffer, purified via phenol-chloroform extraction, and ethanol-precipitated. Strand-specific libraries were prepared from immunoprecipitated (IP) and input RNA using the Illumina TruSeq kit, with library quality assessed via Agilent 2100 and qPCR. Sequencing was performed on the Illumina NovaSeq platform (PE150). Data analysis included quality control (FastQC, Trim Galore), alignment to the reference genome (hg38; HISAT2/Bowtie2), peak calling (MACS2/exomePeak), and functional annotation (DAVID/clusterProfiler). RNase-free conditions were maintained throughout. Antibody specificity was validated by pre-experimental optimization.

**Statistical Analysis**

For biochemical experiments, data analyses were performed using GraphPad Prism software 9 (Boston, MA, USA). For the data with normal distribution, Student's unpaired 2-tailed t test was used to compare two groups, and one-way ANOVA was used for studies with more than 2 groups. Two-way ANOVA was used for studies with two variables. All summary data are expressed as the mean ± SEM. At least 3 biologically experimental replicates were performed for each experiment unless indicated. For the clinical study, statistical analyses were performed using SPSS software (IBM Corp., New York, USA). Data are expressed as the mean ± SEM (normal distribution) or median (interquartile range (IQR): 25%-75%, non-normal distribution). The Shapiro-Wilk test was used to test for normal distribution. Between-group differences of variables were tested using unpaired Student's t test (data with normal distribution) and the Mann-Whitney U test (data with non-normal distribution). Within-group differences of variables from baseline to follow-up were tested using the paired Student's t test or the repeated measures ANOVA (data with normal distribution) and the Wilcoxon Matched-pairs Signed Rank Sum test or the Friedman test (data with non-normal distribution). Between-group differences of proportions were tested using the c2 test. Correlation coefficients are determined by Spearman r correlation coefficient. A p value < 0.05 was considered to be significant for all tests. ns, not significant; *, p < 0.05; **, p < 0.01; ***, p < 0.001; ****, p <

0.0001.

Figure S1

(A) Heatmap of mRNA expression profile in heart tissue of diabetic mice (GSE173384). (B) The expression pattern of the METTL1 protein in diabetic heart tissue in HPA database. (C) Expression of METTL1 in different cells of multiple organs. (D) Expression of METTL1 in different cells of heart, data were from HPA database. (E) Quantitative PCR was employed to assess METTL1 expression levels in various components of ob/ob mouse heart tissue following separation and extraction of mouse immune cells, including T cell sorting, cardiac tissue (CMs), cardiac fibroblasts (CFs), endothelial cells (ECs), bone marrow-derived macrophages (BMDMs). Data are expressed as mean ± SEM.

Figure S2

(A-B) Expression of METTL1 in different tissues and cells, data were from Tabula Muris database. (C) Expression of METTL1 in different cells of heart, data were from Tabula Muris database. (D) Expression of METTL1 in different cells, data were from Single Cell Portal database. (E) Primary cardiac fibroblasts from C57BL/6 wild-type mice (8-week-old) were cultured and treated with low (5mM) or high glucose (HG, 33mM) for 24h. Cells were collected for detection of protein level of METTL1 via WB assay. (F) BMDMs from C57BL/6 wild-type mice (8-week-old) were cultured and treated with HG for 24h. Cells were collected for detection of protein level of METTL1 via WB assay. (G) ECs from C57BL/6 wild-type mice (8-week-old) were cultured and treated with HG for 24h. Cells were collected for detection of protein level of METTL1 via WB assay. (H) AC16 cells were cultured and treated with HG for 24h and collected for detection of protein level of WDR4. (I) NMCM cells were cultured and treated with HG for 24h and collected for detection of protein level of WDR4. The expression levels of WDR4 were detected in heart tissues of ob/m and ob/ob mice (J), HFD + STZ mice (K) and PBMCs from diabetic cardiomyopathy patients and healthy donors (L). Data are expressed as mean ± SEM.

Figure S3

(A) Construction strategy of scheme-specific METTL1 knockout mice. Picture were downloaded from Cyagen Biosciences Inc. (B) Expression levels of METTL1 in various organs from $METTL1^{Cre}$ mice without tamoxifen (up) or after tamoxifen induced (below) for 5 days (75mg/kg) *in vivo*. (C) Body weight of $METTL1^{CKO}$ and $METTL1^{Cre}$ healthy mice after tamoxifen induced in different weeks. (D) Serum glucose levels of $METTL1^{CKO}$ and $METTL1^{Cre}$ healthy mice after tamoxifen induced in different weeks. (E) HE and Masson staining of heart tissues from $METTL1^{CKO}$ and $METTL1^{Cre}$ healthy mice 12 weeks after tamoxifen induction. (F) IPGTT assay of $METTL1^{CKO}$ and $METTL1^{Cre}$ healthy mice 12 weeks after tamoxifen induction. (G-I) Serum BNP, cTnI and EF levels of $METTL1^{CKO}$ and $METTL1^{Cre}$ healthy mice 12 weeks after tamoxifen induction. (J) Relative expression level of Col1α1, α-SMA, ANP, BNP, IL-1β, IL-6 and TNF-α in heart tissues from $METTL1^{CKO}$ and $METTL1^{Cre}$ healthy mice 12 weeks after tamoxifen induction. Data are expressed as mean ± SEM. (K-L) Heart rate, systolic and diastolic blood pressure of $METTL1^{CKO}$ and $METTL1^{Cre}$ healthy mice 12 weeks after tamoxifen induction. Data are expressed as mean ± SEM.

Figure S4

Schematic diagram of tamoxifen induced myocardial METTL1 knock-out mice model. Mice were dived into three groups: $MELLT1^{Cre}$ ($METTL1^{fl/fl}Myh6^{Cre}$ without tamoxifen induced), $METTL1^{CKO}$ ($METTL1^{fl/fl}Myh6^{Cre}$ with tamoxifen induced) and $METTL1^{CKO}$ + AAV9-$METTL1^{OE}$ mice and induced with HFD + STZ for 12 weeks. (A) Serum glucose level of $MELLT1^{Cre}$, $METTL1^{CKO}$ and $METTL1^{CKO}$ + AAV9-$METTL1^{OE}$ mice after HFD + STZ for 12 weeks. (B) Body weight of $MELLT1^{Cre}$, $METTL1^{CKO}$ and $METTL1^{CKO}$ + AAV9-$METTL1^{OE}$ mice after HFD + STZ for 12 weeks. (C) IPGTT assay of $MELLT1^{Cre}$, $METTL1^{CKO}$ and $METTL1^{CKO}$ + AAV9-$METTL1^{OE}$ mice after HFD + STZ for 12 weeks. (D-E) Systolic and diastolic blood pressure of $MELLT1^{Cre}$, $METTL1^{CKO}$ and $METTL1^{CKO}$ + AAV9-$METTL1^{OE}$ mice after HFD + STZ for 12 weeks. Data are expressed as mean ± SEM.

Figure S5

(A) Schematic diagram of tamoxifen induced myocardial METTL1 knock-out mice model. Mice were dived into three groups: METTL1$^{Cre}$ob/ob and METTL1$^{CKO}$ob/ob mice at age 16 weeks. (B) Representative UCG images of METTL1$^{Cre}$ob/ob and METTL1$^{CKO}$ob/ob mice. (C) Representative general photograph of heart tissues of METTL1$^{Cre}$ob/ob and METTL1$^{CKO}$ob/ob mice. Data are expressed as mean ± SEM. (D) HE, Masson and WGA staining of myocardial tissues among METTL1$^{Cre}$ob/ob and METTL1$^{CKO}$ob/ob mice. (E-F) Statistical analysis of fractional shortening and ejection fraction among METTL1$^{Cre}$ob/ob and METTL1$^{CKO}$ob/ob mice. (G) Serum levels of BNP in METTL1$^{Cre}$ob/ob and METTL1$^{CKO}$ob/ob mice at age 16 weeks. (H-I) Representative images and statistical analysis of ANP, BNP, Collagen I, α-SMA, NLRP3, TNF-α, ASC and IL-1β were detected by immunoblotting analysis in heart tissues from METTL1$^{Cre}$ob/ob and METTL1$^{CKO}$ob/ob mice. (J) Comparison of relative mRNA expression levels of genes associated with cardiac hypertrophy (Nppa, Myh7, Nppb and TINT2) among METTL1$^{Cre}$ob/ob and METTL1$^{CKO}$ob/ob mice. (K)Comparison of relative mRNA expression levels of genes associated with pro-inflammatory cytokines (IL-1β, IL-6, TNF-α and Cxcl2) among METTL1$^{Cre}$ob/ob and METTL1$^{CKO}$ob/ob mice. (L) Comparison of relative mRNA expression levels of genes associated with cardiac fibrosis (Col1α1, Acta2, TGF-β1 and Fibronectin) among METTL1$^{Cre}$ob/ob and METTL1$^{CKO}$ob/ob mice. Data are expressed as mean ± SEM.

Figure S6

(A) Differential proteins compared between METTL1$^{CKO}$ and METTL1$^{Cre}$ mice after proteomics analysis were shown by volcano plots. (B) Differential proteins compared between METTL1$^{CKO}$ and METTL1$^{Cre}$ mice after proteomics analysis were shown by heatmap. (C) GO analysis for the differential proteins compared between METTL1$^{CKO}$ and METTL1$^{Cre}$ mice after proteomics analysis. (D) KEGG analysis for the differential proteins compared between METTL1$^{CKO}$ and METTL1$^{Cre}$ mice after proteomics analysis. (E) Heatmap shown the expression levels of TCA Cycle associated proteins. (F) Heatmap shown the expression levels of cholesterol metabolism associated proteins. (G) Heatmap shown the expression levels of fatty acid biosynthesis associated proteins.

Figure S7

(A) Expression levels of lipid metabolism associated metabolites in heart tissues of $METTL1^{Cre}$ob/ob and $METTL1^{CKO}$ob/ob mice via metabonomics analysis. (B) Statistical analysis of metabolites of fatty β-oxidation pathway in heart tissues from $METTL1^{Cre}$ob/ob and $METTL1^{CKO}$ob/ob mice. Data are expressed as mean ± SEM. (C) GSEA analysis for differential proteins in proteomics analysis between $METTL1^{Cre}$ and $METTL1^{CKO}$ mice. (D) FAO Colorimetric Assay for NMCM cells from $METTL1^{Cre}$ and $METTL1^{CKO}$ mice *in vitro* induced via high glucose for 24h. (E) Seahorse analysis for OCR for NMCM cells from $METTL1^{Cre}$ and $METTL1^{CKO}$ mice *in vitro* induced via high glucose for 24h. (F) Statistical analysis of metabolites of fatty β-oxidation pathway in METTL1-OE stable AC16 cells *in vitro* induced via high glucose for 24h. (G) Heatmap expression profile of genes associated with lipid metabolism (GSE173384). (H) GO chordal diagram of differential genes of heart tissues (GSE173384).

Figure S8

(A-B) Analyse the coverage of sequencing reads from m7G-seq of heart tissue of $METTL1^{CKO}$ and $METTL1^{Cre}$ on genes annotated with peaks, and generate distribution plots (along with heatmaps) illustrating the read distribution across functional genomic regions associated with peak-containing genes. (C) Heatmap for the differential genes from m7G-seq. (D-E) Chromosomal distribution of peak counts. (F) Differential genes of m7G-seq and KEGG analysis of apelin signaling pathway (red marked differential genes).

Figure S9

(A) Flow chart of glycoproteomics analysis in cardiac tissue from ob/ob and ob/m mice at age 16 weeks. (B) The number of O-GlcNAcylation modification of sites, peptides and proteins in heart tissue from ob/ob and ob/m mice via glycoproteomics analysis. (C) Venn diagram shown the same O-GlcNAcylation modification of peptides and proteins in heart tissue from ob/ob and ob/m mice via glycoproteomics analysis. (D) The main GO enrichment analysed by enrichment pathway after glycoproteomics detection in heart tissue from ob/ob and ob/m mice.

Figure S10

(A) Distance matrix of amino acid residues between OGT and METTL1, illustrating the pairwise spatial relationships among all residues. (B) AlphaFold3-predicted structural model of the OGT-METTL1 complex. (C) Root Mean Square Deviation (RMSD) of the OGT-METTL1 complex over a 100 ns molecular dynamics simulation. (D) Root Mean Square Fluctuation (RMSF) and B-factor analysis of the OGT-METTL1 complex. (E) Electrostatic potential surface of the OGT-METTL1 complex.

Figure S11

(A) The number of hydrogen bonds formed in the OGT-METTL1 complex during the 100 ns molecular dynamics simulation. (B) Extract the molecular dynamics trajectory of the OGT-METTL1 complex from the last 10 ns (a total of 1000 frames) and calculate the binding free energy between OGT and METTL1 for each frame. (C) Decompose the binding free energy between OGT and METTL1 into the contributions of GGAS and GSOLV. (D) Decompose the binding free energy between OGT and METTL1 into the contributions of the interacting surface amino acids. (E) The contribution of interacting surface amino acids to the binding free energy between OGT and METTL1 for each frame.

Figure S12

(A) Flow chart of ubiquitylome analysis in cardiac tissue from ob/ob and ob/m mice at age 16 weeks. (B) Venn diagram shown the same ubiquitin modification of peptides and proteins in heart tissue from ob/ob and ob/m mice via ubiquitylome analysis. (C)

KEGG pathway analysed by enrichment pathway after ubiquitylome analysis in heart tissue from ob/ob and ob/m mice. (D) The different number of ubiquitin modification of proteins in heart tissue from ob/ob and ob/m mice via ubiquitylome analysis. (E) Subcellular structure localization of different proteins via ubiquitylome analysis. (F) The main GO enrichment analysed by enrichment pathway after ubiquitylome analysis in heart tissue from ob/ob and ob/m mice via Metascape database. (G) The main GO enrichment analysed by enrichment pathway after ubiquitylome analysis in heart tissue from ob/ob and ob/m mice via DAVID database.

Figure S13

(A) AC16 cells were cultured in HG for 24h treating with MG132 for 6h. Cells were collected for further detection the protein levels of METTL1. Data are expressed as mean ± SEM. (B) NMCM cells were cultured in HG for 24h treating with MG132 for 6h. Cells were collected for further detection the protein levels of METTL1. Data are expressed as mean ± SEM. (C) AC16 cells were cultured in HG for 24h treating with chloroquine (CQ) for 6h. Cells were collected for further detection the protein levels of METTL1. Data are expressed as mean ± SEM. (D) NMCM cells were cultured in HG for 24h treating with chloroquine (CQ) for 6h. Cells were collected for further detection the protein levels of METTL1. Data are expressed as mean ± SEM. (E) NMCM cells were treated with HG for 24h and stimulated with cycloheximide (CHX) for different time points. Then cells were collected for further immunoblotting analysis to detect the half-life time of METTL1.

Figure S14

(A) GO analysis for the different proteins associated with metabolism pathway identified via ubiquitylome analysis in cardiac tissue from ob/ob and ob/m mice at age 16 weeks. (B) Represent different proteins associated with hypertrophic cardiomyopathy, diabetic cardiomyopathy, fatty acid metabolism and ubiquitin mediated proteolysis via ubiquitylome analysis in cardiac tissue from ob/ob and ob/m mice at age 16 weeks. (C) GO analysis for the different proteins associated with metabolism pathway identified via glycoproteomics analysis in cardiac tissue from ob/ob and ob/m mice at age 16 weeks. (D) Represent different proteins associated with lipid modification, fatty acid oxidation, lipid particle and fatty acid transport via glycoproteomics analysis in cardiac tissue from ob/ob and ob/m mice at age 16 weeks. (E-G) GSEA analysis for the different proteins via ubiquitylome analysis and glycoproteomics analysis. (H) Different proteins associated with lipid metabolism proteins and its O-GlcNAcylation sites via glycoproteomics analysis in cardiac tissue from ob/ob and ob/m mice at age 16 weeks. (I) Different proteins associated with lipid metabolism proteins and its ubiquitin sites via ubiquitylome analysis in cardiac tissue from ob/ob and ob/m mice at age 16 weeks.

Figure S15

(A) AC16 cells were transfected with METTL1 (WT) and METTL1 (T268A) plasmids and cells were treated with HG for 24h. Cells were then collected for further WB assay to detect the expression levels of ANP, BNP, NLRP3, Collagen I, IL-6 and TNF-α. (B) NMCM cells were transfected with METTL1 (WT) and METTL1 (T262A) plasmids and cells were treated with HG for 24h. Cells were then collected for further WB assay to detect the expression levels of ANP, BNP, NLRP3, Collagen I, IL-6 and α-SMA.

Figure S16

(A-B) HEK293T cells were transfected with human DUB siRNA library and collected for ELISA and WB assay to detected the protein levels of METTL1. (C) Heatmap shown the ELISA assay results of METTL1 in HEK293T cells transfected with human DUB siRNA library. (D) AC16 cells were transfected with different DUBs plasmids and treated with HG for 24h. Cells were then collected for further WB assay to detect the expression level of METTL1.

Figure S17

(A) Mouse heart cell line (HL-1) cells were treated with HG for 24h and cells were collected for further Co-IP assay to detect the interaction of METTL1 and USP5. (B) AC16 cells were treated with HG for 24h and cells were collected for further Co-IP assay to detect the interaction of METTL1 and USP5. (C) NMCM cells were treated with HG for 24h and cells were collected for further Co-IP assay to detect the interaction of METTL1 and USP5. (D) NMCM cells were transfected with His Tag USP5 plasmid then cultured and treated with HG and Thiamet (5μM) for 24h, cells were collected for further Co-IP assay to detect the interaction level of METTL1and USP5.

Figure S18

(A) The Kendall analysis for the correlation of USP5 and METTL1 in heart of human. Data were download from the *GEPIA* Database. (A) The spearman analysis for the correlation of USP5 and METTL1 in aorta, lung, liver, muscle, kidney and whole blood of human. Data were download from the *GEPIA* Database. (B) The spearman analysis for the correlation of USP5 and METTL1 in DCM patient hearts via single-nuclei profiling. Data were download from the *GEPIA* Database.

Figure S19

(A-E) Left: the structure of Top1 to Top5 small molecule drugs. Middle: 3D schematic representation of the binding interaction between the top 1 to top5 ranked small molecule and OGT. Right: 2D schematic representation of the interaction between the top-ranked small molecule (Top1 to Top 5) and the OGT binding site.

Figure S20

(A) Description of constructing a luciferin reporting system for screening and detecting interaction levels of OGT and METTL1. (B) Flow chart of small molecule drug screening using luciferin reporting system of OGT and METTL1. (C) Bubble diagram shows the level of luciferin of OGT and METTL1 coupling (up). The relative levels of luciferin reporting system for screening and detecting interaction levels of OGT and METTL1 after treating cells with different small molecules (below).

Figure S21

(A) RMSD variation curve of the protein (blue) and the small molecule (red) during the 100ns molecular dynamics simulation. (B) Superimposed structures of 100 conformations saved every 1ns during the 100ns molecular dynamics simulation. (C) RMSF analysis plot of the protein during the 20-100ns molecular dynamics simulation (green-marked regions indicate amino acid residues interacting with the molecule. (D) Atom indexing diagram of the small molecule structure (up). RMSF analysis plot of the small molecule during the 20-100ns molecular dynamics simulation (below). (E) Analysis of key amino acids in the binding site to small molecule binding (up). MMGBSA binding free energy of the small molecule and protein over the last 10 ns (a total of 1000 frames) (below). (F) Time evolution of the total number of interactions between the small molecule and the protein during the molecular dynamics simulation (up). Time-dependent variations of critical amino acids mediating small molecule-protein interactions in molecular dynamics simulations (below).

Fig S22

(A-B) Statistical analysis and Serum level of ALT and AST from 24-week-old mice treated with DMSO (WT+DMSO) or HIT106265621 (WT+HIT106265621). (C-D) Statistical analysis and serum level of BUN and creatinine from 24-week-old mice treated with DMSO or HIT106265621. (E-G) Statistical analysis and serum level of white blood cells (WBC), hemoglobin (Hb) and hematocrit (HCT) from 24-week-old mice treated with DMSO or HIT106265621.
